# Whole-genome analyses point to new candidate genes underlying aestivation in amphibians

**DOI:** 10.1101/2025.07.24.666600

**Authors:** Leticia M. Ochoa-Ochoa, Rebecca D Tarvin

**Affiliations:** Universidad Nacional Autónoma de México: Universidad Nacional Autonoma de Mexico; Museum of Vertebrate Zoology and Department of Integrative Biology, University of California, Berkeley, Berkeley, CA 94720 USA

**Keywords:** dormancy, glycolysis, metabolism, oxidative phosphorylation, protein trafficking, TRAPP complex

## Abstract

When amphibians invaded the terrestrial realm, a strategy such as aestivation may have helped them to survive water and oxygen stress in the novel environment. In fact, aestivation appears to be an ancestral state in amphibians. We conducted a literature review about the genes previously linked to aestivation and then we searched for additional candidate genes related to the aestivation phenotype by comparing evolutionary rates of 13,578 genes in 31 amphibian species, including 10 aestivating species and 21 non-aestivating species. Based on the assumption that aestivation has two main requirements – a) metabolic regulation needed to control transitions to/from dormancy, and b) cell preservation strategies needed to sustain biological processes over long-term dormancy – we expected to find distinct relative evolutionary rates (RER) in genes related to those traits. We identified 323 genes with accelerated or decelerated RERs; these genes were enriched for some processes overlapping with our predictions and with literature findings, such as different modes of ATP production. Other genes related to protein and membrane trafficking (TRAPP, SNARE, and Arp2/3 protein complexes) are newly associated with aestivation. RER patterns suggest that aestivation in amphibians relies on a set of highly conserved core processes and other auxiliary processes that have diversified across the phylogeny. Given the vast number and diversity of aestivating amphibian lineages, we predict that more in-depth molecular studies of amphibian aestivation will offer novel insight into hypometabolic processes that could inspire medical innovations to prevent organ atrophy, address problems with angiogenesis, and combat processes underlying cancer.

## Introduction

Amphibians are distributed widely on Earth, except in regions with exceptionally limited access to liquid water such as the Sahara, Arabian, and Gobi deserts, or Antarctica (IUCN 2024, update 2023-1). These amazing vertebrates need humidity to survive, particularly because the vast majority carry out some gas exchange through their moist skin. Nevertheless, they have invaded and diversified in areas that might seem extreme, such as Central Australia or the Mojave and Sonoran deserts (Mayhew 1968). Amphibians living in seasonally arid environments avoid desiccation by producing lipids that waterproof their skin (e.g., Amey and Grigg 1995; Jørgensen 1997) or avoiding the hottest and driest season by resorting to aestivation, a physiological state in which they lower their metabolism and stay dormant for a lengthy period of time (weeks, months or even years), after which, when conditions are suitable, they raise their metabolism to continue daily activities (Navas and Carvalho 2010).

Aestivation is a preemptive state wherein animals prepare for water loss and diminished metabolic fuel supply. To avoid water loss, most amphibians seek refuge by burrowing. This does not mean that they resort to fossoriality because they are not active under the ground; rather they enter a state of depressed metabolism (i.e., hypometabolism, also known as dormancy, Storey 2000). Some amphibians while buried also produce cocoons made of multiple layers of shed skin, increasing the thickness of the cocoon according to the dryness of the environment (Booth 2006; Cartledge et al. 2006). The waterproof properties of the cocoon have been associated with the production of certain hydrophobic proteins such as ceramides and cerebrosides (Sadowski-Fugitt et al. 2012). Water loss is also retarded by colligative means, this implies elevating the osmolality of body fluids via the production of high concentrations of solutes like urea (a waste product of protein catabolism), in species that do not produce cocoons (Storey 2002; Luu et al. 2021). For example, spadefoot toads (*Scaphiopus couchii*) do not produce cocoons, but urea can rise up to 300 mM in their blood after several months of estivation (McClanahan 1972). Moreover, it has been shown in various species that water level does not change across vital organs (except intestine) when aestivating compared to non-aestivating amphibians (Balinsky et al. 1967; Bayomy et al. 2002; Cramp et al. 2005; Biggar et al. 2015a; Smith and Secor 2016; Anderson et al. 2017).

Aestivation must be prefaced by the accumulation of large reserves of endogenous fuels to deal with starvation during the dormant period (Storey 2002), followed by a reduction in energy use during dormancy through a lower metabolic rate. A typical metabolic rate during estivation in amphibians is approximately 10–30% less than the corresponding resting metabolic rate in active individuals at the same temperature (Pinder et al. 1992). There are some extreme cases where, for example, *S. couchii* may reduce their resting metabolic rate by 50% (Seymour 1973) or *Neobatrachus kunapalari* that can reduce it by up to 67% (Flanigan and Guppy 1997). The greater the reduction in metabolic rate, the longer the time that a fixed reserve of fuel can sustain basal metabolism, and the lower the amount of reactive oxygen species produced. It might be the reason why some amphibians seek hypoxic environments to accelerate entry into or deepen the hypometabolic state (Rossi et al. 2020). During aestivation, food ingestion ceases and the digestive tract is substantially reduced in size (Akat et al. 2014; Cramp et al. 2005, 2009). Energy metabolism during aestivation is based primarily on the anaerobic oxidation of lipids, although there seems to be some protein catabolism and a small direct contribution from carbohydrates. For example, in *Spea multiplicata*, the contributions to the total energy while aestivating were calculated as 62% from lipids, 32% from protein, and 5% from carbohydrate (Jones 1980). Protein synthesis, one of the most energy-expensive functions of cells, is strongly suppressed in estivating animals (Storey and Storey 2004, 2010). For example, in liver slices from aestivating frogs in Australia, *Neobatrachus sudellae*, protein synthesis was reduced by 67% compared with the rate in slices from non-aestivating frogs (Fuery et al. 1998). Furthermore, mechanisms of metabolic suppression in aestivators must be (a) rapidly reversible, and (b) require very little *de novo* protein synthesis and reorganization of metabolism, in order to both preserve energy and be prepared to enter into an active state as quickly as possible (Storey 2002; Storey and Storey 2012). Here we focus on the molecular aspects of aestivation but general reviews on aestivation are elsewhere (Navas and Carvalho 2010; Storey and Storey 2012; Wilsterman et al. 2021; Malik et al. 2023, just to name a few).

Aestivation may be an ancestral state in amphibians that has subsequently been lost many times in lineages where, presumably, aestivation is not necessary for survival (Ochoa-Ochoa et al. 2025). Here we take advantage of newly available annotated reference genomes to compare evolutionary rates of more than 13,000 genes between 10 aestivating species and 21 non-aestivating species. In this study we aim to: 1) identify candidate genes and biological processes that underlie estivation in amphibians, 2) evaluate whether these processes are highly conserved or derived in aestivating amphibians, and 3) review whether the genes we identify have been previously studied in amphibians or are newly identified. We employ an untargeted approach, which has the advantage of discovering potentially novel genes and pathways involved in this amazing adaptation.

## Methods

### Literature review

We followed a process similar to Ochoa-Ochoa et al. (2025) that consisted of performing a systematic search using the advanced search tool of each of the following search engines: Scopus, PubMed, and Web of Science, until December 2024. We used the words (aestivat* AND amphibian*) and (estivat* AND amphibian*). Studies were selected according to the following criteria: 1) empirical papers and reviews including theses; 2) empirical articles in which some biological, ecological, physiological, or genetic aspect of the aestivation process were mentioned; and 3) papers where the focus was not related to dormancy processes, but an aestivating species is mentioned. Afterwards we selected all the papers that contained the word “gene”. Each of those published manuscripts was manually reviewed for a gene or protein involved in the aestivation process. Scientific names of amphibian species follow AmphibiaWeb (2024) nomenclature.

We recorded the tissue analyzed in each study (brain, heart, kidney, liver, muscle, lung, gut, and skin) because the same gene, miRNA, or metabolite can have increased activity in one tissue and decreased activity in another. In some manuscripts, skin and muscle were divided (e.g., dorsal and ventral skin), but here we considered these data together, without intra divisions. We also registered whether the activity was significantly increased or decreased in each tissue, and in the case of a non-significant change it was recorded as maintained. Thus, a single gene can be in both increased and decreased datasets. Due to the fact that some manuscripts did not report gene symbols, we used either the name of the gene provided or its description to search for gene symbols based on the *Xenopus tropicalis* ensembl dataset (Dyer et al. 2025).

### Obtaining sequence data

We initially attempted to use Orthofinder v2.5.5 (Emms and Kelly 2019) to identify orthologs across amphibian genomes, but the pipeline using primary transcripts and default settings identified only 161 single-copy orthologous genes. Reviewing the estimated gene duplications output from Orthofinder, three species were estimated to have a large number of duplicated genes: *X. laevis* (19,899), *Pelobates cultripes* (26,583) and *Staurois parvus* (66,924). Although it makes sense that *X. laevis* has so many duplications, as it is a tetraploid (Hughes and Hughes 1993), the other two species are thought to be diploid (Liedtke et al. 2022; Uno 2021). Both of these genomes were annotated by third parties. Although *Hymenochirus boettgeri* was also annotated by a third party, it did not show an inflated number of estimated gene duplications (8,452). After removing *X. laevis*, *P. cultripes,* and *S. parvus* from the Orthofinder analysis, we were still only able to identify 1,845 single-copy orthologs. Thus, we chose to develop a different pipeline based on NCBI gene annotations (see Supplementary Material Appendix A1, available online). We note that there are drawbacks to both of these methods, and neither is perfect. However, we believe that using the pipeline described below is adequate for the purpose of this study.

To get orthologs for amphibian genomes, we first downloaded the full gene set from *X. tropicalis* using the “datasets download gene taxon” command from the NCBI Datasets command line tools (O’Leary et al. 2024). Then we used a custom script (see Appendix A1, available online) to obtain and filter the metadata for gene names and symbols. The list of gene symbols was used to download all orthologs in amphibians using the “datasets download gene symbol” tool in NCBI Datasets. In some cases, multiple isoforms were available for an ortholog; thus, we selected the longest isoform for downstream analyses using a custom script. Once we had all ortholog sequences, we estimated sequence alignments for each gene using MAFFT v7.526 (Katoh and Standley 2013) and gene trees using IQ-TREE v2.4.0 (Minh et al. 2020). Because downstream RER analyses require that the trees have the same topology, we used a tree from Portik et al. (2023) pruned to the species of interest to constrain the tree inference (option - g in IQ-TREE). *Microcaecilia unicolor* was not present in that tree, so we added the tip manually into the newick file. To adjust branching times for the caecilian clade including *M. unicolor*, we integrated estimates from TimeTree.org (Kumar et al. 2022): age of *M. unicolor* + *G. seraphini* = 120.1 MY; age of *M. unicolor* + *G. seraphini* + *R. bivittatum* = 221.3 MY. Finally, we rerooted the trees to standardize the topology using the root and reroot commands in ape v5.8-1 (Paradis and Schliep 2019) and phytools v2.4-4 (Revell 2024). All of the commands to obtain, align, and infer phylogenies of orthologs can be found in Appendix A1 (available online).

### Estimation of evolutionary rates

We estimated the relative evolutionary rates of 13,578 genes based on *X. tropicali*s orthologs. In order to do so we used RERconverge (Chikina et al. 2025) in R v. 4.4.2 (R Core Team 2024) to identify genes with consistent and significant departures from average rates of evolution across a phylogeny. Here we compare evolutionary rates in aestivating species to rates in putatively non-estivating species using binary trait data (species that aestivate = 1, species that do not aestivate or do not have information = 0). To categorize aestivation ability for each species, we used the binary classification based on data available in the literature, reviewed in Ochoa-Ochoa et al. (2025). Of the 31 amphibian species with annotated genomes, ten are recognized to aestivate (Table 1, Fig. 1). We note that this may be an incomplete list because there is no data about aestivation for many species.

**Figure 1.**
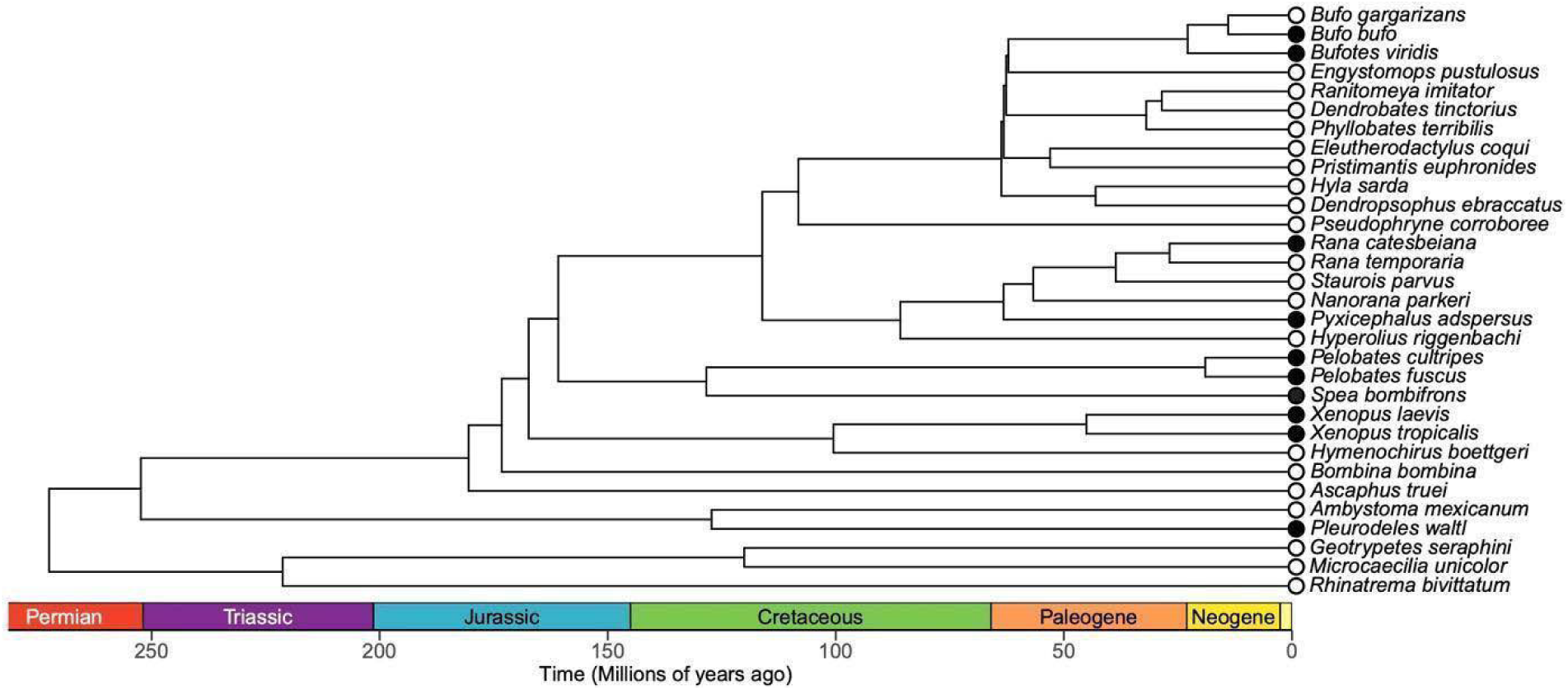
Phylogenetic tree of the amphibian species included in the analyses subsetted from Portik et al. (2023) and the imputed *Microcaecilia unicolor* (see methods). Black circles indicate that according to the literature review the species perform aestivation. White circles represent either lack of information or that in the literature has been reported that the species does not aestivate (i.e., *Ascaphus truei*).

**Table 1.**
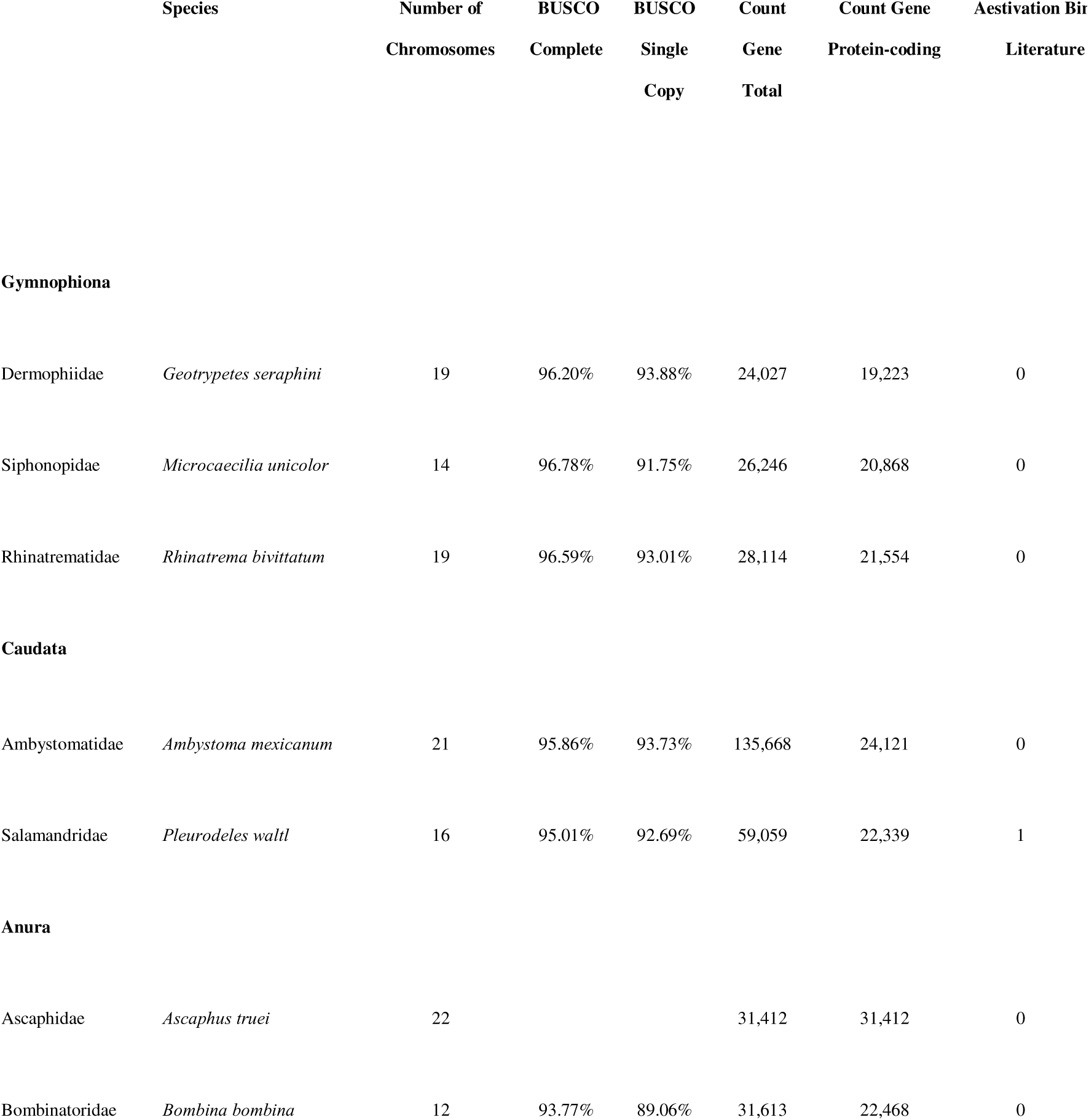
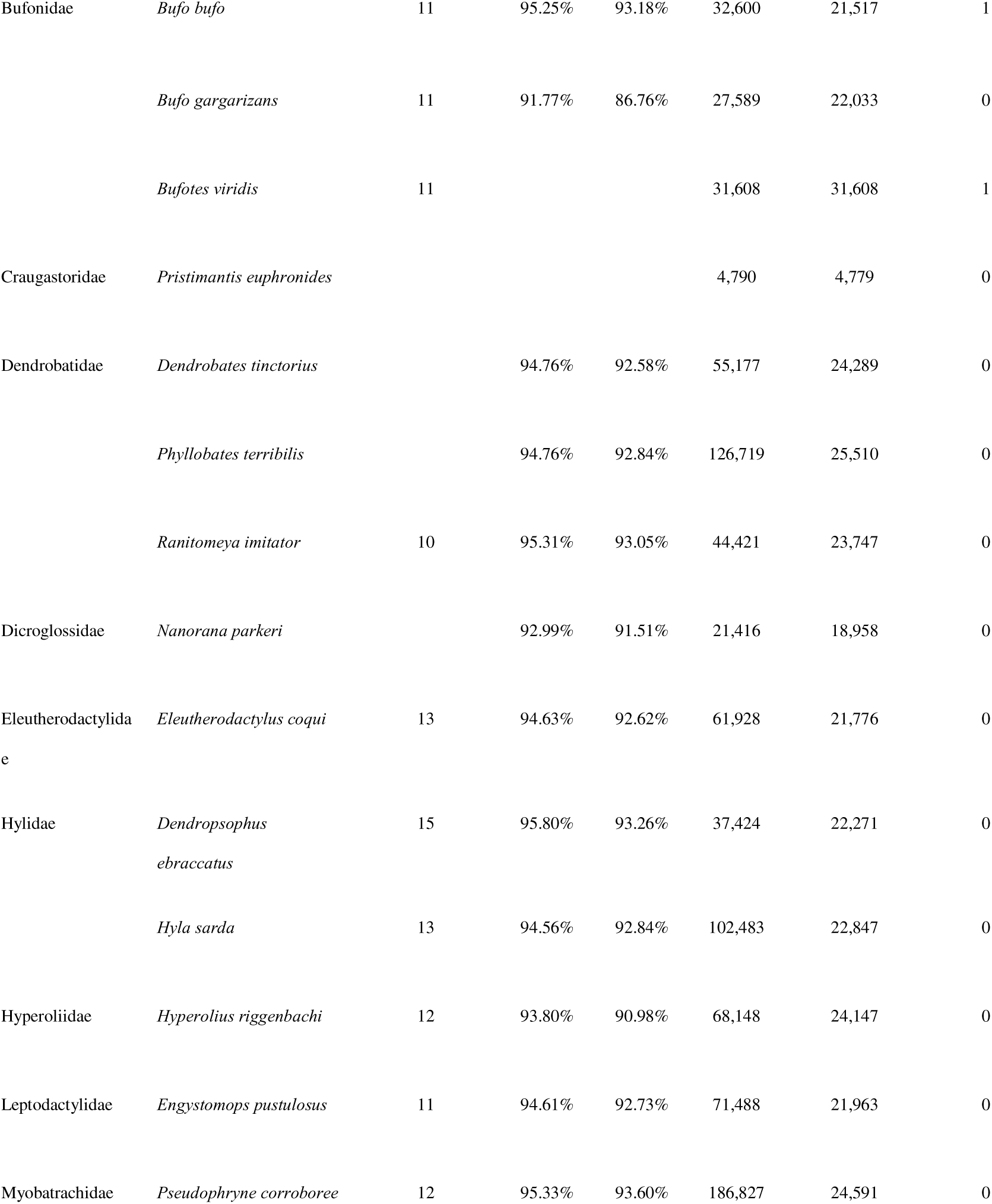
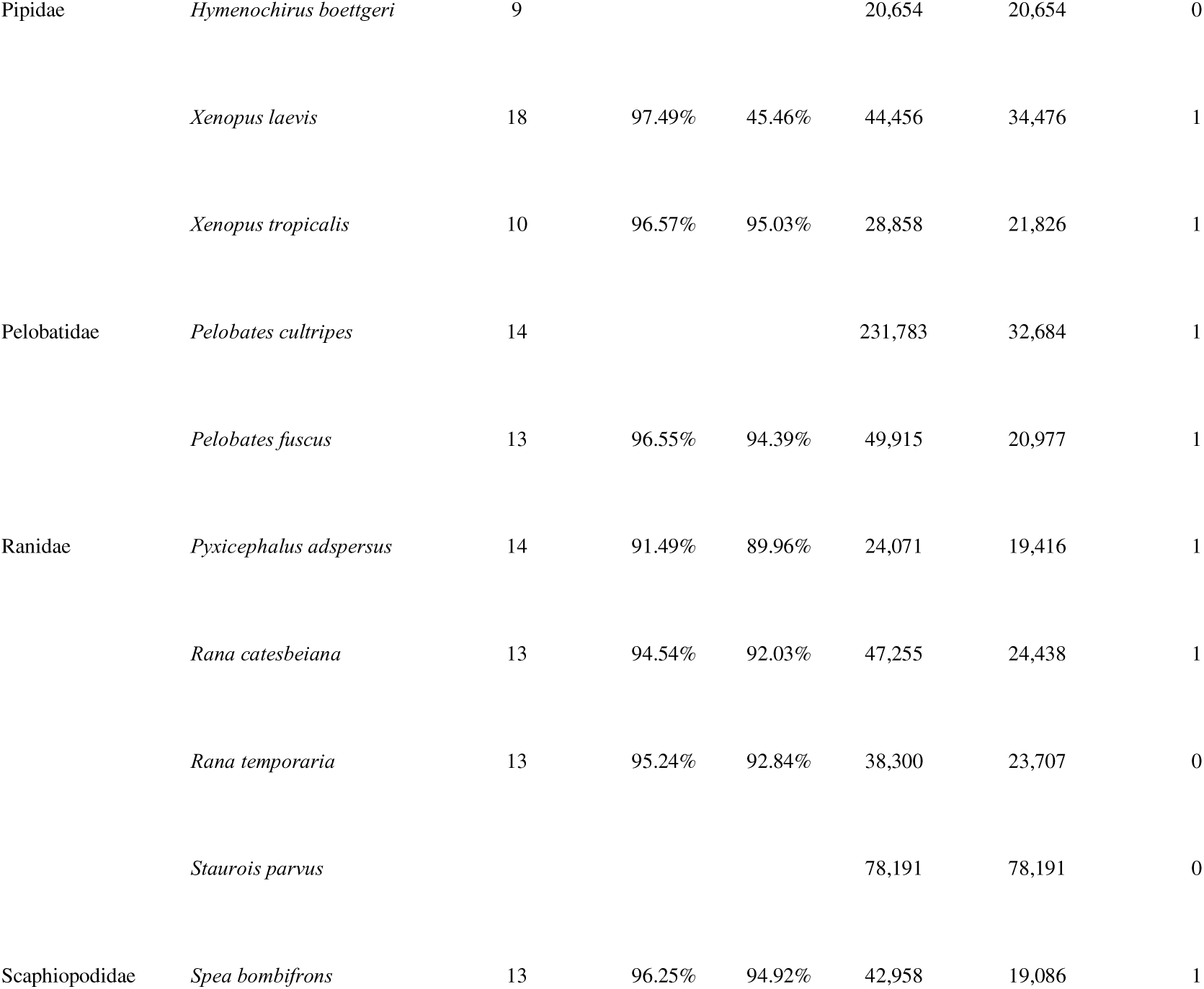
List of amphibian species used in the analyses. All species have annotated genomes and most species have chromosome-level assemblies. BUSCO analyses are based on tetrapoda odb10 and were accessed via reported values on NCBI (accessed May 25, 2025).

We followed the developers’ recommendations: no master tree provided and re-estimation of branch lengths when reading the trees; and for the function getAllResiduals we chose square root transformation, to use weighted regression for relative rates, and to scale relative rates internally for each species subset (see Supplementary Material Appendix A2, available online). We developed three hypotheses:

**H0** There is no set of genes with significant evolutionary rates associated specifically with aestivation.

**H1** Only genes with significantly higher evolutionary rates (i.e., positive selection) are associated with aestivation which would imply diversification of the aestivation phenotype.

**H2** Only genes with significantly lower evolutionary rates (i.e., purifying selection) are associated with aestivation which would imply conservation of the aestivation phenotype. **H3** Both genes with higher evolutionary rates and genes with low evolutionary rates are associated with aestivation, which would imply conservation of some aspects and diversification of others.

Finally, we compared gene symbols between significant RER genes and genes previously studied in our literature review to identify genes present in both datasets. We note that RER provides information on coding sequence evolution while most prior studies of aestivation have focused on gene expression, so we expected some overlap but note that the analyses provide different types of data.

### Gene ontology analysis (GO analyses)

We performed a GO analysis on the genes identified by RER to be evolving at distinct evolutionary rates in aestivating species. We used as background the set of genes that were estimated to have a Rho value, i.e., any gene with a positive or negative evolutionary rate regardless of the significance. In other words, we excluded the genes that have either a missing (NA) or 0 value for Rho. Thus, we had a background dataset containing 13,183 genes, of which 323 had distinctly accelerated (positive) or decelerated (negative) evolutionary rates; based on the gene sets with distinct RERs, we performed two GO analyses, one for positive and one for negative genes. It is important to specify that there is no overlap between the positive and negative datasets, since a gene has either a positive or negative Rho value.

We used the package topGo v2.58.0 (Alexa and Rahnenführer 2024) in R v4.4.2 (R Core Team 2024) to identify significantly enriched terms in our set of 165 / 158 (positive / negative) genes obtained from the RER analysis. For the gene2GO list of gene names and GoTerm identifiers, we used the annotated list for *X. tropicalis* (The UniProt Consortium 2025, https://www.uniprot.org/; downloaded June 12, 2025) that contains 76,381 entries. We used a nodeSize of 10 and for the selected genes and terms we set the topNodes at 50 (full script and data can be found in the Supplementary Material Appendix A3, available online).

We used Revigo (Supek et al. 2011) to simplify the visualization of GO terms through network inference and Cytoscape (Shannon et al. 2003) to visualize the networks. Finally, we searched for the protein name and function of the selected genes (https://www.uniprot.org/; accessed June 18, 2025) and significant terms (https://www.ebi.ac.uk/QuickGO/; accessed June 18, 2025).

### STRING (Search Tool for the Retrieval of Interacting Genes/Proteins)

To further visualize results from RER, we input the positive and negative gene sets into the Search Tool for the Retrieval of Interacting Genes/Proteins (STRING) webtool (Szklarczyk et al. 2023). We estimated full networks (both functional and physical associations) using either *Homo sapiens* or *X. tropicalis* annotation information. We set the maximum number of 1st shell interactors to 10, the maximum number of 2nd shell interactors to 5, the confidence to high (0.700), and hid disconnected nodes from the results. Shell interactors are proteins that are not in the query dataset but are predicted to interact with query proteins; including some of these in cluster inference assists the program in identifying protein complexes that may be relevant to the query dataset. Then we inferred gene network clusters using MCL clustering and default settings (with inflation parameter = 3). Results from *H. sapiens* were qualitatively similar to those from *X. tropicalis* and included slightly more annotations, so we only present networks from *H. sapiens.* However, when clusters were annotated using *X. tropicalis* data but not *H. sapiens*, we include this information clearly marked in the network.

## Results

### Literature review

From the 291 manuscripts found by Ochoa-Ochoa et al. (2025), 251 contained the word gene, and of these, 39 contained information about a gene, metabolite, miRNA, or protein analyzed during aestivation or water restriction, i.e., dehydration (Table S1, Supplementary Material, available online). Some form of metabolic function or gene expression has been analyzed in 20 species, with *X. laevis* being the most common species, used in 15 studies, followed by *Cyclorana alboguttata* with 7 studies and *S. couchii* with 6. Studies have focused on different measurements of metabolic depression such as protein quantification (e.g., Bayomy et al. 2002; Luu et al. 2020, 2018; Madelaire et al. 2020; Malik and Storey 2009; Malik et al. 2023; Pakay et al. 2003; Wu et al. 2017; Zhang et al. 2018), enzyme or metabolite activity (e.g., Bayomy et al. 2002; Cartledge et al. 2006; Flanigan et al. 1990; Katzenback et al. 2014; Patnaik and Sahoo 2021), miRNA abundance (e.g., Biggar et al. 2022; Luu and Storey 2015; Naranjo et al. 2023), targeted RT-qPCR (e.g., Biggar et al. 2022; Luu and Storey 2015; Naranjo et al. 2023; Wu et al. 2013) and untargeted RNAseq (Reilly et al. 2013; Hudson et al. 2024 both with *C. alboguttata*).

1464 unique proteins, metabolites, and genes have been studied, as well as 74 miRNAs, across a total of 39 studies. Because they are not directly comparable to our results, which focus on genes that code for proteins, we exclude metabolites such as ATP, hormones or other compounds such as vasopressin, and miRNAs from further consideration (see full list in Table S2, Supplementary Material, available online). Of the total 1434 genes/proteins previously studied, 707 had an increased activity and 755 had a decreased activity in at least one tissue. Skeletal muscle was the tissue most frequently analyzed followed by liver and kidney (Table S1). We note that of the >2500 genes showing significant changes in expression by Hudson et al. (2024) in *C. alboguttata*, 739 sequences were not annotated (no BLAST results in their paper), and another 660 were not assigned a gene symbol through a search using the accession numbers provided; therefore, 1399 of their genes were not included in our assessment. Probably the most amazing aspect of aestivation that has been demonstrated is that although there is an overall metabolic depression, usually measured by O_2_ consumption, there is a wide range in responses among tissues from no changes generally in the brain (e.g., Chauvet et al. 1993; Flanigan et al. 1990; Flanigan and Guppy 1997; Borah et al. 2020; Luu and Storey 2015; Wu et al. 2017,2020; Cowan et al. 2000) to extreme changes in the skeletal muscle (33 of the 39 articles mentioned found a significant change in muscle, Table S1). Of course, there are exceptions, for example, in aestivating *S. couchii* toads the brain contained increased activities of fructose-1,6-bisphosphatase (FBPase), phosphoenolpyruvate carboxykinase (PEPCK) and malic enzyme (ME), which all have roles to play in gluconeogenesis; it also had an increased activity of b-Hydroxybutyrate dehydrogenase (BDH) suggesting the possible enhanced use of ketone bodies as an aerobic fuel for the brain (Cowan et al. 2000).

### Estimation of evolutionary rates

We obtained orthologs for 31 species (Fig. 1; Table 1), but for the analyses the maximum number of species in a single gene tree was 25. From this set of species only 10 have been recorded to aestivate according to the literature (*Bufo bufo, Bufotes viridis, Rana catesbeiana, Pyxicephalus adspersus, Pelobates cultripes, P. fuscus, Spea bombifrons, X. laevis, X. tropicalis,* and *Pleurodeles waltl*). RER results from 13,578 genes indicated 13,184 with Rho values, 394 without (when few sequences were available), 6,918 with an accelerated evolutionary rate (a positive Rho) and 6,265 with a negative Rho, thus indicating a decelerated evolutionary rate. In total, 165 had a significantly positive RER and 158 had a significantly negative RER, prior to multiple testing correction. When using the Benjamini-Hochberg correction (i.e., an estimate of the false discovery rate), no genes reached significance (full list can be found in Table S3 of the Supplementary Material, available online). Thus, rather than calling these RERs significant, we refer to them as distinct, or as the “accelerated/decelerated gene sets”. We also specifically highlight the top ten genes ranked by p-values from accelerated (Table 2) and decelerated (Table 3) RERs, which showed diverse functions that in some cases were represented by significant terms in the GO analyses or were present in protein complexes inferred by STRING. We want to note that a lack of significance after multiple testing correction as occurred here is fairly common in studies that use RER (Chikina et al. 2016), including in the tutorial outlined in the R package documentation (Kowalczyk et al. 2019; Meyer 2018). It is likely related to the large number of genes involved in these studies.

**Table 2.**
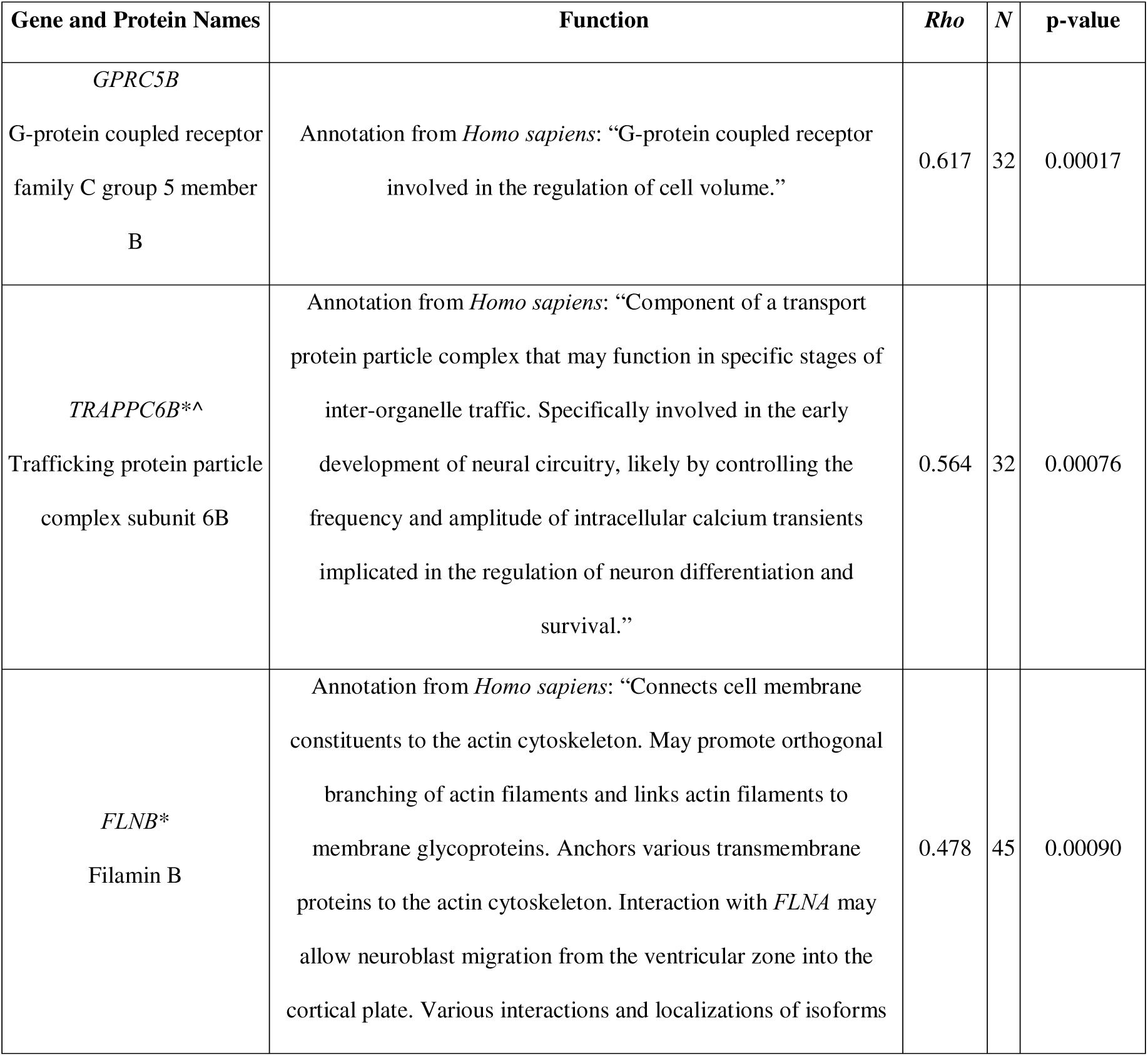
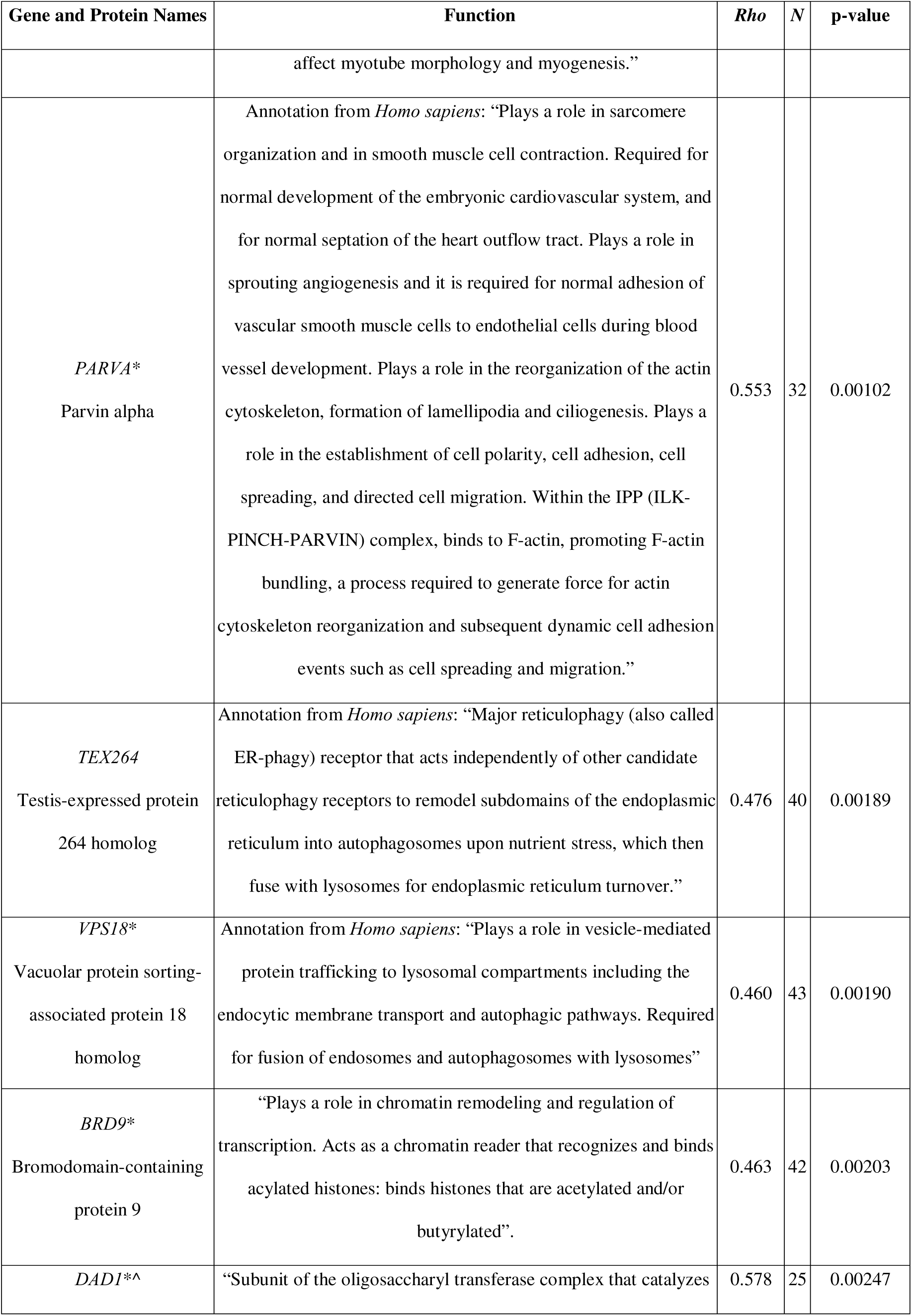
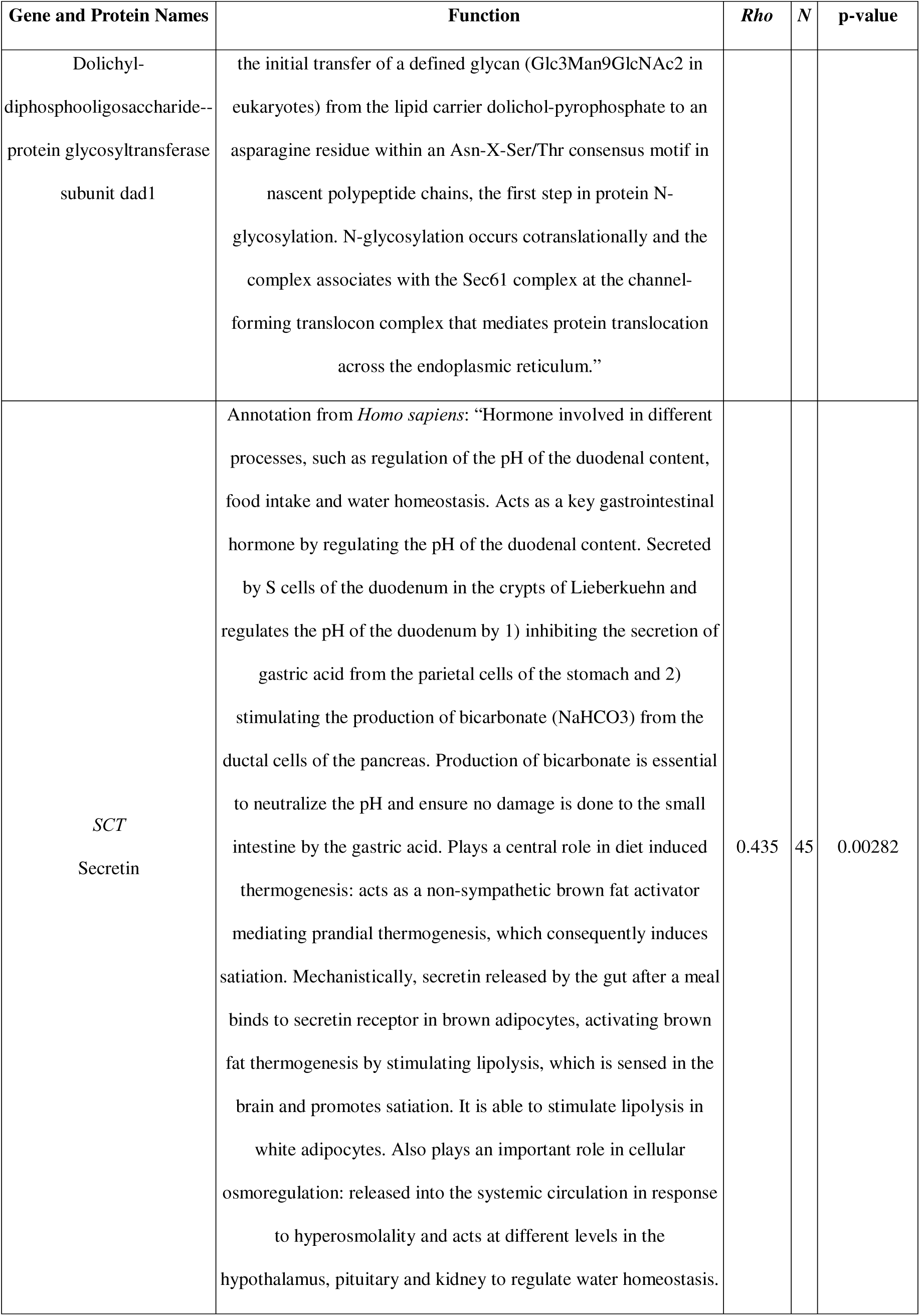
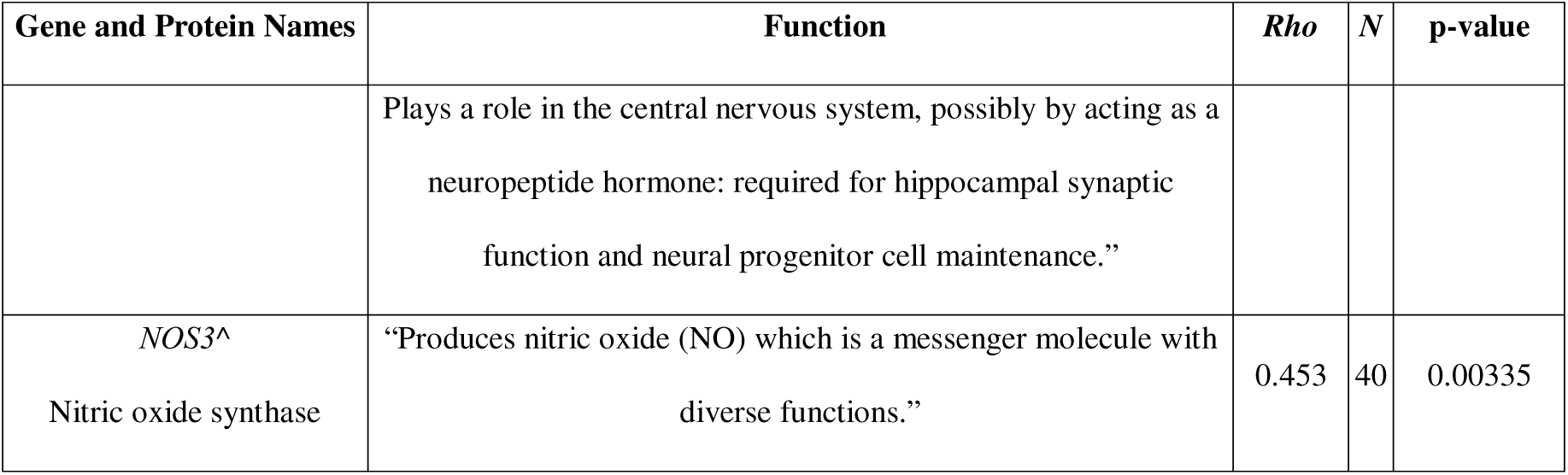
The top 10 genes identified by RER as under relatively accelerated evolutionary rates (p ≤ 0.05). Note: none of these genes were significant after multiple testing corrections. Protein names and functions were copied word-for-word from the UniProt database under a CC BY 4.0 license on 6/9/25 for entries of *Xenopus laevis* or *Xenopus tropicalis* unless otherwise noted (The UniProt Consortium 2025). Text was copied under a CC BY 4.0 license and PubMed citations were excluded for readability. *Denotes genes that were also associated with significantly enriched GO terms. ^Denotes genes that were present in a STRING gene cluster.

**Table 3.**
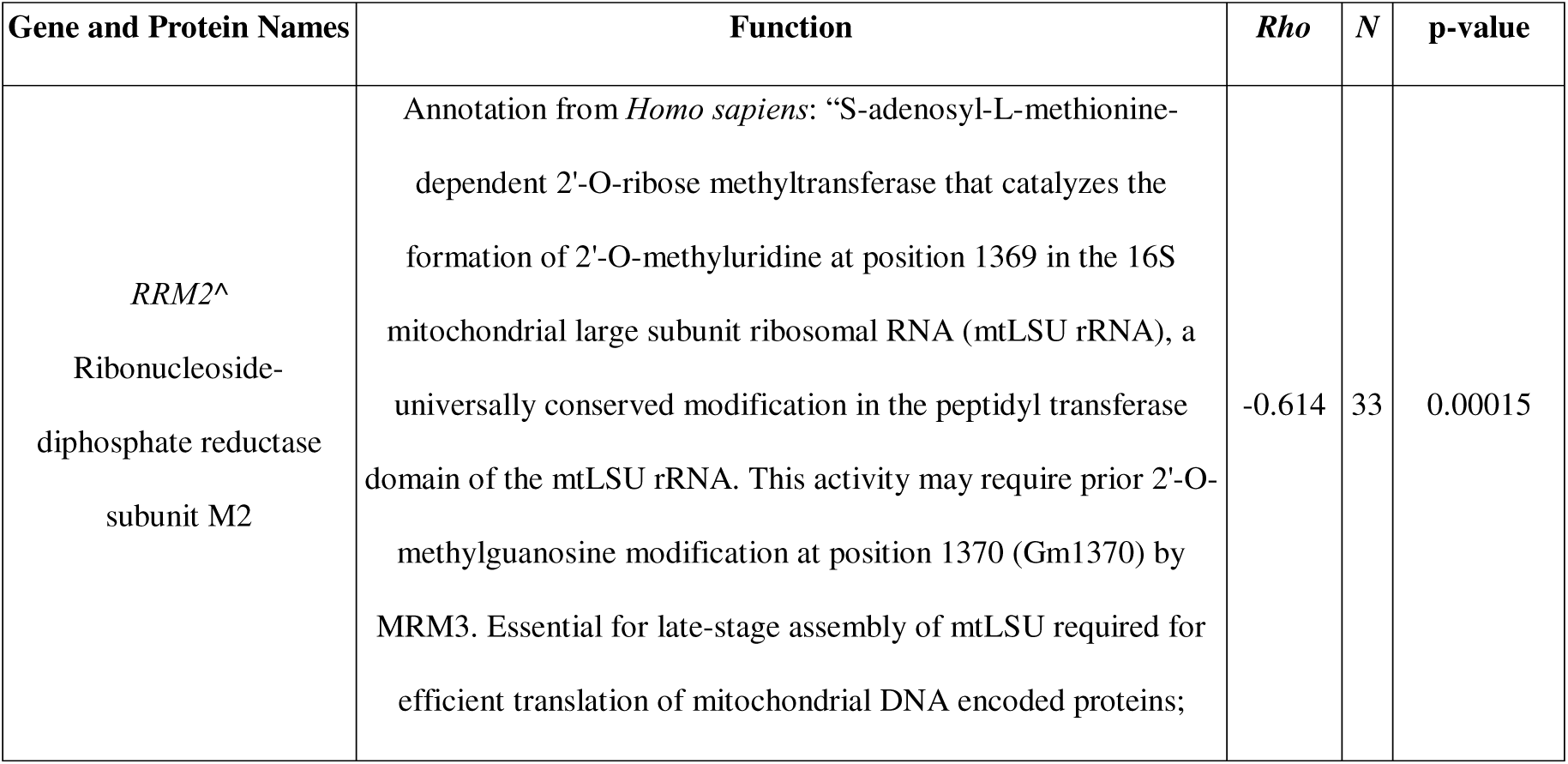
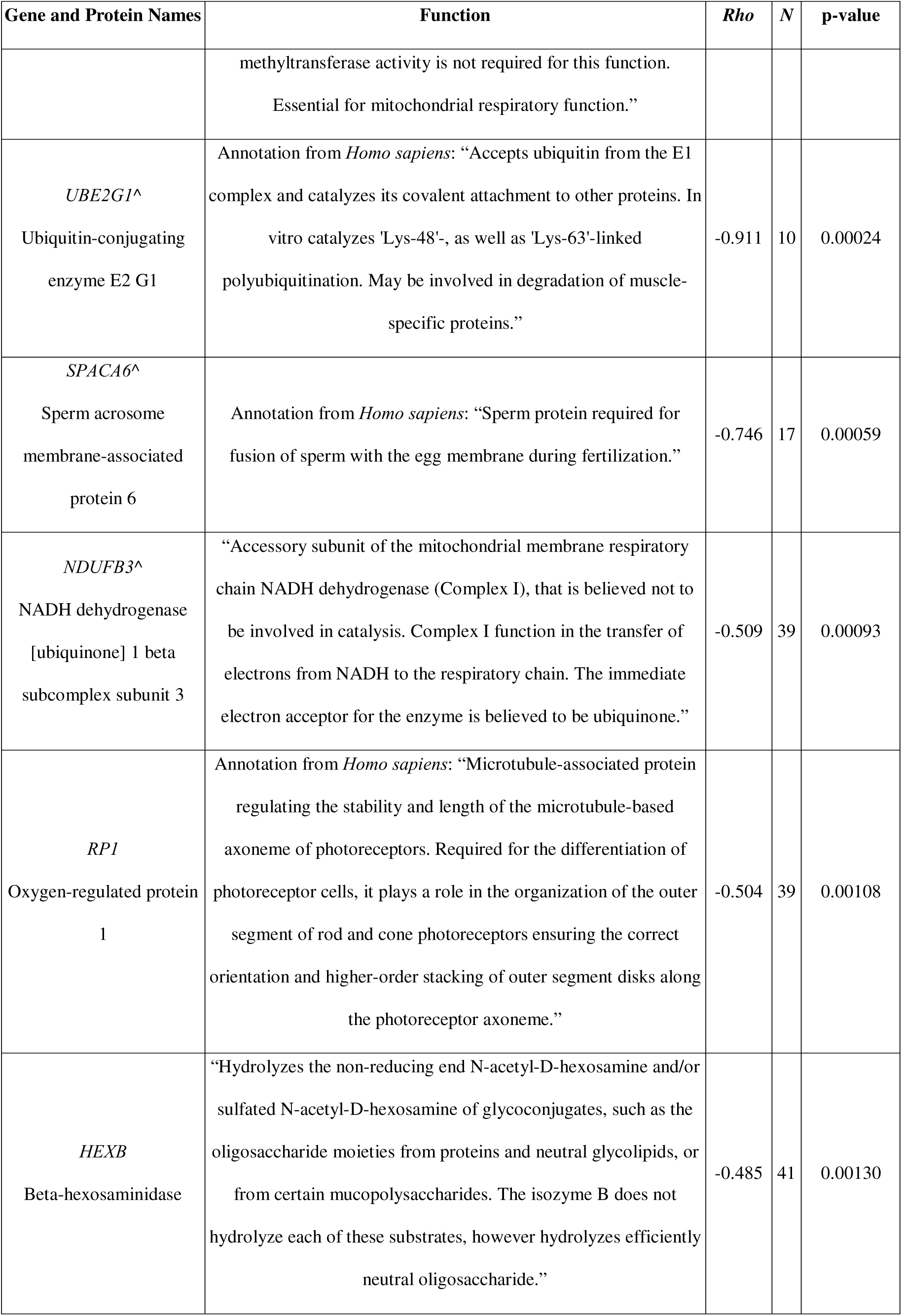
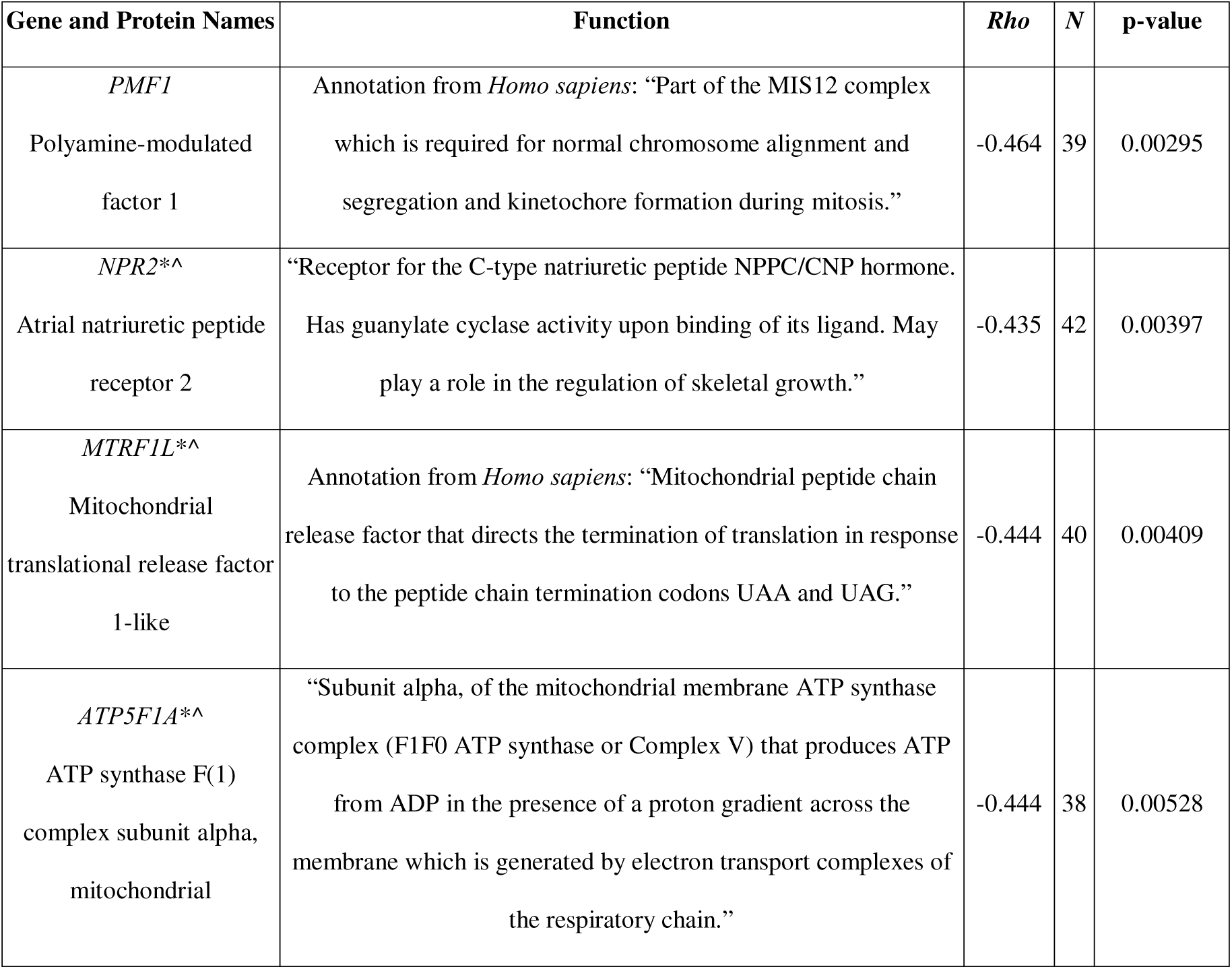
The top 10 genes identified by RER as under relatively decelerated evolutionary rates (p ≤ 0.05). Note: none of these genes were significant in RER after multiple testing corrections. Protein names and functions were word-for-word from the UniProt database under a CC BY 4.0 license on 6/9/25 for entries of *Xenopus laevis* or *Xenopus tropicalis* unless otherwise noted (The UniProt Consortium 2025). Text was copied under a CC BY 4.0 license and PubMed citations were excluded for readability. *Denotes genes that were also associated with significantly enriched GO terms. ^Denotes genes that were present in a STRING gene cluster.

We found 1215 genes that overlapped between the 13,184 *X. tropicali*s orthologs and the 1434 genes identified in the literature review (Table S1). Of these 1215 genes, 30 also showed a distinctly accelerated (11) or decelerated (19) RER. Of these 30, 9 showed increased activity in at least one tissue, and 18 showed decreased activity in at least one tissue; lastly 3 showed increased activity in one tissue and decreased in another (*GPI*, *JCHAIN*, and *PKM*; Table 4).

**Table 4.**
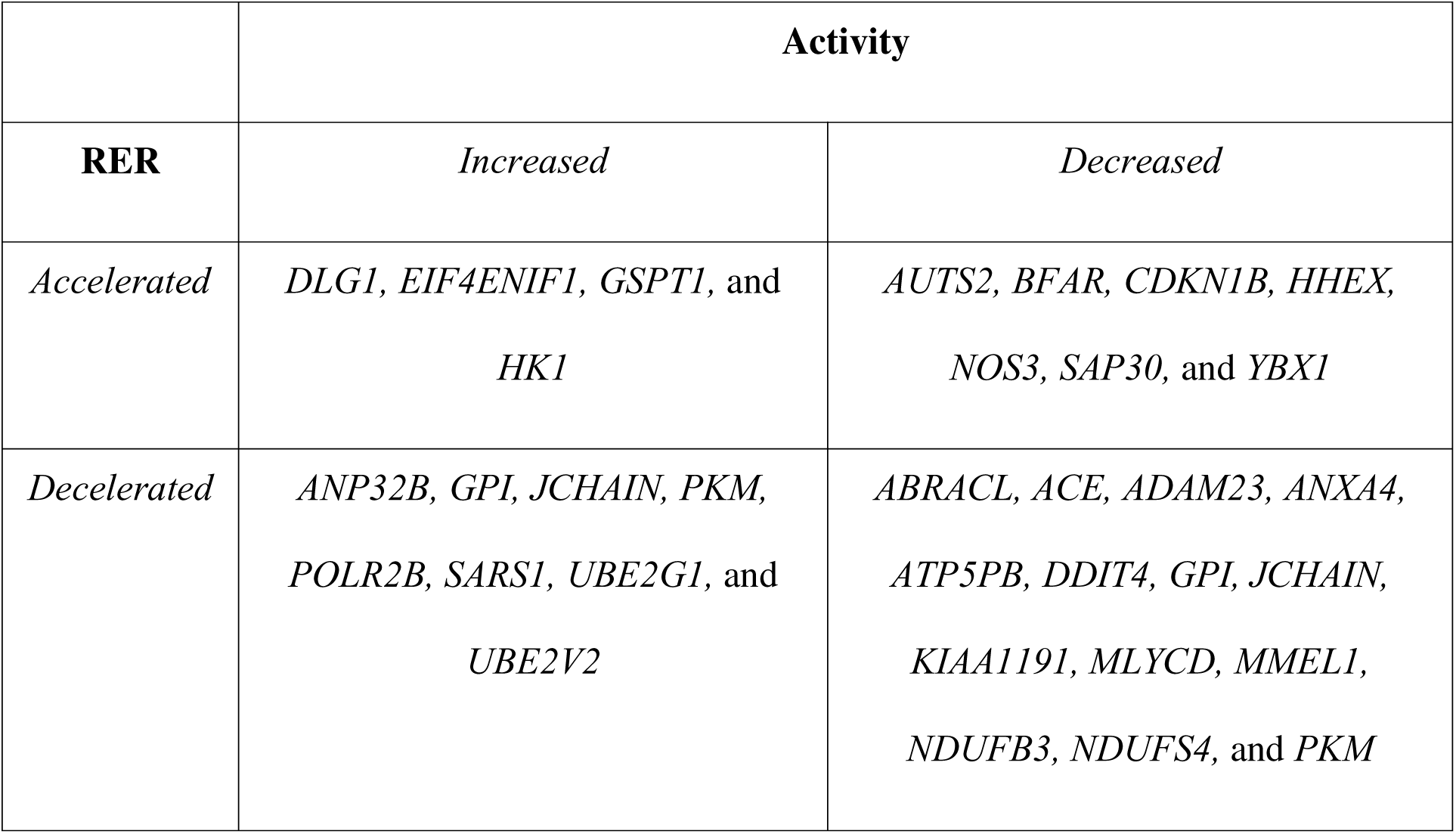
Genes with distinct relative evolutionary rates (RERs) that were also shown to have changes in activity in aestivating amphibians in prior studies that were identified in our literature review. Full descriptions of each gene can be found in Appendix A4 available online.

|  | **Activity** |  |
| --- | --- | --- |
| **RER** | *Increased* | *Decreased* |
| *Accelerated* | *DLG1, EIF4ENIF1, GSPT1, and*<br>*HK1* | *AUTS2, BFAR, CDKN1B, HHEX,*<br>*NOS3, SAP30, and YBX1* |
| *Decelerated* | *ANP32B, GPI, JCHAIN, PKM,*<br>*POLR2B, SARS1, UBE2G1, and*<br>*UBE2V2* | *ABRACL, ACE, ADAM23, ANXA4,*<br>*ATP5PB, DDIT4, GPI, JCHAIN,*<br>*KIAA1191, MLYCD, MMEL1,*<br>*NDUFB3, NDUFS4, and PKM* |

### Gene ontology analysis

We identified a total of 148 significantly enriched terms when comparing genes identified with a distinct RER to all genes in the background dataset (p < 0.05). 80 terms were significantly enriched in the distinctly accelerated gene set compared to the background gene set (p < 0.05). For the distinctly decelerated gene set, 68 terms were significantly enriched (Table S4, Supplementary Material, available online). Many genes were annotated with multiple terms (Fig. S1, in appendix A4, of the Supplementary Material, available online). Specifically, the terms correspond to 73 genes for distinctly accelerated, and 43 genes to decelerated gene sets, annotated to different GO term categories: 23 Molecular Function (MF), 71 Biological Process (BP) and 18 Cellular Compartment (CC) for accelerated and 26 MF, 18 BP and 17 CC for decelerated, respectively. Revigo summarized topGo results by collapsing similar terms into 64 parental terms, 22 for accelerated RER and 42 for decelerated RER.

MF showed 21 parental terms, 9 accelerated and 13 decelerated (Fig. S2, in appendix A4, available online). Accelerated terms, such as “protein-containing complex binding”, “phospholipid binding”, “lipid binding” and “NADP binding” are involved in intracellular trafficking, transport between the trans-Golgi network, and endosomes. This is a pathway that promotes the recycling or degradation of internalized transmembrane proteins, it may also involve macropinocytosis which is a form of endocytosis that results in the internalization of extracellular content (Kay 2021). There are also various terms, such as “phosphatidylinositol phospholipase C activity” and “protein kinase inhibitor activity”, that are involved in the transcription machinery particularly as regulatory and inhibitory processes. The decelerated terms, like “translation factor activity, “RNA binding”, “[DNA-directed 5’-3’] RNA polymerase activity”, and “catalytic activity, acting on a tRNA / RNA” are involved in transcription, ribosome and structural molecule activity related to protein synthesis and degradation of different polypeptides by substrate binding (e.g., “structural constituent of ribosome” and “structural molecule activity”).

BP had the largest number of parental terms after collapsing, 22 accelerated and 13 decelerated (Fig. S3, in appendix A4, available online). It is probably the most complex of all ontologies. Accelerated terms, such as “monosaccharide metabolic process”, “transcription by RNA polymerase III”, “nucleoside phosphate metabolic process”, “phosphorus metabolic process”, “glycosylation”, “pyridine-containing compound metabolic process”, “macroautophagy”, “carbohydrate derivative metabolic process”, and “carbohydrate derivative biosynthetic process”, are involved in the pentose phosphate pathway (PPP) for carbohydrate degradation, an alternative to glycolysis and a major source of reducing power and metabolic intermediates for biosynthetic processes; others such as “vesicle-mediated transport”, “macromolecule localization”, “intracellular transport”, “Golgi vesicle transport” and “cellular localization”, are involved in intracellular and endosomal membrane trafficking and in transport between endoplasmic reticulum and the trans-Golgi network (TGN). The term “cellular process” was the most complex and presented the highest number of genes involved, 70 in total. The decelerated terms, like “purine ribonucleotide metabolic process”, “cyclic nucleotide metabolic process”, “cyclic nucleotide biosynthetic process”, “ribose phosphate metabolic process”, “translation”, “amino acid activation”, and “tRNA aminoacylation for protein translation”, are involved in translation, in the formation of cAMP in response to calcium entry leading to cAMP signaling activation, and in glycolysis.

CC had the smallest number of parental terms after collapsing, 3 accelerated and 3 decelerated (Fig. S4, in appendix A4, available online). The accelerated terms (“endoplasmic reticulum protein-containing complex”, “organelle subcompartment” and “perinuclear region of cytoplasm”) are involved in the retrograde transport from endosome to plasma membrane, a trafficking pathway that promotes the recycling of internalized transmembrane proteins. It also mediates delivery of transmembrane proteins into the lumen of the lysosome for degradation. The decelerated terms, like “mitochondrial inner membrane”, “mitochondrion”, “ribosome” and “perinuclear region of cytoplasm”, are involved with the protein synthesis and the ribosome which is a large ribonucleoprotein complex responsible for the synthesis of proteins in the cell.

### STRING (Search Tool for the Retrieval of Interacting Genes/Proteins) Results

STRING MCL clustering inferred 17 clusters for the accelerated gene set and 20 clusters for the decelerated gene set (full list can be found in appendix A5 of the Supplementary Material, available online). The largest cluster in the accelerated gene set (Fig. 2) was the TRAPP complex (10 genes, although 8 are inferred interactors), followed by an unnamed cluster with 6 genes (1 inferred interactor) possibly related to translation deduced by the involved genes (*SRPRA, RPL37, RPS26, GSPT1, BTF3, TMCO1*), a cluster of 4 genes (1 inferred interactor) related to glucose 6-phosphate metabolism, an unnamed cluster of 3 genes related to angiogenesis (*CDIPT, NOS3, PLCG1*), a 3-gene cluster related to endosome trafficking (labeled PhoX homologous domain), among others. In the decelerated gene set (Fig. 3), the largest cluster was oxidative phosphorylation (25 genes, 9 of which are inferred interactors), followed by mitochondrial translation (10 genes, 1 inferred interactor), regulation of renal output by angiotensin (4 genes), an unnamed cluster (3 genes) related to purine metabolism (*NPR2, PKM, RRM2*), a 3-gene cluster related to RNA polymerase (though labeled as “Abortive elongation of HIV-1 transcript in the absence of Tat”), an unnamed 3-gene cluster related to ribosomes, among others.

**Figure 2.**
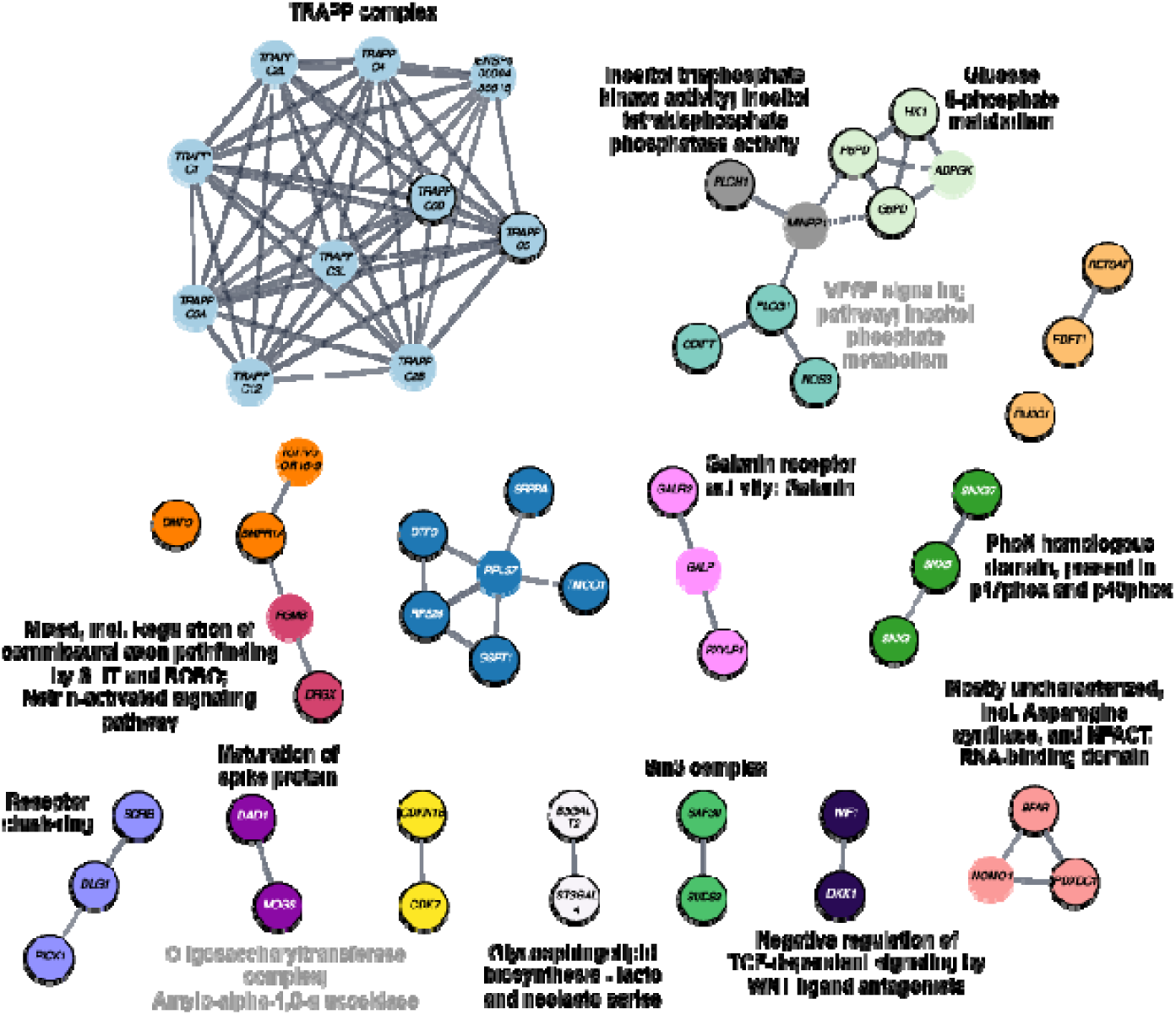
Gene network clusters inferred from distinctly accelerated RER genes, using *Homo sapiens* for annotation. Grey annotations refer to annotations of similar clusters obtained using *Xenopus tropicalis* annotation information. Nodes with no black outline are interactors inferred by STRING that were not present in our original query set. Permalink to the *H. sapiens* STRING network: https://version-12-0.string-db.org/cgi/network?networkId=boDXLAbScr1l. Permalink to the *X. tropicalis* STRING network: https://version-12-0.string-db.org/cgi/network?networkId=bLnhqSov6Wyj

**Figure 3.**
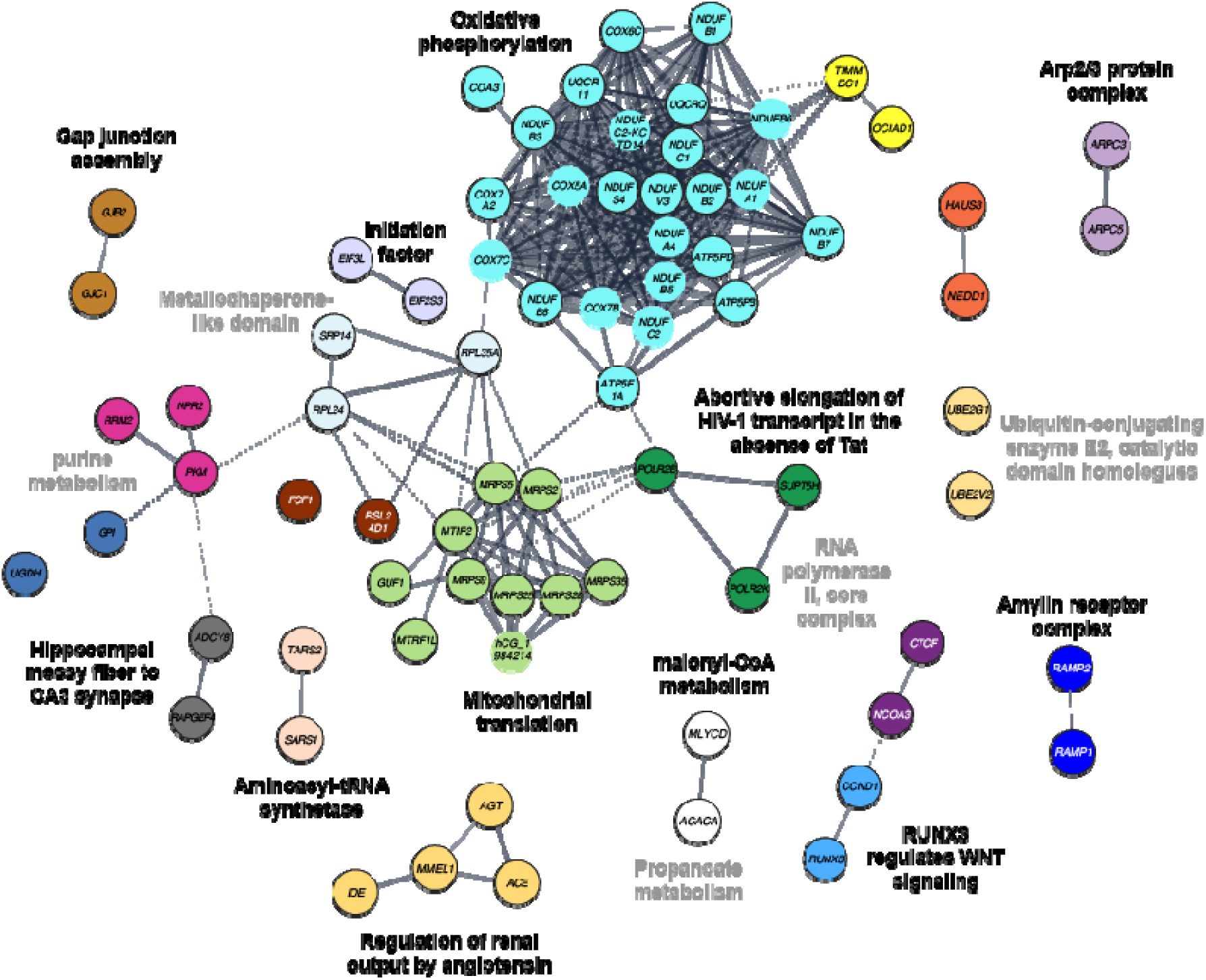
Gene network clusters inferred from distinctly decelerated RER genes, using *Homo sapiens* for annotation. Grey annotations refer to annotations of similar clusters obtained using *Xenopus tropicalis* annotation information. Nodes with no black outline are interactors inferred by STRING that were not present in our original query set. Permalink to the *H. sapiens* STRING network: https://version-12-0.string-db.org/cgi/network?networkId=boDXLAbScr1l. Permalink to the *X. tropicalis* STRING network: https://version-12-0.string-db.org/cgi/network?networkId=b7OklVW1IPb1

## Discussion

In order to respond quickly and appropriately to a stressful environment, organisms can change gene expression, as it underlies virtually all cellular responses to external stress and readjusts the protein content, and therefore the function of the cell. It has been suggested that a hypometabolic state can occur with gene expression changes in only a subset of processes that are key for survival (Storey 2000), such as conservation of energy via two pathways switching to lipid fuel and protein metabolism. Yet, the number of genes that respond during hypometabolism is impressive, as Hudson et al. (2024) observed more than 2500 significantly upregulated or downregulated genes in the skeletal muscle of aestivating frogs (*Cyclorana alboguttata*). This demonstrates that entry into a hypometabolic state requires responses by a huge variety of cell functions (Storey and Storey 2007). Prior studies of aestivation in amphibians have focused on changes in gene expression and resulting fluctuations in metabolites or enzyme activity, and most studies (with a few exceptions including Reilly et al. 2013 and Hudson et al. 2024) have targeted specific processes or genes, such as protein phosphorylation, glycolysis, and the activity of specific transcription factors. In contrast, we take an untargeted approach and use signatures of selection (quantified by distinct evolutionary rates) to identify genes putatively involved in aestivation in amphibians. By doing so, we are not only able to add novel lines of evidence supporting the role of some processes already known to be involved in aestivation, but we can also identify new genes and processes that might underlie hypometabolic states. Here we first review the general themes apparent in our results. Then we revisit our hypotheses and discuss what the different evolutionary signatures we observe can tell us about the aestivation phenotype. Finally, we compare our results to what was previously known to identify how our study contributes to the field.

### Shifts in energy use and production

Two of the main pathways used to produce ATP in cells are anaerobic glycolysis and oxidative phosphorylation that requires oxygen. In aestivating amphibians, aerobic energy production via oxidative phosphorylation is limited due to lower levels of oxygen in tissues from slower breathing rates and reduced blood volume, both are physiological characteristics of aestivation (Storey and Storey 2007). Although oxidative phosphorylation is less costly (in ATP) than glycolysis, some aestivating amphibians switch from oxidative phosphorylation to anaerobic production of ATP, where pyruvate is converted to lactate to regenerate the NAD+ needed to sustain glycolysis (Dawson et al. 2018). However, these patterns seem to vary by tissue type and species. Lactate dehydrogenase (LDH) is responsible for catalyzing the reversible reaction of pyruvate and NADH to lactate and NAD+; yet, there is no consistency in the activity of LDH during aestivation. For example, Cowan et al. (2000) report a decrease of lactate dehydrogenase (LDH) in the brain of *S. couchii* while Wu et al. (2017) report an increase in *X. laevis* and Moreira et al. (2020, 2021) reports no differences on the muscle of aestivating and non-aestivating frogs, *Proceratophrys cristiceps* and *Pleurodema diplolister* respectively (Table S1). Much of the prior research has focused on genes and regulatory mechanisms that are involved in glycolysis or oxidative phosphorylation (more than 20 articles). For example, Katzenback et al. (2014) found differences in kinetics, post-translational modifications, and stabilities of LDH in the liver of *X. laevis*; they also found that its function is maintained at elevated urea levels (150 mM) under dehydration stress.

It is now known that reversible phosphorylation control of pyruvate kinase (PK) is one way of regulating the aerobic versus anaerobic energetic pathways in many species of animals, from aestivating snails to hibernating small mammals (Storey and Storey 1990; Storey 1997; Brooks and Storey 1997; Cowan and Storey 1998). Regulation of PK activity occurs via multiple mechanisms, including tight allosteric regulation and regulation via insulin-mediated cell signaling pathways (Wu et al. 2017). As the action of PK seems to be crucial to deal with dehydration and energy production, and we see little change in the coding sequence in aestivating species (decelerated RER for *PKM*), either its function is similar to the ancestral state or has been modified through other routes such as phosphorylation and gene expression changes (e.g., Luu et al. 2020; Madelaire et al. 2020; Malik and Storey 2009; Pakay et al. 2003; Wu et al. 2017,2020; Zhang et al. 2018). The existing literature seems to support the latter hypothesis. For example, Dawson et al. (2018) showed increased function of PK during severe dehydration in aestivating *X. laevis* compared to control frogs; yet, Wu et al. (2017) found increased pyruvate kinase activity in the brain and kidney but decreased in heart (Table 4, and Table S1). In other studies, a decreased activity was found in skeletal muscle of aestivating frogs (*P. cristiceps*, Moreira et al. 2020; *C. alboguttata*, Reilly et al. 2013; Hudson et al. 2024), and no change in activity in *P. diplolister* (Moreira et al. 2021; Table S1).

Our results further support the importance of energy pathways in aestivating amphibians, as the largest gene cluster identified by STRING in either data set was oxidative phosphorylation (Fig. 3, decelerated). STRING also identified two unnamed clusters (3 genes, 2 genes) related to purine metabolism (including *GPI* and *PKM*, critical to glycolysis), as well as two genes related to fatty-acid metabolism (“malonyl-CoA metabolism”) in the decelerated data set (Fig. 3). *GPI* and *PKM* have been studied previously as mentioned above, whose product is Glucose-6-phosphate isomerase, which is crucial in the inhibition of glycolysis. *GI* was also shown to increase prior to hibernation in the Chinese brown frog (*Rana dybowskii*) (Fan et al. 2022) and during hibernation in the brown bear (*U. arctos*) (Chazarin et al. 2019a). In the accelerated gene set, STRING identified a cluster of genes related to glucose 6-phosphate metabolism (Fig. 2). GO term analyses identified several enriched biological processes related to these pathways including fatty acid and carbohydrate biosynthesis processes as well as ribose phosphate, cyclic nucleotide, and purine ribonucleotide metabolic processes in the decelerated data set and carbohydrate derivative, nucleoside phosphate, pyridine-containing compound, and monosaccharide metabolic processes enriched in the accelerated dataset. Pyridine-containing compound metabolic processes includes NAD+ catabolism, resulting in NADH, and *NAXE*, present in the decelerated dataset, helps keep NADH supplies in the correct format (Manor et al. 2022).

When sufficient NADH and oxygen are available, oxidative phosphorylation is carried out in the mitochondria via five protein complexes, producing relatively large quantities of ATP. Several members of each complex were represented in the decelerated gene set, and none were present in the accelerated gene set, suggesting strong evolutionary conservation of this process: *TIMMDC1*, *NDUFB1*, *NDUFB2*, *NDUFB3* (a top-ten RER; Table 3), *NDUFB7*, *NDUFB8*, *NDUFC1*, *NDUFS4*, *NDUFV3* in Complex I; *DMAC2* in Complex II; *UQCR11* and *UQCRQ* in Complex III; *COA3*, *COX6C*, and *COX7A2* in Complex IV; *ATP5F1A* (another top-ten RER; Table 3), *ATP5PB*, *ATP23*, and *ATP5PD* in complex V. In estivating *C. alboguttata*, the expression of *NDUFB3*, *NDUFS4*, *ATP5PB*, and other related genes are all downregulated, as part of a global repression of ATP production (via oxidative phosphorylation) concomitant with a lower need for energy in the aestivating state (Hudson et al. 2024; Table 4). The mitochondrion itself was an enriched cellular component (Fig. S4, Appendix 4, available online). *MICOS13*, present in the decelerated gene set, is related to the formation of cristae in the mitochondria, which is directly related to its oxidative phosphorylation capacity (Khosravi and Harner 2020). This strong evolutionary conservation might be related to the brief period of activity that most amphibians living in extreme environments have, therefore they have to be absolutely efficient in renewing their energetic reserves. Non-exclusively, if the ancestral function of the oxidative phosphorylation process was adapted to an aestivating lifestyle, there may be strong selection for it to remain relatively unchanged in species that still aestivate.

Many of the genes that are involved in glycolysis change in expression during dormancy (aestivation and hibernation) and hypoxia in other animals, not only in amphibians. The role of G6PD, glucose-6-phosphate dehydrogenase, which hydrolyzes glucose-6-phosphate into a glucose and phosphate group in the PPP, represents a route for the breakdown of carbohydrates besides glycolysis. G6PD has increased activity in the brain and skeletal muscle (but decreased activity in the liver) of aestivating *S. couchii* toads (Cowan et al. 2000), reduced activity in hibernating squirrels (*Spermophilus parryi*) (Smullin and Behrisch, 1981) and was downregulated in diapausing flesh fly pupae (*Sarcophaga crassipalpis*) (Chen et al. 2020). *ACACA*, present in our decelerated gene set, is involved in fatty acid synthesis and has been found to have reduced expression in hibernating grizzly bears (*Ursus arctos horribilis*), black bears (*U. americanus*), Himalayan marmots (*Marmota himalayana*), and tree shrews (*Tupaia belangeri*) (Jansen et al. 2019; Fedorov et al. 2025; Bai et al. 2019; Hou et al. 2023). *HK1*, present in our accelerated gene set and involved in glycolysis, has been shown to have increased expression in the skeletal muscle during aestivation in *C. alboguttata* (Hudson et al. 2024; Table 4), during anoxia in *Trachemys scripta* turtles (Sparks et al. 2021) and *Rattus norvegicus* rats (Waskova-Arnostova et al. 2015), as well as during dormancy in *Apostichopus japonicus* sea cucumbers, and in *Microcebus murinus* mouse lemurs during torpor (Biggar et al. 2015b; Yang et al. 2021). However, *HK1* expression decreased in hibernating grizzly bears (*U. a. horribilis*) and in Horseshoe bats (*Rhinolophus ferrumequinum*) during torpor (Jansen et al. 2019; Xiao et al. 2015), suggesting some variation in the role of glycolysis among species during dormancy. In addition, during dormancy, some organisms transcribe but do not translate certain genes, or pause the glycolysis pathway at later steps, to build up reserves of intermediates that can be accessed immediately upon awakening (Hudson et al. 2024).

Enhancement of antioxidant defenses is proving to be a universal feature of hypometabolism across distant lineages of life (Storey and Storey 2007; Moreira et al. 2017; Giraud-Billoud et al. 2024). In aestivating amphibians, enzymatic and metabolite antioxidant defenses are adaptable systems that can be easily modified in response to stressful conditions (i.e., hot and dry environments) in order to enter a hypometabolic state as well as to exit it when conditions return to suitable conditions (Grundy and Storey, 1998). One antioxidant gene was present in the decelerated gene set: *ATOX1*. In contrast, *NOS3*, which produces nitrous oxide, a source of reactive oxygen species (ROS) (Canty, 2000), was present in the accelerated gene set and was one of the top ten RER genes (Table 2 and 4). *NOS3* was previously found to be downregulated in skeletal muscle in *C. alboguttata* (Reilly et al. 2013). Several genes involved in DNA repair, including *FAM168A* (decelerated), and *SIRT7*, *SSRP1* and *YBX1* (accelerated) were present in our dataset (Fig. 4). In aestivating amphibians, Glucose-6-phosphate dehydrogenase (*G6PD*, accelerated) promotes carbohydrate entry into the PPP to produce sugar phosphates for biosynthesis, producing reduced forms of glutathione and thioredoxin by NADPH-reducing power to drive antioxidant defense. This reducing power is crucial to prevent and minimize oxidative damage to macromolecules during dormancy. G6PD activity has been shown to be increased during aestivation in *S. couchii* and *Rana pipiens* (Cowan et al. 2000; Cowan and Storey 2002). Increased *G6PD* activity during oxidative stress is also present in other species such as land snails (*Otala lactea*), marine mollusks (*Littorina littorea*) and in wood frogs (*R. sylvatica*) (Hermes-Lima et al. 2015; Giraud-Billoud et al. 2024). Complementarily, inhibited and/or reduced G6PD activity has been correlated with reduced antioxidant capacities, even death (Navas and Carvalho 2010). Notably we didn’t find *NRF2* or any *FOXO* genes with distinct RERs, although they are canonical responses to ROS (Giraud-Billoud et al. 2024; Malik and Storey 2009, 2011).

**Figure 4.**
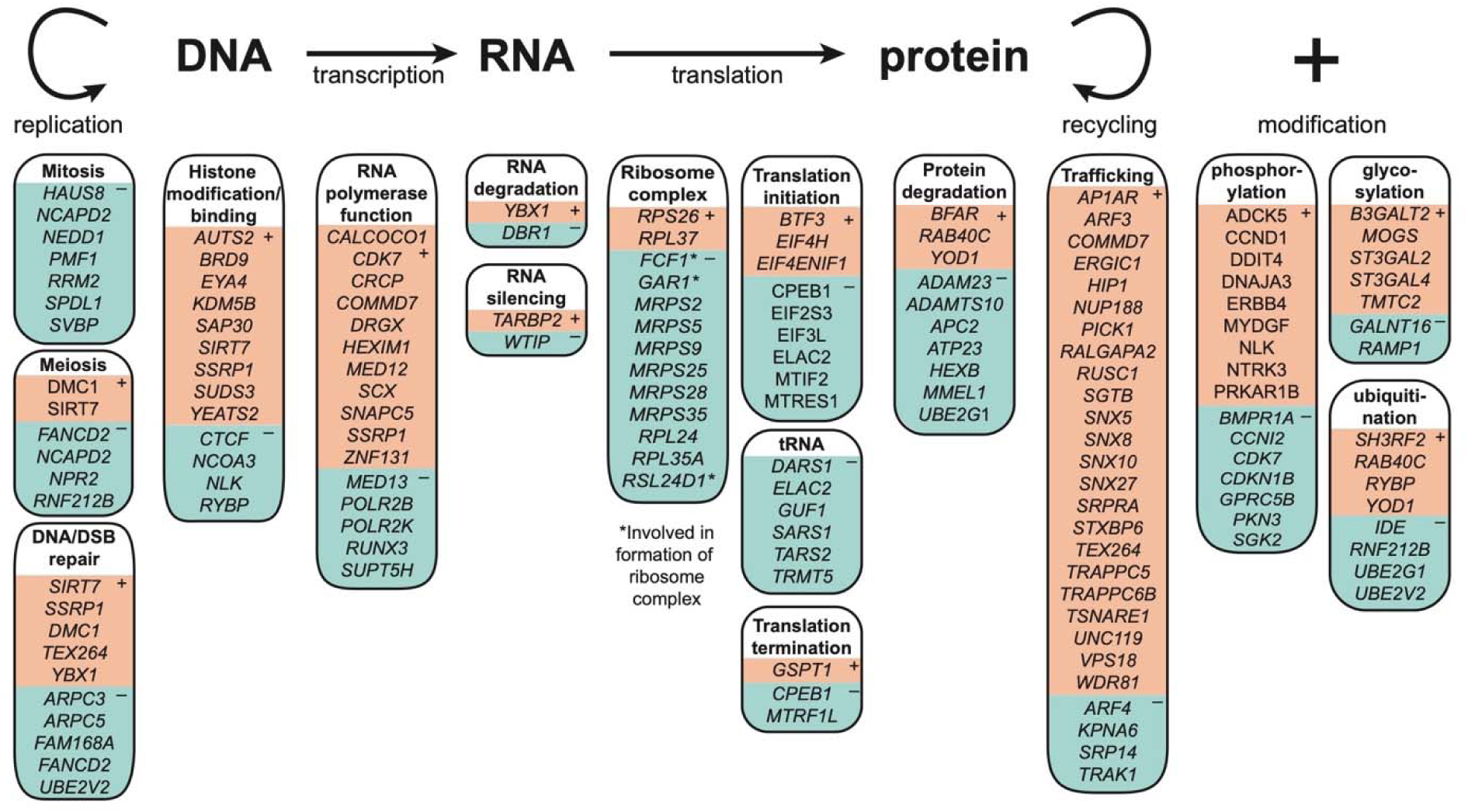
Genes related to the central dogma that we identified to have accelerated (+) or decelerated (–) relative evolutionary rates in aestivating amphibians.

### Transcription and translation may be central targets of selection in aestivating amphibians

Transcription and translation are energetically costly, and during dormancy, animals often reduce these processes to both save energy and avoid generating products that are not in high demand (Storey and Storey 2007; Hawkins and Storey 2020). One way that transcription can be reduced is by increasing methylation or decreasing acetylation of histones; this reduces the accessibility of DNA to transcription factors and ultimately reduces transcription rates (Jiang et al. 2023). STRING identified two members of the Sin3 complex (*SAP30* and *SUDS3*; Fig. 2) involved in histone deacetylation in the accelerated gene set, whose activity would make chromatin more tightly packed and less accessible for transcription (Wang et al. 2023). The Sin3 complex was also studied by Hudson et al. (2008), who found that *SIN3A* and other epigenetic modifiers were upregulated in the skeletal muscle of aestivating *C. alboguttata*. Although histone modification was not identified as an enriched process (e.g., the GO terms ubiquitin-modified histone reader activity, histone binding, or histone deubiquitinase activity are not enriched in our datasets), there were a number of additional genes related to histone binding and accessibility present in the accelerated dataset including *AUTS2*, *BRD9* (top ten RER; Table 2), *EYA4*, *KDM5B*, *SIRT7*, *SSRP1*, and *YEATS2* and in the decelerated dataset, including *CTCF*, *NCOA3*, *NLK*, *RYBP* (Fig. 4). *KDM5B*, a histone demethylase, has been shown to have decreased expression in wood frogs (*R. sylvatica*) during freezing (Taiwo et al. 2024). In addition, two of these genes (*AUTS2*, *RYBP*) are members of the Polycomb group (PcG) proteins, which are a conserved family of epigenetic regulators of gene expression. In the central bearded dragon (*Pogona vitticeps*), PcG plays an important role during hibernation by repressing transcription (Capraro et al. 2019). *SAP30*, involved in the functional recruitment of the class 1 Sin3-histone deacetylase complex (*HDAC*) to the nucleolus which results in a repressed chromatin structure (Kuzmichev et al. 2002), and *AUTS2* were both downregulated in aestivating *C. alboguttata* (Hudson et al. 2024; Table 4).

In reviewing the GO term and STRING results, we noted that many other genes and processes integral to transcription and translation were identified to be enriched in the RER gene sets (Fig. 4; Figs. S2–S4, Appendix 4, available online). For example, “transcription by RNA polymerase III” was a biological process enriched in the accelerated gene set (Fig. S3, Appendix 4, available online). Two genes encoding RNA polymerases present in the decelerated gene set were highlighted by STRING (*POLR2B*, *POLR2K*) along with *SUPT5H*, which is implicated in transcription efficiency (Naguib et al. 2019). *POLR2B*, a catalytic core component of RNA polymerase II, is downregulated in the skeletal muscle in aestivating *C. alboguttata* (Hudson et al. 2024; Table 4). In estivating snails (*Bostryx conspersus*), the expression of RNA polymerases was downregulated (Pourazami 2024). Many transcription factors and proteins related to transcription factor function were also present in the accelerated gene set, including *CALCOCO1*, *CDK7*, *COMMD7*, *DRGX*, *HEXIM1*, *MED12*, *SCX*, *SNAPC5*, *SSRP1*, and *ZNF131*; two were also present in the decelerated gene set (*MED13* and *RUNX3*) (Fig. 4). At least two of these genes have been implicated in responses to hypoxia and hibernation in animals previously: *HEXIM1* is upregulated in heart cells of lab mice (*Mus musculus*) following hypoxia, possibly helping to prevent hypertrophy (Yoshikawa et al. 2012); *MED12* is upregulated during torpor in Himalayan marmots (*M. himalayana*) (Bai et al. 2019).

If produced, mRNA resulting from transcription can be temporarily hidden from translation machinery, for example by being reversibly bound by *CPEB1* (decelerated), which is a mechanism that would allow the buildup of reserve mRNA that can later be translated when protein products are needed (Cheng and Schuh 2024), while avoiding unnecessary energy expense during dormant periods. A few genes related to miRNA and RNA silencing and degradation were also present in the accelerated dataset (*YBX1*, *TARBP2*), while *DBR1* and *WTIP* were in the decelerated dataset (Fig. 4). Expression of *YBX1* was found to be downregulated in aestivating *C. alboguttata* (Hudson et al. 2024, Table 4); this particular gene is involved in translational repression, RNA stabilization, mRNA splicing and transcription regulation. This may seem counter intuitive since protein synthesis is strongly suppressed during aestivation (Storey and Storey 2007; Hawkins and Storey 2020). Nonetheless, the downregulation might be enforced because its secreted form acts as an extracellular mitogen and stimulates cell migration and proliferation.

In contrast to the mixed patterns of selection on genes related to transcription, genes relating to translation were largely characterized by decelerated RERs. An impressive number of the components of the ribosome and mitoribosome were present in the decelerated gene set including *MRPS2*, *MRPS5*, *MRPS9*, *MRPS25*, *MRPS28*, *MRPS35*, *RPL24*, and *RPL35A*; two additional genes, *RPS26* and *RPL37* were present in the accelerated gene set (Fig. 4). Three genes involved in the assembly of the ribosome complex were also identified within the decelerated gene set (*FCF1*, *GAR1*, and *RSL24D1*). Although one may generally expect that translation machinery is under purifying selection, here we find that it is even more conserved in aestivating compared to non-estivating species. A number of genes related to tRNA function, as well as the initiation and termination of translation, were present, mostly in the decelerated gene set (Fig. 4; including *MTRF1L*, one of the top ten decelerated RERs; Table 3). One of these genes, *GUF1*, helps promote mitochondrial protein synthesis under less-than-optimal conditions (Bauerschmitt et al. 2008). The GO terms “translation” and “amino acid activation”, related to amino acids being attached to their respective tRNA, were enriched in the decelerated gene set (Fig. S3), along with their corresponding cellular locations (ribosome, ribonucleoprotein complex, and mitochondrion; Fig. S4, Appendix 4, available online), and associated molecular functions including “catalytic activity, acting on RNA and tRNA”, and “structural constituent of ribosome” (Fig. S3, Appendix 4, available online). One of these decelerated genes, *SARS1*, which encodes the cytosolic seryl-tRNA synthetase (responsible for charging tRNA with serine), had an increased expression in the skeletal muscle of aestivating *C. alboguttata* (Reilly et al. 2013; Table 4). Finally, STRING identified two clusters of genes involved in translation in the decelerated gene set (“Mitochondrial translation,” “initiation factor;” Fig. 3). The expression of translation initiation factors has previously been shown to be altered during periods of dormancy in animals. For example, *EIF4H* (accelerated) is upregulated during torpor in tenrecs (*Tenrec ecaudatus*) (Khudyakov et al. 2021), *EIF2S3* (decelerated) is downregulated in aestivating snails (*Bostryx conspersus*) (Pourazami 2024), and *EIF4ENIF1* (accelerated) is upregulated in the skeletal muscle in aestivating *C. alboguttata* (Hudson et al. 2024; Table 4), promotes miRNA-mediated translational repression and plays a key role in P-bodies to coordinate the storage of translationally inactive mRNAs in the cytoplasm and prevent their degradation (Kamenska et al. 2014; Räsch et al. 2020).

### Signature of selection on intracellular trafficking

While active, cells prioritize growth and efficiency; however, during aestivation, cells must conserve energy and protect themselves from damage. In general, we identified many genes and processes involved in the transport of proteins and macromolecules under distinctly accelerated RERs (Fig. 4; Fig. S3, Appendix 4, available online), suggesting that selection has acted on many aspects of intracellular trafficking in aestivating amphibians (Fig. 5). Thus, it is possible that either survival during aestivation or recovery from aestivation requires more efficient or targeted transport, modification, and/or degradation of macromolecules. In addition to simply moving macromolecules around, intracellular trafficking is also important for membrane remodeling, autophagy, and protein modification. For example, removing aquaporins from cell membranes helps protect against desiccation in a killifish (*Fundulus heteroclitus*) (Tingaud-Sequeira et al. 2013). Further, metabolizing proteins and lipids from cell membranes or from cells undergoing apoptosis can simultaneously provide nutrients while allowing animals to reconfigure their tissues for new environments (Storey and Storey 2004; Hawkins and Storey 2020). In yeast experiencing glucose starvation, cells halt membrane protein recycling, which results in an increased retention of membrane proteins for metabolism and ATP production (Lang et al. 2014). Our STRING analyses highlighted several protein complexes involved in endosome formation, vesicle transport, exocytosis, and membrane fusion processes (see also Fig. 5). The TRAPP (TRAnsport Protein Particle) complex (represented by *TRAPPC5* and *TRAPPC6B*, under accelerated RERs), is involved in docking and/or tethering vesicles to a target membrane (e.g., the ER or the Golgi apparatus) (Sacher et al. 2008). *TRAPPC6B* is the second top accelerated RER gene (Table 2). The SNARE complex (*tSNARE*, *STXBP6*, *VPS18*; all accelerated) helps fuse membranes together, e.g., of an endosome and the Golgi complex (Ungar and Hughson 2003). Sorting nexins (*SNX5*, *SNX8*, *SNX10*, and *SNX27*; all accelerated), highlighted by STRING as a “PhoX homologous domain” cluster, are involved in membrane trafficking and the endosome pathway (Cullen 2008). The Arp2/3 complex (*ARPC3*, *ARPC5*; both decelerated) is involved in forming branched actin, which is necessary to move endosomes (Rotty et al. 2013); *FLNB*, filamin B (a top-ten accelerated RER), is associated with movement along the actin skeleton. Other genes involved in endocytosis (see Fig. 5) were also present in our RER top gene datasets, although they were not highlighted by GO or STRING: *HIP1*, *AP1AR*, *WDR81*, *UNC119*, and *COMMD7* (all accelerated); *RUFY1* (decelerated). A number of genes related to trafficking to the Golgi apparatus, where proteins are subsequently modified and sent for degradation or to the cell membrane, were also present in the top gene sets (*ARF4*, decelerated; *ARF3*, *ERGIC1*, *AP1AR*, all accelerated; Fig. 4). Finally, members of the exocyst complex were also present: *RALGAPA2*, *SYNGR3*, and *VPS18* (all accelerated; *VPS18* is a top-ten RER gene; Table 2); *EXOC1* (decelerated). Thus, the molecular machinery involved in molecule intake, movement, modification, and excretion are generally under accelerated evolutionary rates in aestivating amphibians.

**Figure 5.**
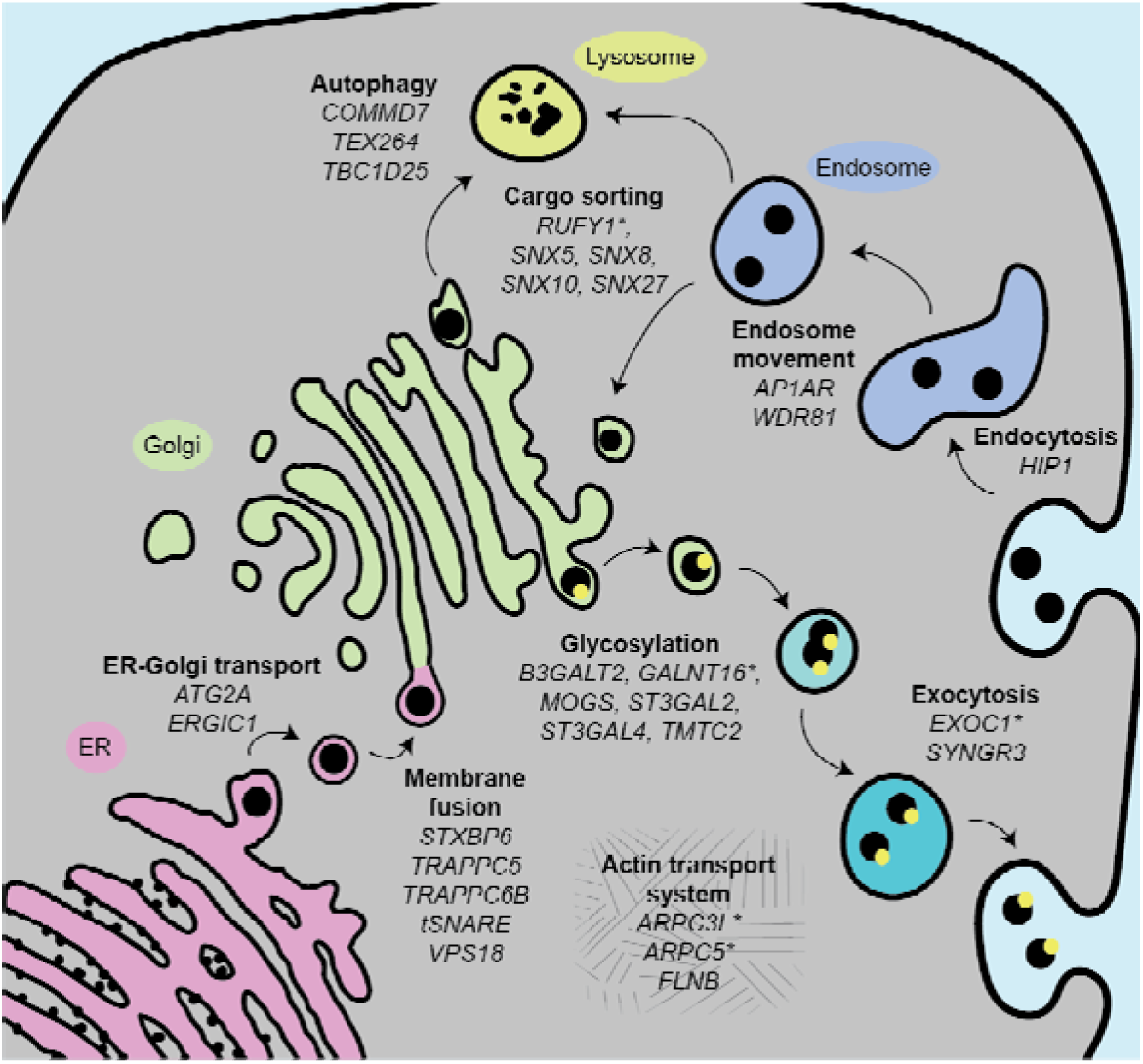
A schematic highlighting some of the genes related to intracellular trafficking that we found to have distinctly accelerated or decelerated (asterisks) relative evolutionary rates in aestivating amphibians. ER, Endoplasmic reticulum.

Related to these functions are the targeting and breakdown of proteins for energy, and/or as a response to ROS damage (see section above). The molecular function GO Terms “metallopeptidase activity” and “metalloendopeptidase activity” were enriched in the decelerated gene sets (Fig. S2, Appendix 4, available online). These are related to protein breakdown. Similarly, polyubiquitination involves tagging a protein with multiple ubiquitin molecules, often then leading to its degradation (Voet et al. 2008). Several ubiquitin enzymes were present in our dataset, including ubiquitin-conjugating enzymes (E2s) *UBE2G1* and *UBE2V2* (both decelerated; Table 4), ubiquitin-protein ligases (E3s) *SH3RF2*, *RAB40C* (both accelerated), and *RNF212B* (decelerated), as well as genes encoding proteins that can remove ubiquitin tags (*YOD1*, accelerated, and *IDE*, decelerated). STRING identified the two E2s as a cluster (Fig. 3) and the RER of *UBE2G1* was the second top in the decelerated gene set (Table 3). Previously it was shown that *UBE2G1* and *UBE2V2* are upregulated in skeletal muscle during aestivation in *C. alboguttata* (Hudson et al. 2024; Table 4) and, more generally, that protein degradation was enriched during torpor in tenrecs (*T. ecaudatus*) (Khudyakov et al. 2021).

Autophagy is another mechanism to recycle nutrients (and reduce damage from ROS) by degrading internal organelles and molecules in endosomes. The biological process GO term “macroautophagy” was enriched in our accelerated RER gene set (Fig. S3, Appendix 4, available online). In total, more than 10 genes involved in autophagy were present in our datasets. Some of these are specifically related to lysosomes or peroxisomes (Fig. 5; accelerated: *ACBD5*, *YOD1*; decelerated: *PSAP*), the formation of autophagosomes (accelerated: *ATG2A*, *TBC1D25*), or ER-phagy (*TEX264*; one of the most accelerated RER genes, Table 2). *DDIT4*, present in the decelerated gene set, has been shown to be downregulated in response to dehydration in skeletal muscle in *C. alboguttata* (Reilly et al. 2013; Table 4), and to be related to autophagosomes (Ho et al. 2020). Membrane fusion is involved in autophagy, and as discussed above, several protein complexes involved in membrane fusion or trafficking were identified in our results.

At a larger scale, apoptosis can also contribute to nutrient metabolism, induced by cellular damage through action by ROS, or through other physiological processes. Three genes with accelerated RERs are related to inhibition of apoptosis (*BFAR*, *DAD1*, *MDM4*; *DAD1* is a top-ten RER) while two with decelerated RERs are related to the promotion of apoptosis (*RUNX3*, *RYBP*). In prior studies, *BFAR* was shown to be downregulated in the skeletal muscle in aestivating *C. alboguttata* (Hudson et al. 2024; Table 4), and *DAD1* was shown to be upregulated during torpor in *Myotis ricketti* bats (Huang et al. 2020) and in hibernating Himalayan marmots (*M. himalayana*) (Bai et al. 2019). A number of other genes with various roles related to homeostasis or indirect interactions with apoptosis genes were present in each gene set (accelerated: *CDK7*, *GRAMD4*, *HIP1*, *NAIF1*; decelerated: *ANP32B*, *DKK1*, *DNAJA3*, *ERBB4*, *RPL35A*). Previously, *ANP32B* was shown to be upregulated in the skeletal muscle in aestivating *C. alboguttata* (Hudson et al. 2024; Table 4). Apoptosis-inducing factor 2 (*AIFM2*) was the gene with the highest increase in expression in the skeletal muscle in aestivating *C. alboguttata* (Hudson et al. 2024). However, although *AIFM2* presented a decelerated relative evolutionary rate in our study, it was not within the top set.

### Other emerging themes

A few unexpected groups of genes were identified. For example, an unnamed cluster of 3 genes containing two related to angiogenesis (*PLCG1*, *NOS3*) was identified by STRING in the accelerated dataset (Fig. 2); similarly, STRING identified a cluster of 4 genes in the decelerated gene set related to the regulation of renal output by angiotensin (Fig. 3). One of these genes, *ACE*, an angiotensin-converting enzyme, was previously identified to be downregulated in response to aestivation in skeletal muscle in *C. alboguttata* (Reilly et al. 2013; Hudson et al. 2024; Table 4). *ACE* inhibitors are commonly used in patients with high blood pressure to block the activation of angiotensin and avoid the constriction of blood vessels and lower blood pressure; they can also cause angioedemas (Cutrell et al. 2023). In Syrian hamsters, decreased *ACE* activity is associated with maintenance of lung tissue during torpor (Talaei et al. 2012). In high-altitude Himalayan marmots (*M. himalayana*), it’s been proposed that changes in the angiogenesis pathway can help avoid over proliferation of blood vessels in response to lower oxygen availability (Bai et al. 2019). A change in the activity of *ACE* may also impact plasma filtration rates in the kidneys (Krečak et al. 2021), adjusting ion and water retention to promote survival during aestivation. The VEGFA-VEGFR2 signaling pathway, involved in angiogenesis, was overrepresented during torpor in tenrecs (*T. ecaudatus*) (Khudyakov et al. 2021). Aestivating amphibians need to respond to low oxygen availability by activating *VEGF* (to stimulate angiogenesis), however it is counter-intuitive to enhance capillary growth since the problem is not oxygen delivery but a lack of oxygen overall. Therefore, upregulation of pigment epithelium-derived factor (*PEDF*) might counteract a *VEGF*-mediated angiogenic response in hypoxic environments (Storey and Storey 2007).

Genes involved in post-translational modifications were also common in our accelerated and decelerated gene sets. Above we discuss ubiquitination, but glycosylation and phosphorylation were also common themes (Fig. 4). Glycosylation is a ubiquitous co- or post-translational protein modification process that affects protein solubility, stability, and function (Lauc and Wuhrer 2017). We identified several genes related to glycosylation in both our accelerated (*B3GALT2*, *MOGS*, *ST3GAL2*, *ST3GAL4*, *TMTC2*) and decelerated (*GALNT16*, *RAMP1*) gene sets (Figs. 4 and 5). Prior studies have shown that *TMTC2* is possibly related to adaptation to high elevation in sheep and pika (Sjodin 2023; Yang et al. 2016). *ST3GAL2* is downregulated in hibernating Daurian Ground Squirrels (*Spermophilus dauricus*), and glycosylation in general was identified as an important process in their ability to avoid muscle atrophy during hibernation (Dang et al. 2020). The GO terms “cellular process”, “glycosylation”, “carbohydrate derivative biosynthetic process”, and “carbohydrate derivative metabolic process” are related to these functions. Protein phosphorylation is another prevalent theme in our top genes (Fig. 4), as well as prior studies of aestivation (e.g., Cowan et al. 1999; Wu et al. 2017; Zhang et al. 2018; Madelaire et al. 2020). We found a prevalence of kinases in our data set including several representatives of the cyclin-dependent kinases (accelerated: *CCND1*, *CDKN1B*; decelerated: *CCNI2*, *CDK7*). *CDKN1B* was found to be downregulated in response to dehydration in skeletal muscle in *C. alboguttata* (Reilly et al. 2013; Table 4). Also present, were *GPRC5B* (top ten accelerated RER; Table 3), which modulates the NF-kB signaling pathway involved in responses to stress, resulting in downstream phosphorylation activity, and *MYDGF* (decelerated), which results in changes in phosphorylation through the MAPK signaling pathway (Luu et al. 2021). Phosphatases, which remove phosphoryl groups from proteins, were not as prevalent in our data (only *EYA4* and *PXYLP1* were present in the accelerated dataset).

Genes related to cell cycle progression were overrepresented in our results. Specifically, in the accelerated gene set, several genes were related to the Wnt-signalling pathway, and the molecular function GO term “beta-catenin binding” was enriched. In addition, two genes in the decelerated gene set related to Wnt-signalling were identified by STRING in “RUNX3 regulates WNT signaling”. More than ten genes associated with cell cycle progression were present in our top gene sets (accelerated: *CCNI2*, *CDKN1B*, *GSPT1*, *HOXD10*, *MDM4*, *SCRIB*, *SIRT7*; decelerated: *ANP32B*, *CCND1*, *DDIT4*, *ERBB4*, *FANCD2*, *FGFR1OP*, *NCAPD2*, *NPR2*, *RRM2*, *RUNX3*, *UBE2V2*, *WTIP*). It was previously shown that *GSPT1* is upregulated in the skeletal muscle in aestivating *C. alboguttata* (Hudson et al. 2024; Table 4); also, *CDKN1B* is upregulated in the testis of a cavefish (*Onychostoma macrolepis*) during torpor (Ma et al. 2024), in killifish (*Nothobranchius furzeri*) during diapause (Hu et al. 2020), and in zebrafish (*Danio rerio*) under hypoxia (Jopling et al. 2012); its action is suggested to inhibit cell proliferation and promote muscle preservation (Ma et al. 2024). *CDK7* and *CDKN1B* were identified as part of an unnamed gene cluster related to the cell cycle by STRING (Fig. 2). The role of cell cycle progression in aestivating amphibians could be two-fold: 1) limit tissue growth during aestivation and 2) promote rapid tissue growth during the awakening period prior to reproduction. More studies of genes involved in cell-cycle progression could help provide insight into mechanisms promoting or preventing the development of cancers. In general, cancer is a rare phenomenon in amphibians (Torres-Dimas et al. 2022; Compton et al. 2025).

Finally, we note several other groups of genes present in our data: homeobox genes, important for development (*DRGX*, *EVX2*, *HHEX*, *HOXB2*, *HOXB9*, *HOXD10*, *NKX2-5*; all accelerated); genes involved in cell-cell adhesion or junctions (accelerated: *CLDN11*, *CLDN7*, *DLG1*, *CLRN1*, *PARVA*, *TEX264*; decelerated: *ADAM23*, *CDHR3*, *DSCAM*, *GJB2*, *GJC1*, *MICOS13*, *WTIP*), and a few genes involved in sperm function (all decelerated: *MMEL1*, *SPACA6*, *SPATA6*) or hormones (accelerated: *GALP*, *GALR2*, also highlighted by STRING in Fig. 2, and SCT; decelerated: *ACE*, *IDE*, *NPR2*). *HHEX*, a transcriptional repressor that regulates the differentiation of endothelial and blood cells was found downregulated in the skeletal muscle in aestivating *C. alboguttata* (Hudson et al. 2024; Table 4). *SCT* is involved in mechanisms of satiation and homeostasis in regard to pH in the stomach (top 10; Table 2). In hibernating amphibians, galanin (encoded by *GALP*, binding to receptor GALR2) may be linked to shifts in metabolic fuel sources or decreased responses to stress (González Nicolini et al. 1998). A gap junction protein (*GJB1*) was found to be upregulated in hibernating thirteen-lined ground squirrels (*Ictidomys tridecemlineatus*) (Hampton et al. 2013) and muscle structure was a theme in gene expression changes in the skeletal muscle in aestivating *C. alboguttata* (Reilly et al. 2013). *ADAM23* gene expression has been associated with cell-cell and cell-matrix interactions and was also found to be downregulated in response to dehydration in skeletal muscle in *C. alboguttata* (Reilly et al. 2013; Hudson et al. 2024; Table 4). Genes involved in sperm function and hormones may be related to the unique life histories of aestivating amphibians, which require rapid periods of reproduction following long aestivation periods (Shalan et al. 2004; Wilsterman et al. 2021).

### Relative evolutionary rates of genes associated with aestivation, revisiting the hypotheses

We hypothesized that the selection patterns in our data would provide insight into both the mechanisms permitting aestivation as well as the evolutionary signatures associated with its origin and maintenance. Given that aestivation is thought to be an ancestral state of amphibians (Ochoa-Ochoa et al. 2025), we hypothesized (H1 & H3) that some genes would be highly conserved and show decelerated relative evolutionary rates (RERs). Yet, the ecological and phylogenetic diversity of the amphibian species that aestivate may result in positive selection on some genes, diversifying their functions in accordance to species-specific ecological and physiological needs (H2 & H3). Parsing apart these patterns can provide insight into possible core and auxiliary mechanisms of the aestivation phenotype.

Most of the processes represented by genes in our results show a mixture of accelerated and decelerated rates, supporting H3 and demonstrating that the retained ancestral phenotype relies both on a core set of highly conserved processes and a set of auxiliary processes that have diversified with time. First, the genes in our datasets that are related to oxidative phosphorylation and translation are all under decelerated RERs. The ubiquity of decelerated relative evolutionary rates in genes linked to these processes suggests that there has been especially strong negative selection on translation and oxidative phosphorylation in aestivating amphibians, and identifies them as a core conserved process in relation to aestivation. Having well-functioning machinery for oxidative phosphorylation and translation could be important both during aestivation, to preserve energy, and during active periods, to quickly recover, reproduce, and renew the energetic reserves (Vieira de Andrade et al. 2016). Nevertheless, these patterns suggest that among aestivating amphibians, it is beneficial or necessary for the genes underlying these processes to remain relatively unchanged from their ancestral sequences. We note that other processes, such as phosphorylation and epigenetic modifications could alter these processes without changing the coding sequences.

Two processes were predominantly under accelerated RERs: the homeobox genes and most of the genes involved in macromolecule transport (Figs. 4 and 5). These patterns suggest that development and intracellular transport is diverse among aestivating amphibian lineages. We hypothesize that many of the genes related to the metabolism of cell membrane lipids and proteins are under accelerated evolution in aestivating amphibians because of the variety of needs of and resources available to each species, necessitating different solutions to conserve energy, recover from the aestivation state, and limit the increase of ROS (Jiang et al. 2023).

We note one additional intriguing pattern. The RERs of the three genes related to inhibition of apoptosis were all accelerated while that of the two genes related to the promotion of apoptosis were decelerated. If this pattern reflects the evolutionary history of selection on apoptosis, it suggests that in aestivating amphibians, the mechanisms promoting apoptosis are highly conserved while those inhibiting it are diverse.

### Comparison between our results and prior studies

In our literature review, we found a predominance of targeted studies (Table S1), which focus on a subset of genes, proteins, or metabolites putatively involved in the hypometabolic state, rather than untargeted studies, which review all available genetic or physiological information to subsequently identify which genes, proteins, metabolites, or functions change the most between states. We only identified two untargeted studies: Reilly et al. (2013) and Hudson et al. (2024), both on *C. alboguttata*. Many articles employed biological assays to measure enzyme activity or content as an approximation of the metabolic scope of a tissue, because metabolic depression can be observed by reduced enzyme levels or function (Flanigan et al. 1990).

Our untargeted approach led to the identification of TRAPP, SNARE, and Arp2/3 complexes, which is novel and highlights the possible importance of intracellular trafficking and membrane remodeling in the aestivation phenotype (Fig. 5). The identification of many genes involved in angiotensin regulation is also novel, although studies of hibernation and hypoxia in other organisms have pointed to its possible involvement (Khudyakov et al. 2021; Bai et al. 2019; Talaei et al. 2012). The role of cell-cycle progression is also of interest and has not been expansively investigated previously, although it was indicated as important in *C. alboguttata* (Reilly et al. 2013). Both cell proliferation and angiogenesis are in part regulated by the NFKB1 transcription factor, which has been shown to be involved in the aestivation response in amphibians (Maistrovski 2011). Similarly, the two other untargeted studies on aestivation in amphibians that we are aware of, Reilly et al. (2013) and Hudson et al. (2024), also provided insight into novel mechanisms, such as the involvement of antiapoptotic mechanisms and the possibility that tissues retain pools of mRNA and glycolysis intermediates poised for rapid translation and catabolism when the animal awakes.

A comparison of distinct evolutionary rates with genes previously identified as associated with the aestivation state identified 30 overlapping genes in one or more of the following patterns: those that were accelerated and had increased activity in aestivators; accelerated and decreased; decelerated and increased; and, finally, decelerated and decreased (see also Appendix 4, available online). While our study focused on evolutionary rates, which is a reflection of negative or positive selection, the prior studies that we reviewed measure gene expression and/or protein quantity. Differences in evolutionary rates and in gene/protein activity represent distinct aspects of how protein functions change over time. Further investigation will be necessary to fully interpret the patterns observed in Table 4, especially because only two prior studies on *C. alboguttata* (Reilly et al. 2013, Hudson et al. 2024) assessed changes in gene expression using an untargeted approach; thus; the data are likely an incomplete picture of gene/protein activity changes associated with aestivation. However, we can propose a few hypotheses. First, genes under accelerated evolutionary rates and increased activity may have a novel function in the aestivation state. We based this on the fact that accelerated evolutionary rates reflect a relative increase in the rate of change in the amino acid sequence, which likely has a functional consequence, and because increased activity of an inherently different protein is consistent with a novel function. Genes in this category include *DLG1*, *EIF4ENIF1*, *GSPT1*, and *HK1*, which range in function from development to translation and phosphorylation (see Appendix 4, available online). On the other hand, genes under accelerated evolutionary rates and relatively decreased activity in the aestivation state may have a novel function in the active state. Genes in this category were predominantly involved in transcription regulation (*AUTS2*, *BFAR*, *CDKN1B*, *HHEX*, *NOS3*, *SAP30*, and *YBX1*). Second, genes that are under decelerated evolutionary rates and increased activity may reflect the “highly conserved core processes” we describe above that could be necessary for the aestivation state. Genes in this category were related to protein synthesis and modification including ubiquitination, transcription, translation, and glycolysis. The largest category of genes was the decelerated evolutionary rates and decreased activity; these, in contrast, may represent conserved processes necessary for the active state in aestivators. Genes in this category had the greatest variation in function including oxidative phosphorylation, fatty acid biosynthesis, glycolysis, cell cycle regulation, and the angiotensin-converting enzyme (Table 4, Appendix 4, available online). Third, there were three genes that are under decelerated evolutionary rates and had evidence of both decreased and increased activity in different studies and/or tissues (*GPI*, *JCHAIN*, and *PKM*); we hypothesize that such a pattern implies a highly conserved core process with tissue-specific changes in regulation, rather than in the coding sequence, of these genes (e.g., see our discussion above of pyruvate kinase in the *Shifts in energy use and production* section).

## Conclusion

There has been great scientific curiosity as well as medical interest in understanding how animals survive and recover from periods of dormancy while maintaining functional organ systems. For example, induced suspended animation is a hypometabolic state of medical interest because it can preserve organ function during life-threatening conditions (Hartmann et al. 2017). Nonetheless, this procedure causes other physiological problems. However, if some of the physiological processes that aestivators or hibernators perform in order to survive a hypometabolic state could be induced in human medicine, the procedure could be improved (Hartmann et al. 2017). In our review of the literature outside of amphibians, many genes that we highlight have been investigated in terms of cancer tumor states of dormancy versus proliferation (Ho et al. 2020), atrial fibrosis (Yoshikawa et al. 2012), and organ atrophy (Chazarin et al. 2019b). However, most studies of any type of dormancy have been focused on hibernation in mammals. Here we identified some processes overlapping with prior targeted studies on energy shifts in amphibians and other species, but we also highlight possible new processes such as protein and membrane trafficking (TRAPP, SNARE, and Arp2/3 complexes). Thus, studying the aestivation phenotype in amphibians presents novel insight into processes that permit such a fascinating adaptation, which could hold promise for human medicine.

## Data and code accessibility statement

All data and code are available as supplementary materials

## Supplementary Material

**Appendix A1.** Script for gene trees based on NCBI gene annotations.

**Appendix A2.** R script for RER convergence analyses.

**Appendix A3.** R script for gene ontology analyses.

**Appendix A4.** Additional results and figures.

**Appendix A5.** List of clusters and associated annotation files from the Search Tool for the Retrieval of Interacting Genes/Proteins database STRING (Szklarczyk et al. 2023).

**Table S1.** List of genes, metabolites, miRNAs, or proteins analyzed during aestivation or water restriction found in the literature review (see Methods section).

**Table S2.** List of genes and proteins used for the comparison between literature and RER results.

**Table S3.** Full results of the RER analyses.

**Table S4.** List of genes and terms that were significantly enriched in the GO analyses based on the RER top gene set compared to the background gene set.

## References

Akat E., H. Arıkan, and B. Göçmen. 2014. Histochemical and biometric study of the gastrointestinal system of *Hyla orientalis* (Bedriaga, 1890) (Anura, Hylidae). Eur J Histochem 58:291–295. doi:10.4081/ejh.2014.2452.

Alexa A., and J. Rahnenführer. 2024. Gene set enrichment analysis with topGO. doi:10.18129/B9.bioc.topGO.

Amey A.P., and G.C. Grigg. 1995. Lipid-reduced evaporative water loss in two arboreal hylid frogs. Comp Biochem Physiol III A:283–291.

AmphibiaWeb. 2024. AmphibiaWeb. University of California, Berkeley, CA, USA. Available at https://amphibiaweb.org.

Anderson R. C., R. P. Bovo, C. E. Eismann, A. A. Menegario, and D. V. Andrade. 2017. Not good, but not all bad: dehydration effects on body fluids, organ masses, and water flux through the skin of *Rhinella schneideri* (Amphibia, Bufonidae). Physiol Biochem Zool 90:313–320. doi:10.1086/690189.

Bai L., B. Liu, C. Ji, S. Zhao, S. Liu, Rong Wang, W. Wang et al. 2019. Hypoxic and cold adaptation insights from the Himalayan marmot genome. iScience 11:519–530. doi:10.1016/j.isci.2018.11.034.

Bauerschmitt H., S. Funes, and J.M. Herrmann. 2008. The membrane-bound GTPase Guf1 promotes mitochondrial protein synthesis under suboptimal conditions. J Biol Chem 283:17139–17146. doi:10.1074/jbc.M710037200.

Bayomy M.F.F., A.G. Shalan, S.D. Bradshaw, P.C. Withers, T. Stewart, and G. Thompson. 2002. Water content, body weight and acid mucopolysaccharides, hyaluronidase and b-glucuronidase in response to aestivation in Australian desert frogs. Comp Biochem Physiol Part A 131:881–892.

Berman D.I., N.A. Bulakhova, and E.N. Meshcheryakova. 2019. The Siberian wood frog survives for months underwater without oxygen. Sci Rep 9. doi:10.1038/s41598-018-31974-6.

Biggar K. K., Y. Biggar, and K. B. Storey. 2015a. Identification of a novel dehydration responsive gene, drp10, from the African clawed frog, Xenopus laevis. J Exp Zool A Ecol Genet Physiol 323:375–381. doi:10.1002/jez.1930.

Biggar K.K., C.W. Wu, S.N. Tessier, J. Zhang, F. Pifferi, M. Perret, and K.B. Storey. 2015b. Modulation of gene expression in key survival pathways during daily torpor in the gray mouse lemur, *Microcebus murinus*. Genom Proteom Bioinform13:111–118. doi:10.1016/j.gpb.2015.03.001.

Biggar Y., W.A. Ingelson-Filpula, and K.B. Storey. 2022. Pro- and anti-apoptotic microRNAs are differentially regulated during estivation in *Xenopus laevis*. Gene 819. doi:10.1016/j.gene.2022.146236.

Booth D.T. 2006. Effect of soil type on burrowing behavior and cocoon formation in the green-striped burrowing frog, *Cyclorana alboguttata*. Can J Zool 84:832–838. doi:10.1139/Z06-062.

Borah B.K., Z. Renthlei, and A.K. Trivedi. 2020. Hypothalamus but not liver retains daily expression of clock genes during hibernation in terai tree frog (*Polypedates teraiensis*). Chronobiol Int 37:485–492. doi:10.1080/07420528.2020.1726373.

Capraro A., D. O’Meally, S.A. Waters, H.R. Patel, A. Georges, and P.D. Waters. 2019. Waking the sleeping dragon: gene expression profiling reveals adaptive strategies of the hibernating reptile *Pogona vitticeps*. BMC Genomics 20. doi:10.1186/s12864-019-5750-x.

Cartledge V.A., P.C. Withers, K.A. McMaster, G.G. Thompson, and S.D. Bradshaw. 2006. Water balance of field-excavated aestivating Australian desert frogs, the cocoon-forming *Neobatrachus aquilonius* and the non-cocooning *Notaden nichollsi* (Amphibia: Myobatrachidae). J Exp Biol 209:3309–3321. doi:10.1242/jeb.02393.

Chauvet J., Y. Ouedraogo, G. Michel, and R. Acher. 1993. Vasotocin and hydrin 2 (vasotocinyl-gly) in the African toad *Bufo regularis*: study under various environmental conditions. Comp Biochem Physiol 104:497–502.

Chazarin B., K.B. Storey, A. Ziemianin, S. Chanon, M. Plumel, I. Chery, C. Durand et al. 2019a. Metabolic reprogramming involving glycolysis in the hibernating brown bear skeletal muscle. Front Zool 16. doi:10.1186/s12983-019-0312-2.

Chazarin B., A. Ziemianin, A.L. Evans, E. Meugnier, E. Loizon, J.M. Arnemo, J.E. Swenson et al. 2019b. Limited oxidative stress favors resistance to skeletal muscle atrophy in hibernating brown bears (*Ursus arctos*). Antioxidants 8. doi:10.3390/antiox8090334.

Cheng S., and M. Schuh. 2024. Two mechanisms repress cyclin B1 translation to maintain prophase arrest in mouse oocytes. Nat Comm15. doi:10.1038/s41467-024-54161-w.

Chikina M., N. Clark, A. Kowalczyk, W. Mao, W. Meyer, and R. Partha. 2025. RERconverge: analysis of convergence between organismal traits and DNA/protein sequences. Rpackage. doi:10.18129/B9.bioc.topGO.

Chikina M., J.D. Robinson, and N.L. Clark. 2016. Hundreds of genes experienced convergent shifts in selective pressure in marine mammals. Mol Biol Evol 33:2182–2192. doi:10.1093/molbev/msw112.

Compton Z.T., W. Mellon, V.K. Harris, S. Rupp, D. Mallo, S.E. Kapsetaki, M. Wilmot et al. 2025. Cancer prevalence across vertebrates. Cancer Discov 15:227–244. doi:10.1158/2159-8290.CD-24-0573.

Cowan K.J., J.A. Macdonald, J.M. Storey, K., and B. Storey. 2000. Metabolic reorganization and signal transduction during estivation in the spadefoot toad. Exp Biol Online 5:61–85.

Cowan K.J., and K.B. Storey. 2002. Urea and KCI have differential effects on enzyme activities in liver and muscle of estivating versus nonestivating species. Biochem Cell Biol 80:745–755. doi:10.1139/o02-144.

Cramp R.L., C.E. Franklin, and E.A. Meyer. 2005. The impact of prolonged fasting during aestivation on the structure of the small intestine in the green-striped burrowing frog, *Cyclorana alboguttata*. Acta Zool 86:13–24. doi:10.1111/j.0001-7272.2005.00180.x.

Cramp R.L., S.M. Kayes, E.A. Meyer, and C.E. Franklin. 2009. Ups and downs of intestinal function with prolonged fasting during aestivation in the burrowing frog, *Cyclorana alboguttata*. J Exp Biol 212:3656–3663. doi:10.1242/jeb.027748.

Cullen P.J. 2008. Endosomal sorting and signaling: an emerging role for sorting nexins. Nat Rev Mol Cell Biol 9:574–582. doi:10.1038/nrm2427.

Cutrell S., I.S. Alhomoud, A. Mehta, A.H. Talasaz, B. Van Tassell, and D.L. Dixon. 2023. ACE-inhibitors in hypertension: a historical perspective and current insights. Curr Hypertens Rep 25:243–250. doi:10.1007/s11906-023-01248-2.

Dang K., H.J. Yu, S.H. Xu, T.R. Ma, H.P. Wang, Y. Li, Z. Li, and Y.F. Gao. 2020. Remarkable homeostasis of protein sialylation in skeletal muscles of hibernating Daurian ground squirrels (*Spermophilus dauricus*). Front Physiol 11. doi:10.3389/fphys.2020.00037.

Dawson N.J., Y. Biggar, A.I. Malik, and K.B. Storey. 2018. Increased transcript levels and kinetic function of pyruvate kinase during severe dehydration in aestivating African clawed frogs, *Xenopus laevis*. Comp Biochem Physiol B Biochem Mol Biol 224:245–252. doi:10.1016/j.cbpb.2018.01.003.

Dyer S.C., O. Austine-Orimoloye, A.G. Azov, M. Barba, I. Barnes, V.P. Barrera-Enriquez, A. Becker et al. 2025. Ensembl 2025. Nucleic Acids Res 53:D948–D957. doi:10.1093/nar/gkae1071.

Emms D.M., and S. Kelly. 2019. OrthoFinder: phylogenetic orthology inference for comparative genomics. Genome Biol 20. doi:10.1186/s13059-019-1832-y.

Fan C., Y. Liu, Y. Wang, A. Zhang, W. Xie, H. Zhang, Q. Weng, and M. Xu. 2022. Expression of glycogenic genes in the oviduct of Chinese brown frog (*Rana dybowskii*) during pre-brumation. Theriogenology 185:78–87. doi:10.1016/j.theriogenology.2022.03.006.

Fedorov V.B., A. Garreau, Ø. Tøien, B.M. Barnes, and A. V. Goropashnaya. 2025. Transcriptome remodeling and adaptive preservation of muscle protein content in hibernating black bears. bioRxiv preprint 1–22. doi:10.1101/2025.03.06.641932.

Flanigan J., P. Withers, K. Storey, and M. Guppy. 1990. Changes in enzyme binding and activity during aestivation in the frog *Neobatrachus pelobatoides*. Comp Biochem Physiol 96B:67–71.

Flanigan J.E., and M. Guppy. 1997. Metabolic depression and sodium-potassium ATPase in the aestivating frog, *Neobatrachus kunapalari*. J Comp Physiol B 167:135–47.

Fuery C.J., P.C. Withers, A.A. Hobbs, and M. Guppy. 1998. The role of protein synthesis during metabolic depression in the Australian desert frog *Neobatrachus centralis*. Comp Biochem Physiol 119:469–476.

Giraud-Billoud M., D.C. Moreira, M. Minari, A. Andreyeva, E.G. Campos, J.M. Carvajalino-Fernández et al. 2024. Review: evidence supporting the ‘preparation for oxidative stress’ (POS) strategy in animals in their natural environment. Comp Biochem Physiol A Mol Integr Physiol 293. doi:10.1016/j.cbpa.2024.111626.

González Nicolini M. V, A.A. Orezzoli, and M.J. Villar. 1998. An inmunohistochemical study of temperature-related changes in galanin and nitric oxide synthase immunoreactivity in the hypothalamus of the toad. Gen Comp Endocrinol 110:175–181.

Hampton M., R.G. Melvin, and M.T. Andrews. 2013. Transcriptomic analysis of brown adipose tissue across the physiological extremes of natural hibernation. PLoS One 8. doi:10.1371/journal.pone.0085157.

Hartmann C., B. Nussbaum, E. Calzia, P. Radermacher, and M. Wepler. 2017. Gaseous mediators and mitochondrial function: the future of pharmacologically induced suspended animation? Front Physiol 8. doi:10.3389/fphys.2017.00691.

Hawkins L.J., and K.B. Storey. 2020. Advances and applications of environmental stress adaptation research. Comp Biochem Physiol A Mol Integr Physiol 240:e110623. doi:10.1016/j.cbpa.2019.110623.

Hermes-Lima M., D.C. Moreira, G.A. Rivera-Ingraham, M. Giraud-Billoud, T.C. Genaro-Mattos, and É.G. Campos. 2015. Preparation for oxidative stress under hypoxia and metabolic depression: revisiting the proposal two decades later. Free Radic Biol Med 89:1122–1143. doi:10.1016/j.freeradbiomed.2015.07.156.

Ho K.H., P.H. Chen, C.M. Chou, C.M. Shih, Y.T. Lee, C.H. Cheng, and K.C. Chen. 2020. A Key Role of DNA damage-inducible transcript 4 (*DDIT4*) connects autophagy and *GLUT3*-mediated stemness to desensitize temozolomide efficacy in glioblastomas. Neurotherapeutics 17:1212–1227. doi:10.1007/s13311-019-00826-0.

Hou D.M., T. Jia, H.J. Wang, Z.K. Wang, and W.L. Zhu. 2023. Differentially expressed genes analysis of brown adipose tissue during cold acclimation in male tree shrews (*Tupaia belangeri*) based on RNA-seq. Pak J Zool 55:591–600. doi:10.17582/journal.pjz/20211210131248

Huang W., C.C. Liao, Y. Han, J Lv, M. Lei, Y. Li, Q. Lv, et al. 2020. Co-activation of Akt, Nrf2, and NF-κB signals under UPRER in torpid Myotis ricketti bats for survival. Commun Biol 3. doi:10.1038/s42003-020-01378-2.

Hudson N. J., T. G. A. Lonhienne, C. E. Franklin, G. S. Harper, and S. A. Lehnert. 2008. Epigenetic silencers are enriched in dormant desert frog muscle. J Comp Physiol B 178:729–734. doi:10.1007/s00360-008-0261-0.

Hudson N.J., R.L. Cramp, and C.E. Franklin. 2024. Dramatic genome-wide reprogramming of mRNA in hypometabolic muscle. Comp Biochem Physiol B Biochem Mol Biol 272. doi:10.1016/j.cbpb.2024.110952.

Hughes M.K., and A.L. Hughes. 1993. Evolution of duplicate genes in a tetraploid animal, *Xenopus laevis*. Mol Biol Evol 10:1360–1369.

IUCN. 2024. The IUCN red list of threatened species. Version 2023-1. http://www.iucnredlist.org. Downloaded October 11, 2024.

Jansen H.T., S. Trojahn, M.W. Saxton, C.R. Quackenbush, B.D. Evans Hutzenbiler, O.L. Nelson, O.E. Cornejo, C.T. Robbins, and J.L. Kelley. 2019. Hibernation induces widespread transcriptional remodeling in metabolic tissues of the grizzly bear. Commun Biol 2. doi:10.1038/s42003-019-0574-4.

Jiang C., K.B. Storey, H. Yang, and L. Sun. 2023. Aestivation in nature: physiological strategies and evolutionary adaptations in hypometabolic states. Int J Mol Sci 24:e14093. doi:10.3390/ijms241814093.

Jones R.M. 1980. Metabolic consequences of accelerated urea synthesis during seasonal dormancy of spadefoot toads, *Scaphiopus couchi* and *Scaphiopus multip*licatus. J Exp Zool 212:255–267. doi:10.1002/jez.1402120212.

Jopling C., G. Suñé, A. Faucherre, C. Fabregat, J. Carlos, and I. Belmonte. 2012. Hypoxia induces myocardial regeneration in zebrafish. Circulation 126:3017–3027. doi:10.1161/CIRCULATIONAHA.

Jørgensen C.B. 1997. Urea and amphibian water economy. Comp Biochem Physiol 117:161–170.

Kamenska A., W. T. Lu, D. Kubacka, H. Broomhead, N. Minshall, M. Bushell, and N. Standart. 2014. Human 4E-T represses translation of bound mRNAs and enhances microRNA-mediated silencing. Nucleic Acids Res 42:3298–3313. doi:10.1093/nar/gkt1265.

Katoh K., and D.M. Standley. 2013. MAFFT multiple sequence alignment software version 7: improvements in performance and usability. Mol Biol Evol 30:772–780. doi:10.1093/molbev/mst010.

Katzenback B.A., N.J. Dawson, and K.B. Storey. 2014. Purification and characterization of a urea sensitive lactate dehydrogenase from the liver of the African clawed frog, X*enopus laevis*. J Comp Physiol B 184:601–611. doi:10.1007/s00360-014-0824-1.

Kay R.R. 2021. Macropinocytosis: biology and mechanisms. Cells Dev 168:203713. doi:10.1016/j.cdev.2021.203713

Khosravi S., and M.E. Harner. 2020. The MICOS complex, a structural element of mitochondria with versatile functions. Biol Chem 401:765–778. doi:10.1515/hsz-2020-0103.

Khudyakov J.I., M.D. Treat, M.C. Shanafelt, J.S. Deyarmin, B.A. Neely, and F. van Breukelen. 2021. Liver proteome response to torpor in a basoendothermic mammal, *Tenrec ecaudatus*, provides insights into the evolution of homeothermy. Am J Physiol Regul Integr Comp Physiol 321:R614–R624. doi:10.1152/ajpregu.00150.2021.

Kowalczyk A., W.K. Meyer, R. Partha, W. Mao, N.L. Clark, and M. Chikina. 2019. RERconverge: an R package for associating evolutionary rates with convergent traits. Bioinformatics 35:4815–4817. doi:10.1093/bioinformatics/btz468.

Krečak I., M. Morić Perić, I. Zekanović, H. Holik, B. Coha, V. Gverić-Krečak, and M. Lucijanić. 2021. Beneficial effect of *ACE* inhibitors on kidney function in polycythemia vera. Wien Klin Wochenschr 133:808–815. doi:10.1007/s00508-021-01812-3.

Kumar S., M. Suleski, J.M. Craig, A.E. Kasprowicz, M. Sanderford, M. Li, G. Stecher, and S.B. Hedges. 2022. TimeTree 5: An expanded resource for species divergence times. Mol Biol Evol 39:msac174. doi:10.1093/molbev/msac174.

Kuzmichev A., Y. Zhang, H. Erdjument-Bromage, P. Tempst, and D. Reinberg. 2002. Role of the Sin3-histone deacetylase complex in growth regulation by the candidate tumor suppressor p33ING1. Mol Cell Biol 22:835–848. doi:10.1128/MCB.22.3.835-848.2002.

Lang M.J., J.Y. Martinez-Marquez, D.C. Prosser, L.R. Ganser, D. Buelto, B. Wendland, and M.C. Duncan. 2014. Glucose starvation inhibits autophagy via vacuolar hydrolysis and induces plasma membrane internalization by down-regulating recycling. J Biol Chem 289:16736–16747. doi:10.1074/jbc.M113.525782.

Lauc G., and M. Wuhrer. 2017. High throughput glycomics and glycoproteomics. Methods and protocols. Humana Press, New York, NY.

Liedtke H.C., F. Cruz, J. Gómez-Garrido, D. Fuentes Palacios, M. Marcet-Houben, M. Gut, T. Alioto, T. Gabaldón, and I. Gomez-Mestre. 2022. Chromosome-level assembly, annotation and phylome of *Pelobates cultripes*, the western spadefoot toad. DNA Research 29:dsac013. doi:10.1093/dnares/dsac013.

Luu B.E., L.J. Hawkins, and K.B. Storey. 2021. Insights from a vertebrate model organism on the molecular mechanisms of whole-body dehydration tolerance. Mol Cell Biochem 476:2381–2392. doi:10.1007/s11010-021-04072-x.

Luu B.E., and K.B. Storey. 2015. Dehydration triggers differential microRNA expression in *Xenopus laevis* brain. Gene 573:64–69. doi:10.1016/j.gene.2015.07.027

Luu B.E., S. Wijenayake, A.I. Malik, and K.B. Storey. 2018. The regulation of heat shock proteins in response to dehydration in *Xenopus laevis*. Cell Stress Chaperones 23:45–53. doi:10.1007/s12192-017-0822-9

Luu B.E., Y. Zhang, and K.B. Storey. 2020. The regulation of *Akt* and *FoxO* transcription factors during dehydration in the African clawed frog (*Xenopus laevis*). Cell Stress Chaperones 25:887–897. doi:10.1007/s12192-020-01123-y.

Ma Y., Y. Chen, Y. Li, S. Chen, C. Zhu, Q. Liu, L. Li, et al. 2024. Seasonal modulation of the testis transcriptome reveals insights into hibernation and reproductive adaptation in *Onychostoma macrolepis*. Fish Physiol Biochem 50:2083–2097. doi:10.1007/s10695-024-01335-4.

Madelaire C.B., F.R. Gomes, and I. Sokolova. 2020. Biomarker-based assessment of the muscle maintenance and energy status of anurans from an extremely seasonal semi-arid environment, the Brazilian Caatinga. Comp Biochem Physiol A Mol Integr Physiol 240. doi:10.1016/j.cbpa.2019.110590.

Maistrovski Y. 2011. Regulation of anti-apoptotic pathways in skeletal muscle and liver of an estivating species, Xenopus laevis. Carleton University, Ottawa, ON.

Malik A.I., J.M. Storey, and K.B. Storey. 2023. Regulation of the unfolded protein response during dehydration stress in African clawed frogs, *Xenopus laevis*. Cell Stress Chaperones 28:529–540. doi:10.1007/s12192-022-01275-z.

Malik A.L., and K.B. Storey. 2009. Activation of extracellular signal-regulated kinases during dehydration in the African clawed frog, *Xenopus laevis*. J Exp Biol 212:2595–2603. doi:10.1242/jeb.030627

Malik A.L., and K.B. Storey. 2011. Transcriptional regulation of antioxidant enzymes by *FoxO1* under dehydration stress. Gene 485:114–119. doi:10.1016/j.gene.2011.06.014.

Manor J., D. Calame, C. Gijavanekar, K. Fisher, J. Hunter, E. Mizerik, C. Bacino, F. Scaglia, and S.H. Elsea. 2022. NAXE deficiency: a neurometabolic disorder of NAD(*P*)HX repair amenable for metabolic correction. Mol Genet Metab 136:101–110. doi:10.1016/j.ymgme.2022.04.003.

Mayhew W.W. 1968. Biology of desert amphibians and reptiles. Pp. 195–356 in Brown G., ed., Desert biology: special topic on the physical and biological aspects of arid regions. Academic Press, New York, NY.

McClanahan L. 1972. Changes in body fluids of burrowed Spadefoot toads as a function of soil water potential. Copeia 1972:209–216.

Meyer W. 2024. RERconverge analysis walkthrough. 1–29 pp.

Minh B.Q., H.A. Schmidt, O. Chernomor, D. Schrempf, M.D. Woodhams, A. Von Haeseler, R. Lanfear, and E. Teeling. 2020. IQ-TREE 2: new models and efficient methods for phylogenetic inference in the genomic era. Mol Biol Evol 37:1530–1534. doi:10.1093/molbev/msaa015.

Moreira D.C., M.F. Oliveira, L. Liz-Guimarães, N. Diniz-Rojas, É.G. Campos, and M. Hermes-Lima, 2017. Current trends and research challenges regarding “preparation for oxidative stress”. Front Physiol 8:702. doi:10.3389/fphys.2017.00702.

Moreira D.C., J.M. Carvajalino-Fernández, W.P. Silva, F. Kuzniewski, C.A. Navas, J.E. de Carvalho, and M. Hermes-Lima. 2020. Preparation for oxidative stress in *Proceratophrys cristiceps* (Anura, Odontophrynidae) naturally estivating in the Brazilian caatinga. Sci Total Environ 723:137957. doi:10.1016/j.scitotenv.2020.137957.

Moreira D.C., J.M. Carvajalino-Fernández, C.A. Navas, J.E. de Carvalho, and M. Hermes-Lima. 2021. Metabolic and redox biomarkers in skeletal muscle underlie physiological adaptations of two estivating anuran species in a south American semi-arid environment. Front Physiol 12:769833. doi:10.3389/fphys.2021.769833

Naguib A., T. Sandmann, F. Yi, R.J. Watts, J.W. Lewcock, and W.E. Dowdle. 2019. *SUPT4H1* depletion leads to a global reduction in RNA. Cell Rep 26:45–53.e4. doi:10.1016/j.celrep.2018.12.004.

Naranjo M., S.A. Breedon, and K.B. Storey. 2023. Cardiac microRNA expression profile in response to estivation. Biochimie 210:22–34. doi:10.1016/j.biochi.2023.01.001.

Navas C.A., and J.E. Carvalho. 2010. Aestivation molecular and physiological aspects. Springer-Verlag, Berlin Heidelberg, Germany.

Ochoa-Ochoa L.M., J.D. Vásquez-Restrepo, M. Masache, and R.D. Tarvin. 2025. Slowing down: a macroevolutionary approach to the hypometabolic strategies of amphibians. bioRxiv preprint doi:10.1101/2025.07.19.665691.

O’Leary N.A., E. Cox, J.B. Holmes, W. R. Anderson, R. Falk, V. Hem, M.T.N. Tsuchiya et al. 2024. Exploring and retrieving sequence and metadata for species across the tree of life with NCBI Datasets. Sci Data 11. doi:10.1038/s41597-024-03571-y.

Pakay J.L., A.A. Hobbs, S.R. Kimball, and M. Guppy. 2003. The role of eukaryotic initiation factor 2α during the metabolic depression associated with estivation. J Exp Biol 206:2363–2371. doi:10.1242/jeb.00422.

Paradis E., and K. Schliep. 2019. Ape 5.0: an environment for modern phylogenetics and evolutionary analyses in R. Bioinformatics 35:526–528. doi:10.1093/bioinformatics/bty633.

Patnaik P., and D. Das Sahoo. 2021. Variations in oxidative stress and antioxidant defense level during different phases of hibernation in common Asian toad, *Duttaphrynus melanostictus*. Biol Open 10. doi:10.1242/bio.058567.

Pinder A.W., K.B. Storey, and G.R. Ultsch. 1992. Estivation and hibernation. Pp. 250–274 in M.E. Feder and W.W. Burggren, eds. Environmental physiology of the amphibians. The University of Chicago Press, Chicago, IL.

Portik D.M., J.W. Streicher, and J.J. Wiens. 2023. Frog phylogeny: a time-calibrated, species-level tree based on hundreds of loci and 5,242 species. Mol Phylogenet Evol 188. doi:10.1016/j.ympev.2023.107907.

Pourazami S. 2024. Snails in a dry sauna: a transcriptomic tale of Bostryx conspersus. PhD diss. Norwegian University of Science and Technology - NTNU, Trondheim, Norway.

R Core Team. 2024. R: a language and environment for statistical computing.

Räsch F., R. Weber, E. Izaurralde, and C. Igreja. 2020. 4E-T-bound mRNAs are stored in a silenced and deadenylated form. Genes Dev 34:847–860. doi:10.1101/gad.336073.119.

Reilly B.D., D.I. Schlipalius, R.L. Cramp, P.R. Ebert, and C.E. Franklin. 2013. Frogs and estivation: transcriptional insights into metabolism and cell survival in a natural model of extended muscle disuse. Physiol Genomics 45:377–388. doi:10.1152/physiolgenomics.00163.2012.

Revell L.J. 2024. phytools 2.0: an updated R ecosystem for phylogenetic comparative methods (and other things). PeerJ 12:e16505. doi:10.7717/peerj.16505.

Rossi G. S., R. L. Cramp, P. A. Wright, and C. E. Franklin. 2020. Frogs seek hypoxic microhabitats that accentuate metabolic depression during dormancy. J Exp Biol 223(2):jeb218743. doi:10.1242/jeb.218743.

Rotty J.D., C. Wu, and J.E. Bear. 2013. New insights into the regulation and cellular functions of the ARP2/3 complex. Nat Rev Mol Cell Biol 14:7–12. doi:10.1038/nrm3492

Sacher M., Y.G. Kim, A. Lavie, B.H. Oh, and N. Segev. 2008. The TRAPP complex: insights into its architecture and function. Traffic 9:2032–2042. doi:10.1111/j.1600-0854.2008.00833.x.

Sadowski-Fugitt L.M., C.R. Tracy, K.A. Christian, and J.B. Williams. 2012. Cocoon and epidermis of Australian *Cyclorana* frogs differ in composition of lipid classes that affect water loss. Physiol Biochem Zool 85:40–50. doi:10.1086/663695.

Seymour R.S. 1973. Gas exchange in spadefoot toads beneath the ground. Copeia 1973:452–460.

Shalan A.G., S.D. Bradshaw, P.C. Withers, G. Thompson, M.F.F. Bayomy, F.J. Bradshaw, and T. Stewart. 2004. Spermatogenesis and plasma testosterone levels in western Australian burrowing desert frogs, *Cyclorana platycephala, Cyclorana maini*, and *Neobatrachus sutor*, during aestivation. Gen Comp Endocrinol 136:90–100. doi:10.1016/j.ygcen.2003.12.005.

Shannon P., A. Markiel, O. Ozier, N.S. Baliga, J.T. Wang, D. Ramage, N. Amin, B. Schwikowski, and T. Ideker. 2003. Cytoscape: a software environment for integrated models of biomolecular interaction networks. Genome Res 13:2498–2504. doi:10.1101/gr.1239303.

Sjodin B. 2023. Unravelling climate and high elevation adaptation in the American pika (Ochotona princeps) through a comparative genomics framework. PhD diss. The University of British Columbia, Vancouver.

Smith M. E., and S. M. Secor. 2017. Physiological responses to fasting and estivation for the three-toed amphiuma (*Amphiuma tridactylum*). Physiol Biochem Zool 90:240–256. 10.1086/689216

Sparks K. 2021. Gene expression of hypoxia-inducible factor (HIF-1Α), its regulators, and its putative downstream targets in ventricle and telencephalon of the anoxia-tolerant turtle (Trachemys scripta). PhD diss. University of Alaska Anchorage, Anchorage.

Storey K.B. 2000. Turning down the fires of life: metabolic regulation of hibernation and estivation. Comp Biochem Physiol B Biochem Mol Biol 126:S90. doi:10.1016/s0305-0491(00)80178-9.

Storey K.B. 2002. Life in the slow lane: molecular mechanisms of estivation. 133. 733–754 pp.

Storey, K.B., and J.M. Storey. 2004. Metabolic rate depression in animals: transcriptional and translational controls. Biol Rev 79:207–233. doi:10.1017/S1464793103006195

Storey, K.B., and J.M. Storey. 2007. Tribute to P. L. Lutz: putting life on “pause” - molecular regulation of hypometabolism. J Exp Biol 210:1700–1714. doi:10.1242/jeb.02716

Storey, K.B., and J.M. Storey. 2010. Metabolic Regulation and Gene Expression During Aestivation. Pp. 25–45 in Navas C.A. and J.E. Carvalho, eds. Aestivation: molecular and physiological aspects. Springer-Verlag, Berlin Heidelberg, Germany.

Storey, K.B., and J.M. Storey. 2012. Aestivation: signaling and hypometabolism. J Exp Biol 215:1425–1433. doi:10.1242/jeb.054403.

Supek F., M. Bošnjak, N. Škunca, and T. Šmuc. 2011. Revigo summarizes and visualizes long lists of gene ontology terms. PLoS One 6. doi:10.1371/journal.pone.0021800.

Szklarczyk D., R. Kirsch, M. Koutrouli, K. Nastou, F. Mehryary, R. Hachilif, A.L. Gable et al.2023. The STRING database in 2023: protein-protein association networks and functional enrichment analyses for any sequenced genome of interest. Nucleic Acids Res 51:D638–D646. doi:10.1093/nar/gkac1000.

Taiwo O.O., S.A. Breedon, and K.B. Storey. 2024. Epigenetic regulation by histone methylation and demethylation in freeze-tolerant frog kidney. Cell Biochem Funct 42:e70036. doi:10.1002/cbf.70036.

Talaei F., H.R. Bouma, M.N. Hylkema, A.M. Strijkstra, A.S. Boerema, M. Schmidt, and R.H. Henning. 2012. The role of endogenous *H2S* formation in reversible remodeling of lung tissue during hibernation in the Syrian hamster. J Exp Biol 215:2912–2919. doi:10.1242/jeb.067363.

The UniProt Consortium. 2025. UniProt: the universal protein knowledgebase in 2025. Nucleic Acids Res 53:D609–D617. doi:10.1093/nar/gkae1010.

Tingaud-Sequeira A., J.J. Lozano, C. Zapater, D. Otero, M. Kube, R. Reinhardt, and J. Cerdà. 2013. A rapid transcriptome response is associated with desiccation resistance in aerially-exposed killifish embryos. PLoS One 8:e64410. doi:10.1371/journal.pone.0064410.

Torres-Dimas E., A. Cruz-Ramírez, and R.M. Bermúdez-Cruz. 2022. Cancer in Amphibia, a rare phenomenon? Cell Biol Int 46:1992–1998. doi:10.1002/cbin.11888.

Ungar D., and F.M. Hughson. 2003. SNARE protein structure and function. Annu Rev Cell Dev Biol 19:493–517.

Uno Y. 2021. Inference of evolution of vertebrate genomes and chromosomes from genomic and cytogenetic analyses using amphibians. Chromosome Sci 24:3–12. doi:10.11352/scr.24.3.

Vieira de Andrade D., C.R. Bevier, and J.E. Carvalho. 2016. Amphibian and reptile adaptations to the environment. Interplay between physiology and behavior. Taylor & Francis Group, Boca Raton, FL.

Voet D., J.G. Voet, and C.W. Pratt. 2008. Fundamentals of biochemistry: life at the molecular level. 3rd Edition. John Wiley & Sons, Inc., Hoboken, NJ.

Wang C., Z. Guo, C. Chu, Y. Lu, X. Zhang, and X. Zhan. 2023. Two assembly modes for SIN3 histone deacetylase complexes. Cell Discov 9. doi:10.1038/s41421-023-00539-x.

Waskova-Arnostova P., B. Elsnicova, D. Kasparova, D. Hornikova, F. Kolar, J. Novotny, J. Zurmanova, and Z.J. Cardioprotective. 2015. Cardioprotective adaptation of rats to intermittent hypobaric hypoxia is accompanied by the increased association of hexokinase with mitochondria. J Appl Physiol 119:1487–1493. doi:10.1152/japplphysiol.01035.2014.

Wilsterman K., M.A. Ballinger, and C.M. Williams. 2021. A unifying, eco-physiological framework for animal dormancy. Funct Ecol 35:11–31. doi:10.1111/1365-2435.13718.

Wu C.W., K.K. Biggar, and K.B. Storey. 2013. Dehydration mediated microRNA response in the African clawed frog *Xenopus laevis*. Gene 529:269–275. doi:10.1016/j.gene.2013.07.064

Wu C.W., S.N. Tessier, and K.B. Storey. 2017. Regulation of the insulin-Akt signaling pathway and glycolysis during dehydration stress in the African clawed frog *Xenopus laevis*. Biochem Cell Biol 95:663–671. doi:10.1139/bcb-2017-0117.

Wu C.W., S.N. Tessier, and K.B. Storey. 2020. Dehydration stress alters the mitogen-activated-protein kinase signaling and chaperone stress response in *Xenopus laevis*. Comp Biochem Physiol B Biochem Mol Biol 246–247. doi:10.1016/j.cbpb.2020.110461.

Xiao Y., Y. Wu, K. Sun, H. Wang, B. Zhang, S. Song, Z. Du et al. 2015. Differential expression of hepatic genes of the greater horseshoe bat (*Rhinolophus ferrumequinum*) between the summer active and winter torpid states. PLoS One 10:e0145702. doi:10.1371/journal.pone.0145702.

Yang J., W.R. Li, F.H. Lv, S.G. He, S.L. Tian, W.F. Peng, Y.W. Sun, et al. 2016. Whole-genome sequencing of native sheep provides insights into rapid adaptations to extreme environments. Mol Biol Evol 33:2576–2592. doi:10.1093/molbev/msw129.

Yang Q., X. Zhang, Z. Lu, R. Huang, N.T. Tran, J. Wu, F. Yang et al. 2021. Transcriptome and metabolome analyses of sea cucumbers *Apostichopus japonicus* in southern China during the summer aestivation period. J Ocean Univ China 20:198–212. doi:10.1007/s11802-021-4482-0.

Yoshikawa N., N. Shimizu, T. Maruyama, M. Sano, T. Matsuhashi, K. Fukuda, M. Kataoka et al. 2012. Cardiomyocyte-specific overexpression of *HEXIM1* prevents right ventricular hypertrophy in hypoxia-induced pulmonary hypertension in mice. PLoS One 7:e52522. doi:10.1371/journal.pone.0052522

Zhang Y., B.E. Luu, and K.B. Storey. 2018. FoxO4 activity is regulated by phosphorylation and the cellular environment during dehydration in the African clawed frog, *Xenopus laevis*. Biochim Biophys Acta Gen Subj 1862:1721–1728. doi:10.1016/j.bbagen.2018.05.002

