## Appendices A1-A3 for "Whole-genome analyses point to new candidate genes underlying aestivation in amphibians": Appendix_A1_Genes.sh

```
#########################################################
################### RER clean script ####################
################# RDT & LOO 14 May 2025 #################
#########################################################

# This pipeline has 4 steps
# 1. PULL ORTHOLOGS
#		Input: <species>, <taxon group> 
#		Software needed: datasets and dataformat from NCBI SRA toolkit
#		Output: fasta files of proteins from all species in <taxon group> matching annotated genes for <species>
# 3. PULL ISOFORMS
#		Input: unzipped output files from PULL ORTHOLOGS
#		Software needed: BioPython
#		Output: fasta files with one protein sequence per species per gene, plus matching TSVs noting which isoform was pulled
# 3. ESTIMATE TREES
#		Input: constraint tree, output files from PULL ISOFORMS
#		Software needed: iqtree2, mafft
#		Output: a protein sequence alignment and tree for each ortholog
# 4. REROOT TREES
#		Input: output files from ESTIMATE TREES, list of species to root tree
#		Software needed: R (ape and phytools packages)
#		Output: a rerooted tree for each ortholog


#########################################################
################## 1. PULL ORTHOLOGS ####################
#########################################################

# to get orthologs for amphibian genomes, we 1) pulled down a gene set from xenopus tropicalis, 
# 2) then used those gene names to find orthologs from other genomes 

# download all genes from xenopus
mkdir genes_2025-05-14
cd genes_2025-05-14/

# the following commands are fast and can be run in the interactive node
datasets download gene taxon "xenopus tropicalis" --include protein # 29,183 gene records 

# generate metadata for gene list
dataformat tsv gene --package ncbi_dataset.zip --fields tax-name,gene-id,symbol,group-id > gene_list.txt 
cat gene_list.txt | awk '{print $4}' |sort > symbol_list.txt # right format
rm ncbi_data.zip # clean folder
mkdir sequences

# download gene records from all amphibian genomes for each gene symbol
# this takes >48 hours and should be submitted as a job; could be made parallel if needed
# if you want to increase the speed, remove the "LOC" genes from the symbol_list.txt before running
for i in `cat symbol_list.txt`
do
	datasets download gene symbol ${i} --ortholog amphibia --filename "sequences/${i}.zip"
done

## unzip into folder with name of gene
# takes 15 minutes
cd sequences
for i in `cat /global/scratch/users/rdtarvin/ncbi_ortho/genes_2025-05-14/symbol_list.txt`
do
	unzip -o ${i}.zip -d ${i} 
done
cd ..

#########################################################
################### 2. PULL ISOFORMS  ###################
#########################################################

# in some cases, multiple isoforms were available for an ortholog
# thus, we selected the longest isoform for downstream analyses

## I gave these instructions to ChatGPT
# I have >10,000 fasta files. Each file is for a specific gene. Each file contains a list 
# of sequences from different species. The header has this format: 
# >XP_072274703.1 MAP2 [organism=Pyxicephalus adspersus] [GeneID=140335714] [isoform=X8] 
# Some species (labeled organism) have multiple sequences (called isoforms) in a file. 
# I want to choose the longest one for each species. The script should be in python using 
# BioPython. I want to iteratively go through each file; for each, a new output file should 
# be made with only one sequence per species.

## it gave me this file, which I have edited for directory locations and a few other corrections
# takes  minutes
module load python
python pull_isoforms.py

###################################
#####     pull_isoforms.py   ######
from Bio import SeqIO
import os
import re

# Set input and output directories
input_dir = "."
output_dir = "primary_isoform"
os.makedirs(output_dir, exist_ok=True)

# Regex to extract organism from header
organism_pattern = re.compile(r'\[organism=(.+?)\]')
geneid_pattern = re.compile(r'\[GeneID=(\d+)\]')

# Base path where all gene folders are located
base_dir = "sequences"  # or wherever your gene directories are located

with open("/global/scratch/users/rdtarvin/ncbi_ortho/genes_2025-05-14/symbol_list.txt") as f:
    gene_names = [line.strip() for line in f]

# Iterate over all FASTA files
    for gene in gene_names:
        input_path = os.path.join(base_dir, gene, "ncbi_dataset", "data", "protein.faa")
        if not os.path.isfile(input_path):
            print(f"Skipping {gene}: file not found at {input_path}")
            continue

        print(f"Processing {input_path}")

        #input_path = os.path.join(input_dir, gene)
        output_path = os.path.join(output_dir, "protein_"+gene+".faa")

        tsv_output_path = os.path.splitext(output_path)[0] + ".tsv"

        longest_per_species = {}

        for record in SeqIO.parse(input_path, "fasta"):
            header = record.description
        
        # Extract fields
            org_match = organism_pattern.search(header)
            geneid_match = geneid_pattern.search(header)

            if not org_match:
                continue

            organism = org_match.group(1)
            organism_clean = organism.replace(" ", "_")
            accession = record.id
            geneid = geneid_match.group(1) if geneid_match else "NA"

        # Modify record
            record.id = organism_clean
            record.description = f"{accession} GeneID={geneid}"

        # Keep only longest per species
            current_best = longest_per_species.get(organism)
            if current_best is None or len(record.seq) > len(current_best['record'].seq):
                longest_per_species[organism] = {
                    'record': record,
                    'accession': accession
                }

     # Write FASTA
        filtered_records = [entry['record'] for entry in longest_per_species.values()]
        SeqIO.write(filtered_records, output_path, "fasta")

    # Write TSV: species and accession
        with open(tsv_output_path, "w") as tsv_out:
            tsv_out.write("species\taccession\n")
            for organism, entry in longest_per_species.items():
                tsv_out.write(f"{organism}\t{entry['accession']}\n")

print("Done: FASTAs with accession and gene ID in headers, plus matching TSVs.")

###################################


#########################################################
################## 3. ESTIMATE TREES  ###################
#########################################################

# once we had all ortholog sequences, we needed to estimate one tree per gene
# RER requires that the trees have the same topology, so we used a tree
# from Portik et al. 2023 pruned to have the species of interest to constrain
# phylogenetic inference

###################################
########## pruned.tree ############
((Rhinatrema_bivittatum:105.550084,(Microcaecilia_unicolor:105.550084,Geotrypetes_seraphini:105.550084)):165.773038,((Pleurodeles_waltl:126.073007,Ambystoma_mexicanum:126.073006):125.126994,(Bombina_bombina:172.076736,(Xenopus_tropicalis:166.170958,((Spea_bombifrons:127.216106,Pelobates_fuscus:127.216108):32.472808,((((Eleutherodactylus_coqui:62.016805,(Ranitomeya_imitator:61.48752,((Bufo_bufo:12.809321,Bufo_gargarizans:12.809321):48.178857,Engystomops_pustulosus:60.988179):0.499342):0.529285):0.527491,(Dendropsophus_ebraccatus:41.887083,Hyla_sarda:41.887082):20.657216):44.50102,Pseudophryne_corroboree:107.045316):7.914059,(Hyperolius_riggenbachi:84.686689,(Pyxicephalus_adspersus:62.06699,(Nanorana_parkeri:55.549748,Rana_temporaria:55.549748):6.517242):22.619699):30.272686):44.729541):6.482042):5.905779):79.123264):20.123121);

# updated to have additional genomes that were annotated as of 2025/05/23, then changed Rana catesbeiana to Aquarana catesbeiana
((Rhinatrema_bivittatum:105.550084,(Microcaecilia_unicolor:105.550084,Geotrypetes_seraphini:105.550084)):165.773038,((Pleurodeles_waltl:126.073007,Ambystoma_mexicanum:126.073006):125.126994,(Ascaphus_truei:179.341108,(Bombina_bombina:172.076736,(((Xenopus_tropicalis:43.884641,Xenopus_laevis:43.88464):55.468469,Hymenochirus_boettgeri:99.353109):66.817848,((Spea_bombifrons:127.216106,(Pelobates_fuscus:17.851064,Pelobates_cultripes:17.851064):109.365044):32.472808,(((((Pristimantis_euphronides:51.854766,Eleutherodactylus_coqui:51.854766):10.162039,((Phyllobates_terribilis:30.774834,(Dendrobates_tinctorius:27.366076,Ranitomeya_imitator:27.366075):3.408757):30.712688,(((Bufo_bufo:12.809321,Bufo_gargarizans:12.809321):8.890904,Bufotes_viridis:21.700225):39.287953,Engystomops_pustulosus:60.988179):0.499342):0.529285):0.527491,(Dendropsophus_ebraccatus:41.887083,Hyla_sarda:41.887082):20.657216):44.50102,Pseudophryne_corroboree:107.045316):7.914059,(Hyperolius_riggenbachi:84.686689,(Pyxicephalus_adspersus:62.06699,(Nanorana_parkeri:55.549748,((Rana_temporaria:25.672777,Aquarana_catesbeiana:25.672777):11.780926,Staurois_parvus:37.453704):18.096045):6.517242):22.619699):30.272686):44.729541):6.482042):5.905779):7.264372):71.858892):20.123121);

###################################


# then we used the slurm script below to submit a job to infer all trees using iqtree2
# this takes > 48h

###################################
########## trees.slurm ############

#!/bin/bash
#SBATCH --job-name=trees
#SBATCH --account=co_rosalind
#SBATCH --partition=savio2_htc
#SBATCH --nodes=1
#SBATCH --time=72:00:00
#SBATCH --output=trees_%j.out
#SBATCH --error=trees_%j.err

## Command(s) to run:
module load python
conda activate iqtree
bash align_and_tree.sh

###################################

###################################
######## align_and_tree.sh ########
#!/bin/bash

# Input directory containing your raw sequence files
INPUT_DIR="primary_isoform"

# Output directory for aligned files
OUTPUT_DIR="trees"

# Create the output directory if it doesn't exist
mkdir -p "$OUTPUT_DIR"

# Iterate through each file in the input directory
for file in "$INPUT_DIR"/*.faa; do
    # Extract the filename without the extension for output naming
    filename=$(basename "$file" .faa)

    # Align the file using MAFFT
    mafft "$file" > "$OUTPUT_DIR/$filename.mafft.aln"
    
    # Remove any extra tips from the constraint tree
    python prune_tree2.py "$filename"

    # Analyze the aligned file using IQ-TREE & a constrained topology
    iqtree2 -s "$OUTPUT_DIR/$filename.mafft.aln" -T AUTO -g "pruned_trees/$filename.pruned.tree" --redo

done
###################################

# this pruning script is necessary because the -g option won't run if the tree has more tips than the alignment

###################################
########## prune_tree.py ##########

import os
import sys
from Bio import SeqIO
from ete3 import Tree

# Path to your constraint tree
constraint_tree_path = "pruned.tree"

# gene name 
filename = sys.argv[1]

# Directory with alignment files
alignment_file = os.path.join('trees', filename+".mafft.aln")

# Load the constraint tree once
original_tree = Tree(constraint_tree_path, format=1)

# Output directory for pruned trees
output_dir = "pruned_trees"
os.makedirs(output_dir, exist_ok=True)

# Extract gene name from filename
gene_name = os.path.basename(alignment_file).split('_')[1].split('.')[0]

# Extract taxa from alignment file
taxa_in_alignment = set()
for record in SeqIO.parse(alignment_file, "fasta"):
    taxon = record.id.split()[0]  # Take first part: Dendropsophus_ebraccatus
    taxa_in_alignment.add(taxon)

# Copy and prune the tree
pruned_tree = original_tree.copy()
all_tree_leaves = set(pruned_tree.get_leaf_names())

# Remove tips not in the alignment
tips_to_prune = all_tree_leaves - taxa_in_alignment
if tips_to_prune:
    pruned_tree.prune(list(taxa_in_alignment), preserve_branch_length=True)

# Save pruned tree
output_path = os.path.join(output_dir, filename+ ".pruned.tree")
pruned_tree.write(outfile=output_path)

print("Pruned tree saved for",filename)
###################################


#########################################################
#################### 4. REROOT TREES ####################
#########################################################

# For RER to run correctly, the trees should all be rooted at the same place

#### with chatgpt wrote R script to reroot the tree
####################### reroot.R ###################################
# Load libraries
library(ape)
library(phytools)

# Define list of taxa to use for MRCA-based rooting
target_taxa <- c(
  "Rhinatrema_bivittatum",
  "Microcaecilia_unicolor",
  "Geotrypetes_seraphini"
)

# Find all *.treefile files in the current directory
tree_files <- list.files(pattern = "\\.treefile$")

# Loop through each tree file
for (tree_file in tree_files) {
  cat("Processing:", tree_file, "\n")
  
  # Read the tree
  tree <- read.tree(tree_file)
  
  # Check which target taxa are present
  present_taxa <- intersect(target_taxa, tree$tip.label)
  missing_taxa <- setdiff(target_taxa, present_taxa)
  
  if (length(present_taxa) == 0) {
    cat("  Skipping - no target taxa present.\n")
    next
  }
  
  if (length(missing_taxa) > 0) {
    cat("  Warning - missing taxa:", paste(missing_taxa, collapse = ", "), "\n")
  }
  
  # Reroot based on available taxa
  if (length(present_taxa) == 1) {
    # Root at single tip
    tip_node <- which(tree$tip.label == present_taxa)
    rooted_tree <- reroot(tree, node.number = tip_node)
    position <- tree$edge.length[tip_node]
    cat("  Rooting at single tip.\n")
    rooted_tree$edge.length[rooted_tree$edge == (length(tree$tip.label) + 1)] <- c(position / 2, position / 2)

  } else {
    # Root at MRCA of present taxa
    mrca_node <- getMRCA(tree, present_taxa)

    if (setequal(present_taxa, tree$tip.label)) {
      cat("  Special case: only target taxa present.\n")
    }

    if (mrca_node == (Ntip(tree) + 1)) {
      cat("  Already rooted at MRCA; reapplying rerooting for consistency.\n")
    } else {
      cat("  Rooting at MRCA node.\n")
    }

    rooted_tree <- reroot(tree, node.number = mrca_node, position = 0)
    position <- tree$edge.length[mrca_node]
    if (is.na(position)) position <- 0
    rooted_tree$edge.length[rooted_tree$edge == (length(tree$tip.label) + 1)] <- position / 3
  }

  # Clean edge lengths and ladderize for clarity
  rooted_tree$edge.length <- na.omit(rooted_tree$edge.length)
  rooted_tree <- ladderize(rooted_tree)

  # Clean up labels to prevent "1:" output
  rooted_tree$node.label <- NULL
  attr(rooted_tree, "node.label") <- NULL
  rooted_tree$tip.label <- as.character(rooted_tree$tip.label)

  # Save the rooted tree
  output_file <- sub("\\.treefile$", ".treefile.rooted", tree_file)
  write.tree(rooted_tree, file = output_file)
  
  cat("  Saved rooted tree to:", output_file, "\n\n")
}


# in some cases (~1000 genes) the rooting only works if you include salamanders in the taxa list
```
