## Appendices A1-A3 for "Whole-genome analyses point to new candidate genes underlying aestivation in amphibians": Appendix_A2_RER_analyses.R

```
# Whole-genome analyses point to new candidate genes underlying aestivation in amphibians
# RER analysis
# Ochoa-Ochoa & Tarvin (2025)
# RData available for this script in the same repository
# For questions and further assistance regarding this script:
# ---


# ====== Section 1: Load required libraries ======

library(devtools)
library(RColorBrewer)
library(gplots)
library(phytools)
library(maps)
library(Rcpp)
library(geiger)
library(knitr)
library(RcppArmadillo)
library(weights)
library(phangorn)
library(RERconverge)
library(dplyr)
library(tidyverse)
library(data.table)

#set your working directory
setwd("[set your working directory]")

# ====== Section 2: Prepare the data ======
#Make the two-column file for readTrees
files = list.files(pattern=".rooted") 
data_list <- lapply(files, function(file) {
  # Read the .txt file
  data <- read.table(file, header = F, sep = "\t") # Adjust header and separation as needed
  # Add a column with the filename
  data$file_name <- basename(file)
  return(data)
})
combined_data <- rbindlist(data_list, fill = TRUE, use.names = TRUE)

prueba<-combined_data %>% mutate(file_name = str_remove_all(file_name, ".mafft.aln.treefile.rooted"))
all_geneTrees<-prueba %>% relocate(file_name, .before = V1)

write.table(all_geneTrees,"[your working directory and file name]", sep="\t", row.names = F,
            col.names = F)

#Read all gene trees

protein_trees_xtro=readTrees("[your working directory and file name]", 
                             masterTree = NULL,  minTreesAll = 20, reestimateBranches = F, minSpecs = 5)

#Perform the main RER computation function

protRERw_xtro <- getAllResiduals(protein_trees_xtro,useSpecies= NULL,cutoff= 0.0001,
                                 transform = "sqrt", weighted = T, scale = T, plot = T) # the best selection, cutoff 0 or 0.0001, 


#Generate a phenotype paths vector from a phenotype tree
# Analyses binary trait evolution, presence-absence of aestivation with only the species from literature for presence
#trait_binary<-read.csv("[your working directory and file name]", header = F, row.names=1)

vec_binary <- setNames(trait_binary[[1]], rownames(trait_binary))


#Xenopus tropicalis
charpaths_xtro=char2Paths(vec_binary, protein_trees_xtro)

corEstBinary_xtro=correlateWithContinuousPhenotype(protRERw_xtro, charpaths_xtro, min.sp = 2, winsorizeRER = 3, winsorizetrait = 3)
write.csv(corEstBinary_xtro, file = "~/Desktop/Bioinformatic_paper/Results_June2025/RER_analyses/Xtro_results/corEstBinary_xtro_July2025.csv")
```
