## Appendices A1-A3 for "Whole-genome analyses point to new candidate genes underlying aestivation in amphibians": Appendix_A3_GO_analyses.R

```
# Whole-genome analyses point to new candidate genes underlying aestivation in amphibians
#Gene ontology analysis
# Ochoa-Ochoa & Tarvin (2025)
# RData available for this script in the same repository
# For questions and further assistance regarding this script:
# ---

# ====== Section 1: Activate and install necessary libraries ======
library(tidyverse)
library(pheatmap)
library(ggpubr)
library(ggplot2)
library(plotly)
library(ggVennDiagram)
library(stringr)
library(readr)
library(remotes)
#if (!require("BiocManager", quietly = TRUE))
#install.packages("BiocManager")
#BiocManager::install("topGO")
library(topGO)
#BiocManager::install("AnnotationDbi")
library(AnnotationDbi)
#install.packages("circlize")
#install.packages("reshape2")  # Si aún no lo tienes
library(dplyr)
library(circlize)
library(reshape2)

# ====== Section 2: Set working directory and load data ======
# Set working directory (update to your specific path)

setwd("[your working directory and file name]")

#Files from analysis 1) Correlate gene evolution with binary trait 
#evolution with aestivation data from literature

Gene_background<-read.csv("Xtro_Lit_background.csv")
Gene_negative<-read.csv("Xtro_Cont_negative.csv")
Gene_positive<-read.csv("Xtro_Cont_positive.csv")

Gene_background<- toupper(Gene_background$Gene)
Gene_negative<-toupper(Gene_negative$Gene)
Gene_positive<-toupper(Gene_positive$Gene)


### GO terms

# 1. Read your GO term file (e.g., from UniProt or eggNOG, Chicken GO terms)
# As specified in the manuscript for the analyses we use Xenopus tropicalis

ortho_go_raw_car <- read.delim(file.choose(), sep = "\t", header = TRUE)
ortho_go_raw_car$Gene.Names<-toupper(ortho_go_raw_car$Gene.Names)

# 2. Separate GO terms into rows and rename

gene2GO_car_t <- split(ortho_go_raw_car$Gene.Ontology.IDs, ortho_go_raw_car$Gene.Names)
gene2go <- lapply(gene2GO_car_t, function(go_terms) {
  go_terms %>%
    unlist() %>%
    strsplit(";\\s*") %>%   # split by "; " or ";"
    unlist() %>%
    unique() %>%
    .[. != ""]              # remove empty strings
})


## All the genes universe: 
# Your full universe

run_topGO <- function(gene_ids, background, onto, name) {
  geneList <- factor(as.integer(background %in% gene_ids))
  names(geneList) <- background
  
  GOdata <- new("topGOdata",
                ontology = onto,  # "MF"or "BP", "CC"
                allGenes = geneList,
                annot = annFUN.gene2GO,
                gene2GO = gene2go,
                nodeSize = 10)
  
  resultFisher <- runTest(GOdata, algorithm = "classic", statistic = "fisher")
  tab <- GenTable(GOdata, classicFisher = resultFisher, topNodes = 50)
  tab$group <- name
  tab$log10p <- -log10(as.numeric(tab$classicFisher))  # <- add this line
  
  return(tab)
}
# ====== Section 3: GO Analyses ======

## Only genes with significant evolutionary rate
## Be aware that the third element has to be specified for "MF"or "BP", "CC"

top_positive<- run_topGO(Gene_positive, Gene_background, "CC", "Positive")
top_negative<- run_topGO(Gene_negative, Gene_background, "CC", "Negative")

go_combined_up <- dplyr::bind_rows(top_positive, top_negative)
#write a file for each of the different category analyses
#write.csv(go_combined_up, file="Xtro_Cont_CC_top10.csv")


##### For getting the significant terms and the genes ########

geneList <- factor(as.integer(Gene_background %in% Gene_positive))
names(geneList) <- Gene_background

GOdata <- new("topGOdata",
              ontology = "CC",  # "MF"or "BP", "CC"
              allGenes = geneList,
              annot = annFUN.gene2GO,
              gene2GO = gene2go,
              nodeSize = 10)
resultFisher <- runTest(GOdata, algorithm = "classic", statistic = "fisher") 

sg <- sigGenes(GOdata)
str(sg)
numSigGenes(GOdata)
as.data.frame(sg)
#write.csv(sg, file="Sig_Genes_Lit_BP_negative10_xtro.csv")

# Table of top significant terms (optional: set topNodes or p-value cutoff)
topGO_results <- GenTable(GOdata,
                          classicFisher = resultFisher,
                          orderBy = "classicFisher",
                          topNodes = 50)  # Change to desired number of top GO terms
                          

# ---- EXTRACT GENES ----

# Get significant GO term IDs
sigTerms <- topGO_results$GO.ID

# Get all genes annotated to each GO term
go2genes <- genesInTerm(GOdata, sigTerms)

# ---- FORMAT AS DATA FRAME ----

# Create tidy data frame: GO_ID | Gene
go_gene_df <- bind_rows(lapply(names(go2genes), function(go_id) {
  tibble(GO_ID = go_id, Gene = go2genes[[go_id]])
}))

# Optionally join with GO term descriptions
go_gene_df <- left_join(go_gene_df, topGO_results[, c("GO.ID", "Term", "Significant", "classicFisher")],
                        by = c("GO_ID" = "GO.ID"))

write.csv(go_gene_df, file= "Lit_Xtr_CC_significant.csv")

# ====== Section 4: Graphs ======

# Prepare chord_df: select relevant columns
chord_df <- go_combined_up[, c("group", "GO.ID", "Significant")]

colnames(chord_df) <- c("group", "GO_term", "count")

# Prepare GO term info: merge GO.ID with description
go_info <- unique(go_combined_up[, c("GO.ID", "Term")])
colnames(go_info) <- c("GO_term", "Term")


# Merge GO descriptions into main data
chord_df_annot <- merge(chord_df, go_info, by = "GO_term")

# Add readable label: "GO:xxxxxxx - Description"
chord_df_annot$label <- paste0(chord_df_annot$GO_term, ":", chord_df_annot$Term)

# Select data for wide matrix format
chord_data <- chord_df_annot[, c("group", "label", "count")]

# Create group x GO label matrix
chord_matrix <- acast(chord_data, group ~ label, value.var = "count", fill = 0)

# Define custom colors for each group
grid.col <- c(
  "Positive" = "#ea7e24",
  "Negative"   = "#004483"
)

# Count sectors for proper gap setting
all_sectors <- unique(c(rownames(chord_matrix), colnames(chord_matrix)))
n_sectors <- length(all_sectors)

# Start PDF device
pdf("CC_Xtro_Cont.pdf", width = 10, height = 10)

# Clear any previous settings and define circular plot parameters
circos.clear()
par(mar = c(1, 1, 1, 1))  # Remove extra margins
circos.par(
  start.degree = 90,
  gap.after = rep(2, n_sectors),  # uniform gaps
  track.margin = c(0.01, 0.01),
  canvas.xlim = c(-1.5, 1.5),
  canvas.ylim = c(-1.5, 1.5)
)

# Draw the chord diagram
chordDiagram(
  chord_matrix,
  grid.col = grid.col,
  annotationTrack = "grid",
  preAllocateTracks = list(track.height = 0.08)
)

# Add rotated text labels (GO:description)
circos.trackPlotRegion(
  track.index = 1,
  panel.fun = function(x, y) {
    sector_name <- get.cell.meta.data("sector.index")
    xcenter <- get.cell.meta.data("xcenter")
    ycenter <- get.cell.meta.data("ycenter")
    circos.text(
      x = xcenter,
      y = ycenter,
      labels = sector_name,
      facing = "clockwise",
      niceFacing = TRUE,
      adj = c(0, 0.5),
      cex = 0.5  # adjust size if needed
    )
  },
  bg.border = NA
)

# Close the device
dev.off()

########### Venn Diagrams ###############
library(VennDiagram)
library(RColorBrewer)

All_genes<-read.csv("~/Desktop/Bioinformatic_paper/Results_June2025/Gene_ontology/binary.csv")
All_terms<-read.csv("~/Desktop/Bioinformatic_paper/Results_June2025/Gene_ontology/Terms.csv")

term_positive<-subset(All_genes, GeneSet == "pos")
term_negative<-subset(All_genes, GeneSet == "neg")

list_all <- list(
  Positive = term_positive$GO_ID,
  Negative = term_negative$GO_ID
)

ggVennDiagram(list_all, label_alpha = 1) +
  scale_fill_gradient(low = "white", high = "#ea7e24") +
  theme_void()

list_genes <- list(
  Positive = term_positive$Gene,
  Negative = term_negative$Gene
)

ggVennDiagram(list_genes, label_alpha = 1) +
  scale_fill_gradient(low = "white", high = "#ea7e24") +
  theme_void()

##########
MF<-subset(All_genes, Function == "MF")
BP<-subset(All_genes, Function == "BP")
CC<-subset(All_genes, Function == "CC")

Functions <- list(
  MF= MF$GO_ID,
  BP=BP$GO_ID,
  CC=CC$GO_ID)

ggVennDiagram(Functions, color = "black", lwd = 0.8, lty = 1) + 
  scale_fill_gradient(low = "#F4FAFE", high = "#4981BF")

Genes <- list(
  MF= MF$Gene,
  BP=BP$Gene,
  CC=CC$Gene)

ggVennDiagram(Genes, color = "black", lwd = 0.8, lty = 1) + 
  scale_fill_gradient(low = "#F4FAFE", high = "#4981BF")+
guides(fill = guide_legend(title = "Significant Genes")) +
  theme(legend.title = element_text(color = "black"),
        legend.position = "bottom")
```
