## Appendix A4 for "Whole-genome analyses point to new candidate genes underlying aestivation in amphibians"

### Supplementary material

### Additional methods and results

**Figure S1.** Number of genes shared among the different categories of gene term ontology: Molecular Function (MF), Biological Process (BP) and Cellular Component (CC).

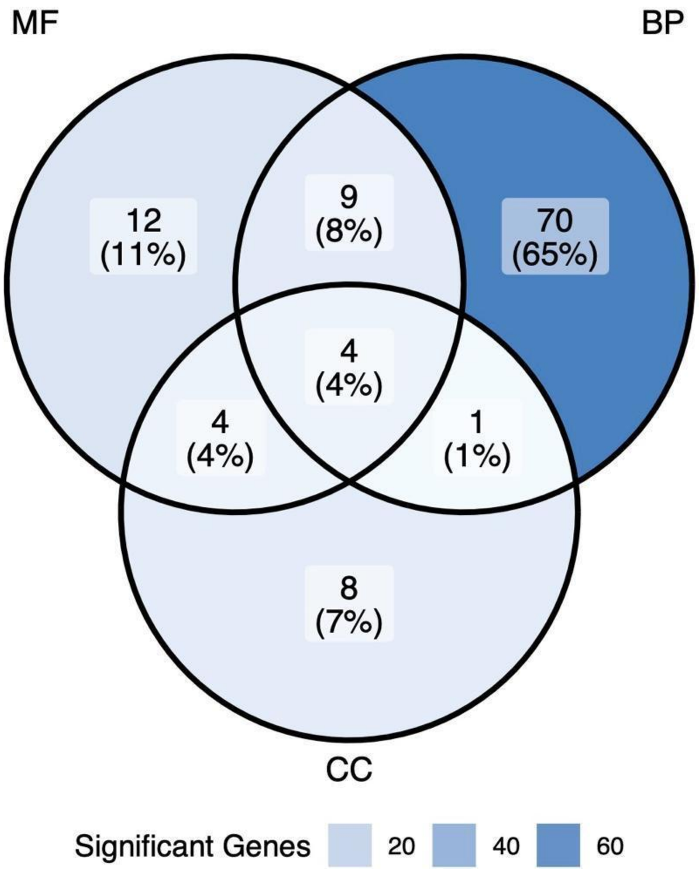

**Figure S2.** Molecular function GO terms that were significantly enriched in the set of RER genes with significantly evolutionary rates. See supplementary table S5 for the parent-children term relationship.

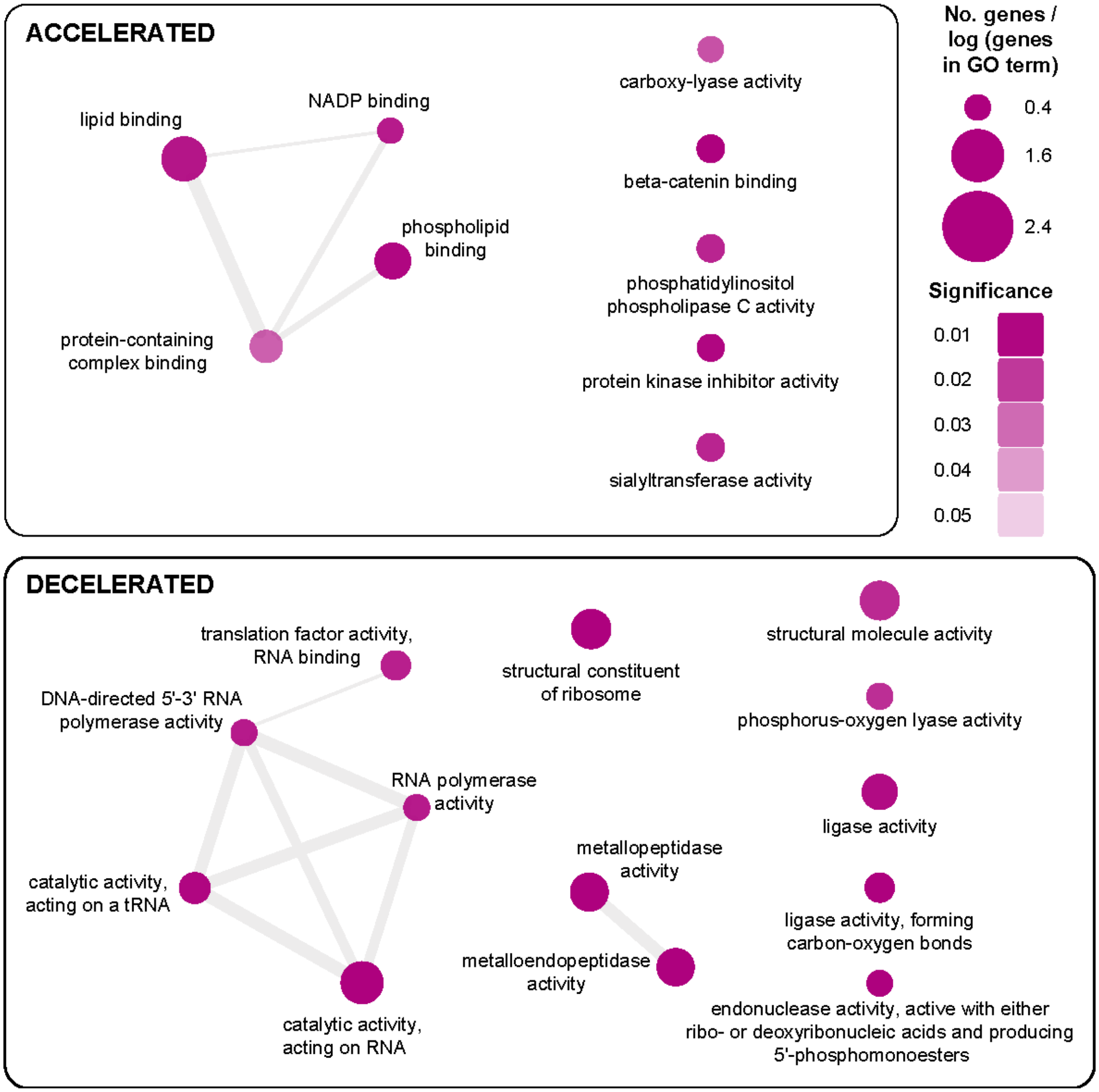

**Figure S3.** Biological Process GO terms that were significantly enriched in the set of RER genes with significantly evolutionary rates. See supplementary table S5 for the parent-children term relationship.

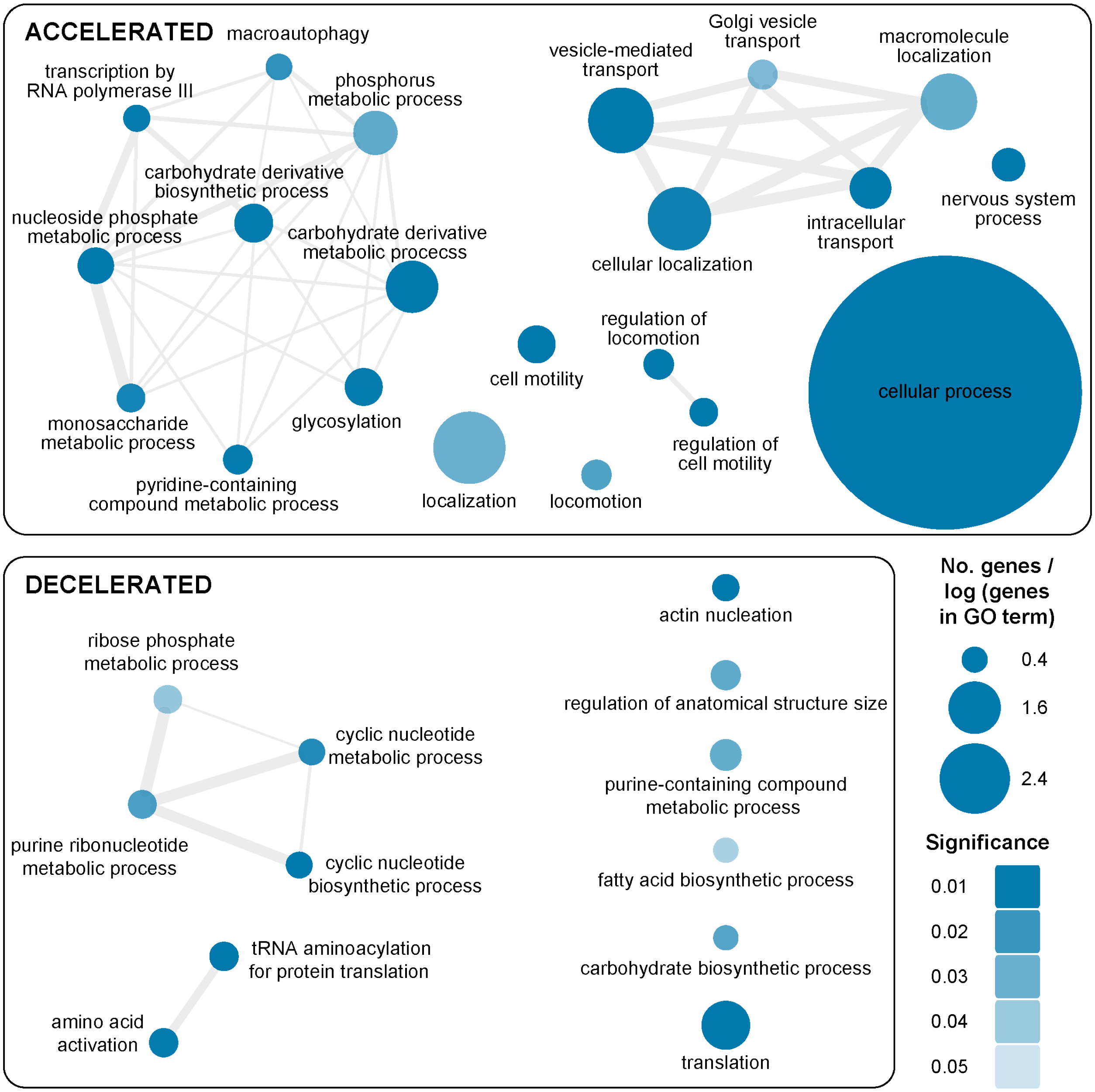

**Figure S4.** Cellular Component GO terms that were significantly enriched in the set of RER genes with significantly evolutionary rates. See supplementary table S5 for the parent-children term relationship.

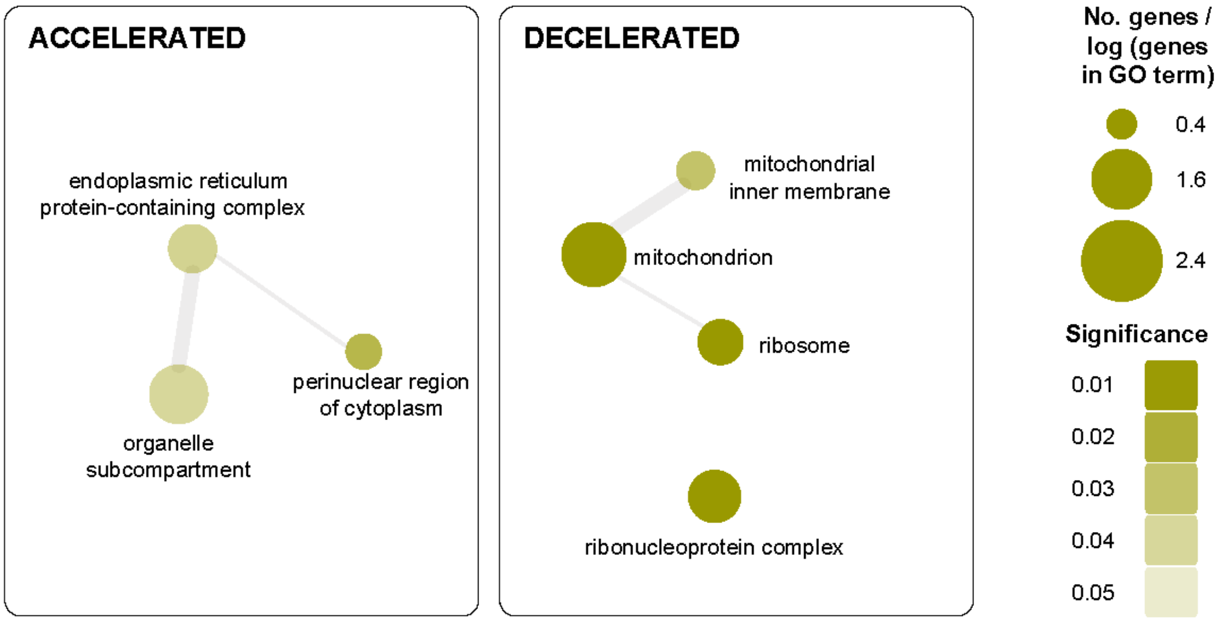


**Additional Results**

**Description of the genes with top relative evolutionary rates (RERs) that were also the focus of prior studies identified in our literature review (Table 4). All gene descriptions were copied word-for-word from UniProt database (The UniProt Consortium 2025) under a CC BY 4.0 license [https://www.uniprot.org/](https://www.uniprot.org/) (accessed Dec 12, 2025). When descriptions from *Xenopus tropicalis* were not available we searched for those from *Homo sapiens*.**

### Accelerated & Increased

Accelerated and increased genes mean those that are changing rapidly and becoming different from the ancestral form and more diverse among lineages;  and that in experiments showed an increased activity in at least one tissue. Four genes (*DLG1, EIF4ENIF1, GSPT1*, and *HK1*) fall within this category, all from the skeletal muscle of aestivating *Cyclorana alboguttata* (Hudson et al. 2024).

***DLG1.*** "Essential multidomain scaffolding protein required for normal development (By similarity). Recruits channels, receptors and signaling molecules to discrete plasma membrane domains in polarized cells. Promotes epithelial cell layer barrier function via maintaining cell-cell adhesion (By similarity). May also play a role in adherens junction assembly, signal transduction, cell proliferation, synaptogenesis and lymphocyte activation. Regulates the excitability of cardiac myocytes by modulating the functional expression of Kv4 channels. Functional regulator of Kv1.5 channel. During long-term depression in hippocampal neurons, it recruits ADAM10 to the plasma membrane (PubMed:23676497)."

***EIF4ENIF1.*** "EIF4E-binding protein that regulates translation and stability of mRNAs in processing bodies (P-bodies) (PubMed:16157702, PubMed:24335285, PubMed:27342281, PubMed:32354837). Plays a key role in P-bodies to coordinate the storage of translationally inactive mRNAs in the cytoplasm and prevent their degradation (PubMed:24335285, PubMed:32354837). Acts as a binding platform for multiple RNA-binding proteins: promotes deadenylation of mRNAs via its interaction with the CCR4-NOT complex, and blocks decapping via interaction with eIF4E (EIF4E and EIF4E2), thereby protecting deadenylated and repressed mRNAs from degradation (PubMed:27342281, PubMed:32354837). Component of a multiprotein complex that sequesters and represses translation of proneurogenic factors during neurogenesis (By similarity). Promotes miRNA-mediated translational repression (PubMed:24335285, PubMed:27342281, PubMed:28487484). Required for the formation of P-bodies (PubMed:16157702, PubMed:22966201, PubMed:27342281, PubMed:32354837). Involved in mRNA translational repression mediated by the miRNA effector TNRC6B by protecting TNRC6B-targeted mRNAs from decapping and subsequent decay (PubMed:32354837). Also acts as a nucleoplasmic shuttling protein, which mediates the nuclear import of EIF4E and DDX6 by a piggy-back mechanism (PubMed:10856257, PubMed:28216671)."

***GSPT1.*** "GTPase component of the eRF1-eRF3-GTP ternary complex, a ternary complex that mediates translation termination in response to the termination codons UAA, UAG and UGA (PubMed:15987998, PubMed:19417105, PubMed:2511002, PubMed:27863242). GSPT1/ERF3A mediates ETF1/ERF1 delivery to stop codons: The eRF1-eRF3-GTP complex binds to a stop codon in the ribosomal A-site (PubMed:27863242). GTP hydrolysis by GSPT1/ERF3A induces a conformational change that leads to its dissociation, permitting ETF1/ERF1 to accommodate fully in the A-site (PubMed:16777602, PubMed:27863242). Component of the transient SURF complex which recruits UPF1 to stalled ribosomes in the context of nonsense-mediated decay (NMD) of mRNAs containing premature stop codons (PubMed:24486019). Required for SHFL-mediated translation termination which inhibits programmed ribosomal frameshifting (-1PRF) of mRNA from viruses and cellular genes (PubMed:30682371)."

***HK1.*** "Catalyzes the phosphorylation of various hexoses, such as D-glucose, D-glucosamine, D-fructose, D-mannose and 2-deoxy-D-glucose, to hexose 6-phosphate (D-glucose 6-phosphate, D-glucosamine 6-phosphate, D-fructose 6-phosphate, D-mannose 6-phosphate and 2-deoxy-D-glucose 6-phosphate, respectively) (PubMed:1637300, PubMed:25316723, PubMed:27374331). Does not phosphorylate N-acetyl-D-glucosamine (PubMed:27374331). Mediates the initial step of glycolysis by catalyzing phosphorylation of D-glucose to D-glucose 6-phosphate (By similarity). Involved in innate immunity and inflammation by acting as a pattern recognition receptor for bacterial peptidoglycan (PubMed:27374331). When released in the cytosol, N-acetyl-D-glucosamine component of bacterial peptidoglycan inhibits the hexokinase activity of HK1 and causes its dissociation from mitochondrial outer membrane, thereby activating the NLRP3 inflammasome (PubMed:27374331)."

### Accelerated & Decreased

Accelerated and decreased genes mean those that are changing rapidly and becoming different from the ancestral form and more diverse among lineages; and that in experiments showed a decreased activity (downregulated) in at least one tissue. Seven genes (*AUTS2, BFAR, CDKN1B, HHEX, NOS3, SAP30*, and *YBX1*) fall within this category, all from the skeletal muscle of aestivating *Cyclorana alboguttata*, most of them reported in Hudson et al. (2024), only *CDKN1B* and *NOS3* from Reilly et al. (2013).

***AUTS2.*** "Component of a Polycomb group (PcG) multiprotein PRC1-like complex, a complex class required to maintain the transcriptionally repressive state of many genes, including Hox genes, throughout development. PcG PRC1 complex acts via chromatin remodeling and modification of histones; it mediates monoubiquitination of histone H2A 'Lys-119', rendering chromatin heritably changed in its expressibility (PubMed:25519132). The PRC1-like complex that contains *PCGF5, RNF2, CSNK2B, RYBP* and *AUTS2* has decreased histone H2A ubiquitination activity, due to the phosphorylation of RNF2 by *CSNK2B* (PubMed:25519132). As a consequence, the complex mediates transcriptional activation (PubMed:25519132). In the cytoplasm, plays a role in axon and dendrite elongation and in neuronal migration during embryonic brain development. Promotes reorganization of the actin cytoskeleton, lamellipodia formation and neurite elongation via its interaction with RAC guanine nucleotide exchange factors, which then leads to the activation of *RAC1* (By similarity)."

***BFAR.*** "Membrane-bound E3 ubiquitin ligase that plays a role in several processes including apoptosis regulation or reticulum endoplasmic stress. Has anti-apoptotic activity, both for apoptosis triggered via death-receptors and via mitochondrial factors. Contributes to the dynamic control of IRE1/ERN1 signaling during ER stress by inducing BAX inhibitor 1/TMBIM6 proteasomal degradation. Promotes the activation of TGF-beta signaling by mediating the 'Lys-

63'-linked ubiquitination of TGFBR1 which is critical to activate the pathway. Together with NGFR, negatively regulates NF-kappa-B and JNK-related signaling pathways. Promotes the proteasome-mediated degradation of PNPLA3, a protein involved in lipid metabolism."

***CDKN1B.*** "Transcription factor that controls hypaxial muscle development by down-regulating myod1 and cdkn1b/p27, thereby allowing myoblasts to proliferate before the onset of terminal differentiation. Important regulator of cell cycle progression. Inhibits the kinase activity of CDK2 bound to cyclin A, but has little inhibitory activity on CDK2 bound to SPDYA (PubMed:28666995). Involved in G1 arrest. Potent inhibitor of cyclin E- and cyclin A-CDK2 complexes. Forms a complex with cyclin type D-CDK4 complexes and is involved in the assembly, stability, and modulation of CCND1-CDK4 complex activation. Acts either as an inhibitor or an activator of cyclin type D-CDK4 complexes depending on its phosphorylation state and/or stoichometry."

***HHEX.*** "Recognizes the DNA sequence 5'-ATTAA-3'. Transcriptional repressor. Regulates the differentiation of both endothelial and blood cells. Probably plays a role in the proliferation of vascular endothelial cells during blood vessel development. Establishes anterior identity at two levels; acts early to enhance canonical wnt-signaling by repressing expression of tle4, and acts later to inhibit nodal-signaling by directly targeting nodal/nr1 and nodal2/nr2. May play a role in liver development. Induces heart development (By similarity)."

***NOS3.*** "Produces nitric oxide (NO) which is a messenger molecule with diverse functions."

***SAP30.*** "Functions as a transcription repressor, probably via its interaction with histone deacetylase complexes. Involved in the functional recruitment of the class 1 Sin3-histone deacetylase complex (HDAC) to the nucleolus. Binds DNA, apparently without sequence-specificity, and bends bound double-stranded DNA. Binds phosphoinositol phosphates (phosphoinositol 3-phosphate, phosphoinositol 4-phosphate and phosphoinositol 5-phosphate) via the same basic sequence motif that mediates DNA binding and nuclear import."

***YBX1.*** "DNA- and RNA-binding protein involved in various processes, such as translational repression, RNA stabilization, mRNA splicing and transcription regulation (By similarity). Binds preferentially to the 5'-[CU]CUGCG-3' RNA motif and specifically recognizes mRNA transcripts modified by C5-methylcytosine (m5C) (By similarity).
Promotes mRNA stabilization: acts by binding to m5C-containing mRNAs and preventing mRNA decay (By similarity). Plays a role in the maternal-to-zygotic transition in early embryo by binding to m5C-containing maternal mRNAs and preventing their degradation (By similarity).
Also promotes maternal-to-zygotic transition in oocytes and embryos by promoting translation repression; molecular mechanisms governing translation repression are unknown (By similarity).

Plays a key role in RNA composition of extracellular exosomes by defining the sorting of small non-coding RNAs, such as tRNAs, Y RNAs, Vault RNAs and miRNAs (By similarity). Probably sorts RNAs in exosomes by recognizing and binding C5-methylcytosine (m5C)-containing RNAs (By similarity). Acts as a key effector of epidermal progenitors by preventing epidermal progenitor senescence: acts by regulating the translation of a senescence-associated subset of cytokine mRNAs, possibly by binding to m5C-containing mRNAs (By similarity). Also involved in pre-mRNA alternative splicing regulation: binds to splice sites in pre-mRNA and regulates splice site selection (By similarity). Also able to bind DNA and regulate transcription (PubMed:2247479). Binds to promoters that contain a Y-box (5'-CTGATTGGCCAA-3') (PubMed:2247479). Promotes separation of DNA strands that contain mismatches or are modified by cisplatin (By similarity). Has endonucleolytic activity and can introduce nicks or breaks into double-stranded DNA, suggesting a role in DNA repair (By similarity).
The secreted form acts as an extracellular mitogen and stimulates cell migration and proliferation (By similarity)."

### Decelerated & Increased

Decelerated and increased genes mean those that are conserved with reference from the ancestral form and are very similar among lineages; and that in experiments showed an increased activity (upregulated) in at least one tissue. Eight genes (*ANP32B, GPI, JCHAIN, PKM, POLR2B, SARS1, UBE2G1,* and *UBE2V2*) fall within this category, all from the skeletal muscle of aestivating *Cyclorana alboguttata*, most of them reported in Hudson et al. (2024), *SARS1* only reported in Reilly et al. (2013). *GPI* was also reported in Reilly et al. (2013) and for dehydrated *Xenopus laevis* (Wu et al., 2017). *PKM* has been reported in various articles, Wu et al. (2017) found an increased activity in brain and kidney but decreased in heart (Table 4, and S1). In other studies, a decreased activity was found in skeletal muscle of aestivating frogs (*P. cristiceps*, Moreira et al. 2020; *C. alboguttata*, Reilly et al. 2013; Hudson et al. 2024), and a maintained activity in *P. diplolister* (Moreira et al. 2021; Table S1). Finally, *JCHAIN* also have a decreased activity in skeletal muscle of aestivating *Cyclorana alboguttata* (Reilly et al. 2013).

***ANP32B.*** "Multifunctional protein working as a cell cycle progression factor as well as a cell survival factor. Required for the progression from the G1 to the S phase. Anti-apoptotic protein which functions as a caspase-3 inhibitor. Has no phosphatase 2A (PP2A) inhibitor activity. Exhibits histone chaperone properties, stimulating core histones to assemble into a nucleosome (By similarity)."

***GPI.*** "In the cytoplasm, catalyzes the conversion of glucose-6-phosphate to fructose-6-phosphate, the second step in glycolysis, and the reverse reaction during gluconeogenesis. Besides its role as a glycolytic enzyme, also acts as a secreted cytokine: acts as an angiogenic factor (AMF) that stimulates endothelial cell motility. Acts as a neurotrophic factor, neuroleukin, for spinal and sensory neurons. It is secreted by lectin-stimulated T-cells and induces immunoglobulin secretion."

***JCHAIN.*** "Serves to link two monomer units of either IgM or IgA. In the case of IgM, the J chain-joined dimer is a nucleating unit for the IgM pentamer, and in the case of IgA it induces dimers and/or larger polymers. It also helps to bind these immunoglobulins to secretory component."

***PKM.*** "Catalyzes the final rate-limiting step of glycolysis by mediating the transfer of a phosphoryl group from phosphoenolpyruvate (PEP) to ADP, generating ATP (PubMed:15996096, PubMed:1854723, PubMed:20847263).
The ratio between the highly active tetrameric form and nearly inactive dimeric form determines whether glucose carbons are channeled to biosynthetic processes or used for glycolytic ATP production (PubMed:15996096, PubMed:1854723, PubMed:20847263).
The transition between the 2 forms contributes to the control of glycolysis and is important for tumor cell proliferation and survival (PubMed:15996096, PubMed:1854723, PubMed:20847263)."

***POLR2B.*** "DNA-dependent RNA polymerase catalyzes the transcription of DNA into RNA using the four ribonucleoside triphosphates as substrates. RNA-dependent RNA polymerase that catalyzes the extension of a non-coding RNA (ncRNA) at the 3'-end using the four ribonucleoside triphosphates as substrates. An internal ncRNA sequence near the 3'-end serves as a template in a single-round Pol II-mediated RNA polymerization reaction. May decrease the stability of ncRNAs that repress Pol II-mediated gene transcription. Pol II-mediated transcription cycle proceeds through transcription initiation, transcription elongation and transcription termination stages. During transcription initiation, Pol II pre-initiation complex (PIC) is recruited to DNA promoters, with focused-type promoters containing either the initiator (Inr) element, or the TATA-box found in cell-type specific genes and dispersed-type promoters that often contain hypomethylated CpG islands usually found in housekeeping genes. Once the polymerase has escaped from the promoter it enters the elongation phase during which RNA is actively polymerized, based on complementarity with the template DNA strand. Transcription termination involves the release of the RNA transcript and polymerase from the DNA (PubMed:27193682, PubMed:30190596, PubMed:9852112).
Forms Pol II active center together with the largest subunit POLR2A/RPB1. Appends one nucleotide at a time to the 3' end of the nascent RNA, with POLR2A/RPB1 most likely contributing a Mg2+-coordinating DxDGD motif and POLR2B/RPB2 participating in the coordination of a second Mg2+ ion and providing lysine residues believed to facilitate Watson-Crick base pairing between the incoming nucleotide and template base. Typically, Mg2+ ions direct a 5' nucleoside triphosphate to form a phosphodiester bond with the 3' hydroxyl of the

preceding nucleotide of the nascent RNA, with the elimination of pyrophosphate. The reversible pyrophosphorolysis can occur at high pyrophosphate concentrations (By similarity) (PubMed:30190596, PubMed:9852112). Can proofread the nascent RNA transcript by means of a 3' -> 5' exonuclease activity. If a ribonucleotide is mis-incorporated, backtracks along the template DNA and cleaves the phosphodiester bond releasing the mis-incorporated 5'-ribonucleotide (By similarity) (PubMed:8381534). RNA-dependent RNA polymerase that catalyzes the extension of a non-coding RNA (ncRNA) at the 3'-end using the four ribonucleoside triphosphates as substrates. An internal ncRNA sequence near the 3'-end serves as a template in a single-round Pol II-mediated RNA polymerization reaction. May decrease the stability of ncRNAs that repress Pol II-mediated gene transcription."

***SARS1***. "Catalyzes the attachment of serine to tRNA(Ser) in a two-step reaction: serine is first activated by ATP to form Ser-AMP and then transferred to the acceptor end of tRNA(Ser) (PubMed:22353712, PubMed:24095058, PubMed:26433229, PubMed:28236339, PubMed:34570399, PubMed:36041817, PubMed:9431993). Is probably also able to aminoacylate tRNA(Sec) with serine, to form the misacylated tRNA L-seryl-tRNA(Sec), which will be further converted into selenocysteinyl-tRNA(Sec) (PubMed:26433229, PubMed:28236339, PubMed:34570399, PubMed:9431993). In the nucleus, binds to the VEGFA core promoter and prevents MYC binding and transcriptional activation by MYC (PubMed:24940000). Recruits SIRT2 to the VEGFA promoter, promoting deacetylation of histone H4 at 'Lys-16' (H4K16). Thereby, inhibits the production of VEGFA and sprouting angiogenesis mediated by VEGFA (PubMed:19423847, PubMed:19423848, PubMed:24940000)."

***UBE2G1*. "**Accepts ubiquitin from the E1 complex and catalyzes its covalent attachment to other proteins. In vitro catalyzes 'Lys-48'-, as well as 'Lys-63'-linked polyubiquitination. May be involved in degradation of muscle-specific proteins. Mediates polyubiquitination of CYP3A4."

***UBE2V2*.** "Has no ubiquitin ligase activity on its own. The UBE2V2/UBE2N heterodimer catalyzes the synthesis of non-canonical poly-ubiquitin chains that are linked through 'Lys-63'. This type of poly-ubiquitination does not lead to protein degradation by the proteasome. Mediates transcriptional activation of target genes. Plays a role in the control of progress through the cell cycle and differentiation. Plays a role in the error-free DNA repair pathway and contributes to the survival of cells after DNA damage."

### Decelerated & Decreased

Decelerated and decreased genes mean those that are conserved with reference from the ancestral form and are very similar among lineages; and that in experiments showed a decreased activity (downregulated) in at least one tissue. Thirteen genes (*ABRACL, ACE, ADAM23, ANXA4,*

*ATP5PB, DDIT4, GPI, JCHAIN, KIAA1191, MLYCD, MMEL1, NDUFB3,* and *NDUFS4*) fall within this category, all from the skeletal muscle of aestivating *Cyclorana alboguttata*, all of them reported in Hudson et al. (2024) and *DDIT4, GPI,* and *JCHAIN* reported in Reilly et al. (2013).

***ABRACL.*** No info

***ACE.*** "Catalysis of the hydrolysis of peptide bonds by a mechanism in which water acts as a nucleophile, one or two metal ions hold the water molecule in place, and charged amino acid side chains are ligands for the metal ions."

***ADAM23.*** "May play a role in cell-cell and cell-matrix interactions. This is a non-catalytic metalloprotease-like protein."

***ANXA4.*** "Calcium/phospholipid-binding protein which promotes membrane fusion and is involved in exocytosis."

***ATP5PB.*** "Subunit b, of the mitochondrial membrane ATP synthase complex ($F_1F_0$ ATP synthase or Complex V) that produces ATP from ADP in the presence of a proton gradient across the membrane which is generated by electron transport complexes of the respiratory chain. ATP synthase complex consist of a soluble $F_1$ head domain - the catalytic core - and a membrane $F_1$ domain - the membrane proton channel. These two domains are linked by a central stalk rotating inside the $F_1$ region and a stationary peripheral stalk. During catalysis, ATP synthesis in the catalytic domain of $F_1$ is coupled via a rotary mechanism of the central stalk subunits to proton translocation. In vivo, can only synthesize ATP although its ATP hydrolase activity can be activated artificially in vitro. Part of the complex $F_0$ domain. Part of the complex $F_0$ domain and the peripheric stalk, which acts as a stator to hold the catalytic $alpha_3beta_3$ subcomplex and subunit a/ATP6 static relative to the rotary elements."

***DDIT4.*** "Regulates cell growth, proliferation and survival via inhibition of the activity of the mammalian target of rapamycin complex 1 (mTORC1). Inhibition of mTORC1 is mediated by a pathway that involves ddit4/redd1, akt1, the tsc1-tsc2 complex and the GTPase rheb. Plays an important role in responses to cellular energy levels and cellular stress, including responses to hypoxia and DNA damage, via its effect on mTORC1 activity. Plays a role in neuronal differentiation, neuron migration during embryonic brain development and in neuronal cell death (By similarity)."

***GPI.*** "In the cytoplasm, catalyzes the conversion of glucose-6-phosphate to fructose-6-phosphate, the second step in glycolysis, and the reverse reaction during gluconeogenesis. Besides it's role as a glycolytic enzyme, also acts as a secreted cytokine: acts as an angiogenic factor (AMF) that stimulates endothelial cell motility. Acts as a neurotrophic factor, neuroleukin, for spinal and sensory neurons. It is secreted by lectin-stimulated T-cells and induces immunoglobulin secretion."

***JCHAIN.*** "Serves to link two monomer units of either IgM or IgA. In the case of IgM, the J chain-joined dimer is a nucleating unit for the IgM pentamer, and in the case of IgA it induces dimers and/or larger polymers. It also helps to bind these immunoglobulins to secretory component."

***KIAA1191.*** "Potential NADPH-dependent oxidoreductase. May be involved in the regulation of neuronal survival, differentiation and axonal outgrowth."

***MLYCD.*** "Catalyzes the conversion of malonyl-CoA to acetyl-CoA. In the fatty acid biosynthesis MCD selectively removes malonyl-CoA and thus assures that methyl-malonyl-CoA is the only chain elongating substrate for fatty acid synthase and that fatty acids with multiple methyl side chains are produced. In peroxisomes it may be involved in degrading intraperoxisomal malonyl-CoA, which is generated by the peroxisomal beta-oxidation of odd chain-length dicarboxylic fatty acids. Plays a role in the metabolic balance between glucose and lipid oxidation in muscle independent of alterations in insulin signaling. May play a role in controlling the extent of ischemic injury by promoting glucose oxidation."

***MMEL1.*** "Metalloprotease involved in sperm function, possibly by modulating the processes of fertilization and early embryonic development. Degrades a broad variety of small peptides with a preference for peptides shorter than 3 kDa containing neutral bulky aliphatic or aromatic amino acid residues. Shares the same substrate specificity with MME and cleaves peptides at the same amide bond (By similarity)."

***NDUFB3.*** "Accessory subunit of the mitochondrial membrane respiratory chain NADH dehydrogenase (Complex I), that is believed not to be involved in catalysis. Complex I functions in the transfer of electrons from NADH to the respiratory chain. The immediate electron acceptor for the enzyme is believed to be ubiquinone."

***NDUFS4.*** "Accessory subunit of the mitochondrial membrane respiratory chain NADH dehydrogenase (Complex I), that is believed not to be involved in catalysis. Complex I functions in the transfer of electrons from NADH to the respiratory chain. The immediate electron acceptor for the enzyme is believed to be ubiquinone."

***PKM.*** "Catalyzes the final rate-limiting step of glycolysis by mediating the transfer of a phosphoryl group from phosphoenolpyruvate (PEP) to ADP, generating ATP (PubMed:15996096, PubMed:1854723, PubMed:20847263).

The ratio between the highly active tetrameric form and nearly inactive dimeric form determines whether glucose carbons are channeled to biosynthetic processes or used for glycolytic ATP production (PubMed:15996096, PubMed:1854723, PubMed:20847263).
The transition between the 2 forms contributes to the control of glycolysis and is important for tumor cell proliferation and survival (PubMed:15996096, PubMed:1854723, PubMed:20847263)."

**Literature Cited**


Hudson N. J., T. G. A. Lonhienne, C. E. Franklin, G. S. Harper, and S. A. Lehnert. 2008. Epigenetic silencers are enriched in dormant desert frog muscle. J Comp Physiol B 178:729–734. doi:10.1007/s00360-008-0261-0.

Reilly B.D., D.I. Schlipalius, R.L. Cramp, P.R. Ebert, and C.E. Franklin. 2013. Frogs and estivation: transcriptional insights into metabolism and cell survival in a natural model of extended muscle disuse. Physiol Genomics 45:377–388. doi:10.1152/physiolgenomics.00163.2012.

The UniProt Consortium. 2025. UniProt: the universal protein knowledgebase in 2025. Nucleic Acids Res 53:D609–D617. doi:10.1093/nar/gkae1010.
